# The genetic architecture of human DNA methylome variation

**DOI:** 10.64898/2026.11.01.763325

**Authors:** Tianjie Liu, Juan F. Macias-Velasco, Xiaoyu Zhuo, Jiadong Lin, David Porubsky, Wenjin Zhang, Daofeng Li, Chad Tomlinson, Edward A. Belter, Zheng Dong, Shihua Dong, Human Pangenome Reference Consortium (HPRC), Evan E. Eichler, Ting Wang

## Abstract

Population DNA methylome analyses remain largely confined to reference-defined genomic space despite extensive human genetic diversity. A central challenge is therefore to define comparable methylation phenotypes across diverse genomes and map the genetic architecture of methylome variation. Here, using 460 haplotypes from the Human Pangenome Reference Consortium Release 2, we develop two complementary pangenome-based frameworks: one compares methylation within homologous regions while retaining haplotype-specific structural configuration and CpG content; the other represents CpG island (CGI) variation at homologous loci as sequence-resolved alleles. We identified 47,039 CGI alleles, including 8,040 absent from CHM13; 62% were attributable to structural variants and 31% to small variants. Through local genealogies, we reconstructed CGI gain, loss and turnover, revealing how present-day methylation differences align with CGI evolutionary trajectories. We found that genome-wide, methylation variability was buffered against genetic diversity across more than 90% of the genome, with exceptions at extreme nucleotide diversity and locally around structural variants. Our methylation framework enabled population-scale allele-aware methylation quantitative trait locus (mQTL) mapping into non-reference and structurally complex sequence, where mQTL-associated intervals were enriched. mQTLs linked methylation to transcriptomic phenotypes, population differentiation and complex traits, supporting 8% of population differentially methylated regions and colocalizing with more than half of genome-wide association study loci. Together, our study redefines human methylome analysis and our findings reveal a genome-wide architecture of genetic-epigenetic coupling, reshaping how methylation variation is interpreted across human development and disease.

## Main

In mammalian somatic cells, 5-methylcytosine occurs predominantly at CpG dinucleotides. CpG islands (CGIs) were originally identified as CpG-rich, unmethylated regions^1^ and are now annotated from sequence composition^2,3^. CGIs overlap many promoters and regulatory elements, and their gain and loss across species have been implicated in regulatory evolution^4,5^. CpG sites and CGIs are sequence-defined, but their presence is not fixed across genomes. Nonetheless, most human methylomes are analyzed using CpGs defined on a single reference genome^6–9^. Reference-based methylome comparisons can therefore conflate variation in the underlying genomic sequence with variation in methylation state. How human methylome variation should be defined and compared when CpG content and CGI landscape are themselves polymorphic remains poorly resolved.

Genetic variation is a major determinant of inter-individual variation in DNA methylation^7^. Studies of methylation quantitative trait loci (mQTLs) have established widespread cis genetic effects and connected methylation-associated variants to molecular phenotypes and complex traits^8,10–14^. Variants act through at least four routes: they create or remove a CpG^15^, alter a transcription-factor binding motif^9,16,17^, change local chromatin^16,18,19^, or add, remove or reposition sequence through structural variation^20,21^. Genetic control of methylation therefore spans scales, from individual CpGs and local regulatory sequences to large structural variations. At the population level, these genetic effects can translate differences in allele frequency into differences in methylation, even when the same allele has the same effect across ancestries^22,23^. Resolving the genetic architecture of methylation variation across diverse genomes remains challenging because genetic divergence from a single reference biases functional-genomic measurements, particularly in structurally variable and difficult-to-map regions^24,25^. Long-read sequencing and haplotype-resolved assemblies now recover sequence and methylation from the same molecules, extending analysis into repeats, structural variants (SVs) and non-reference sequence^26–28^. The Human Pangenome Reference Consortium Release 2 (HPRC2) panepigenome identified 17.6 million CpGs absent from T2T-CHM13 across 460 haplotypes, a 51.9% expansion over the reference^29^. Personal-assembly and graph-based approaches therefore expand the genomic space that can be measured^30^, but do not by themselves solve the central problem of defining comparable methylation phenotypes across homologous regions that differ in length, orientation, structural configuration, CpG content or CGI presence.

Here we combine 460 haplotype-resolved assemblies from 230 individuals across five superpopulations with matched long-read methylomes and 202 matched long-read transcriptomes, all from lymphoblastoid cell lines (Supplementary Table 1)^29,31^. We establish two complementary frameworks. The first establishes a common coordinate system to directly compare methylation across homologous regions in genetically diverse haplotypes. The second, PanCGI, defines CGIs independently for each assembly and integrates them into a sequence-resolved catalogue of CGI alleles at each homologous locus. These new frameworks enable us to elucidate the genetic architecture of human DNA methylome variation.

### Pangenome mapping jointly models variation in CpG content and DNA methylation

Homologous regions can differ among haplotypes in length, orientation, repeat content and CpG number. To implement the first framework, we partitioned T2T-CHM13 into non-overlapping 200-bp intervals and projected each interval through the pangenome graph onto each phased assembly. The sequence recovered on each haplotype defined its haplotype-specific interval, in which we measured CpG content and methylation (**Fig. 1a**). At a chromosome 8 locus, for example, one haplotype contained a 16-bp insertion and a 332-bp Alu insertion, a second contained both insertions within a large inversion, and a third contained an 88-bp deletion within the same inverted region. Methylation was then compared across homologous haplotype-specific intervals despite these distinct structural configurations (**Fig. 1b**).

**Fig. 1.**
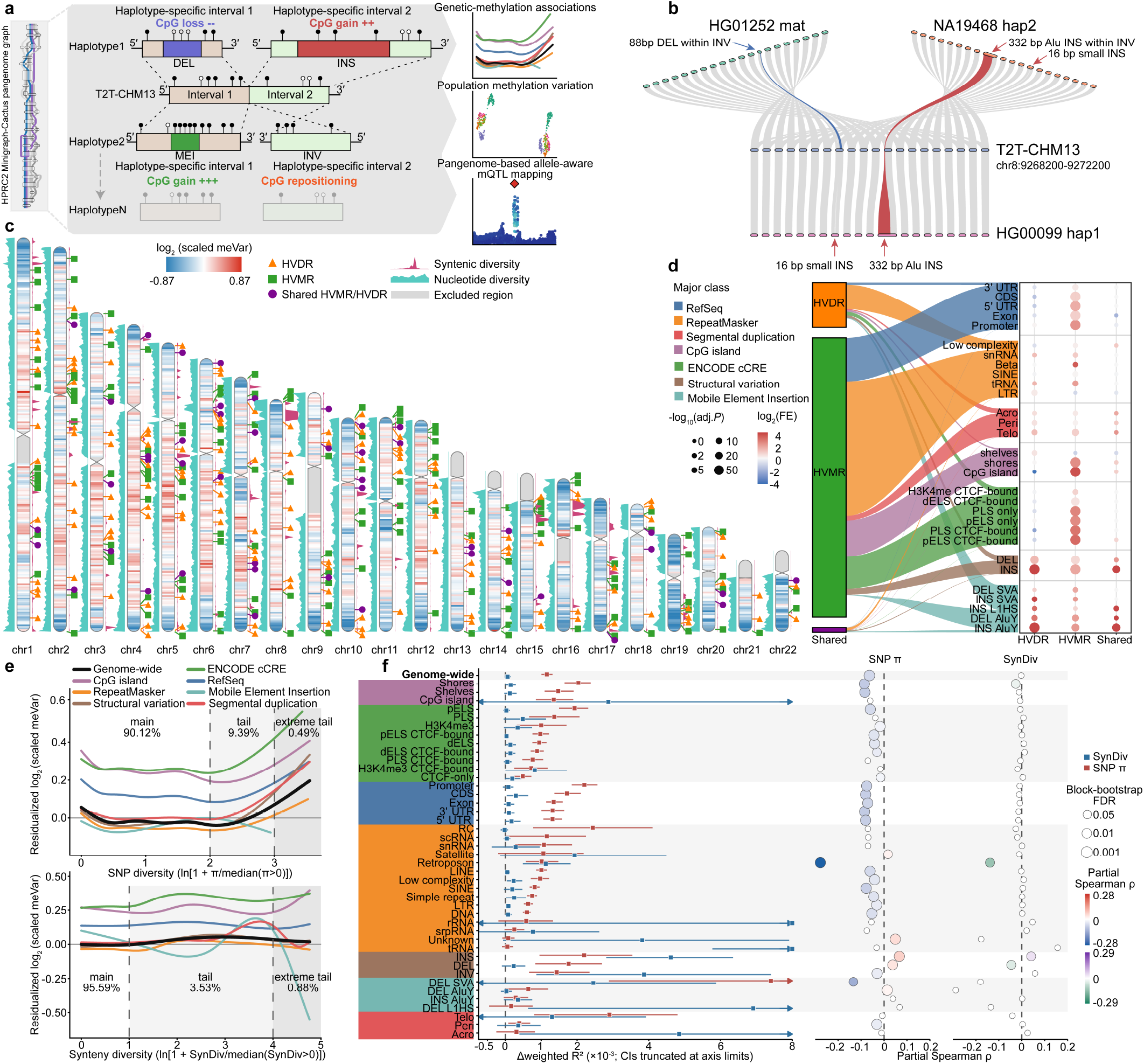
Pangenome-resolved variation in CpG content and DNA methylation. a, Non-overlapping 200-bp T2T-CHM13 intervals were projected onto each phased Human Pangenome Reference Consortium Release 2 (HPRC2) assembly, and methylation and CpG density were measured in the homologous haplotype-specific interval. Each T2T-CHM13 interval, together with its homologous haplotype-specific intervals, defines a common unit for downstream analyses of genetic-methylation associations, population methylation variation and allele-aware methylation quantitative trait locus (mQTL) mapping. b, Representative chromosome 8 interval (CHM13 chr8:9,268,200–9,272,200). HG00099 hap1 carries a 16-bp insertion and a 332-bp Alu insertion, NA19468 hap2 carries both insertions within a large inversion, and the HG01252 maternal haplotype carries an 88-bp deletion within the same inverted region. Grey ribbons connect homologous sequence; red and blue ribbons mark inserted and deleted sequence, respectively. c, Autosomal 1-Mb summaries of methylation variability (meVar), nucleotide diversity (SNP π) and synteny diversity (SynDiv), which quantifies variation in collinearity among homologous projections, together with representative positions of highly variable methylation regions (HVMRs), highly variable CpG-density regions (HVDRs) and shared HVMR/HVDR intervals, defined as intervals called in both analyses. Green squares, orange triangles and purple circles mark HVMRs, HVDRs and shared HVMR/HVDR intervals, respectively; grey segments denote excluded regions. Chromosome interiors show the log₂-transformed median of scaled meVar across constituent 200-bp intervals. Teal profiles to the left show SNP π estimated directly in 1-Mb windows, whereas purple profiles to the right show mean SynDiv across constituent 200-bp intervals. Scaled meVar is the observed residual variance divided by its mean-dependent expected value. d, Genomic-feature enrichment of 24,022 HVMRs, 3,869 HVDRs and 45 shared HVMR/HVDR intervals. Flow widths show fractional membership across genomic-feature groups. Dot color gives log₂ observed-to-expected overlap, and dot area gives −log₁₀ Benjamini–Hochberg-adjusted *P*. e, Covariate-adjusted partial-residual curves relating residualized log₂(scaled meVar) to SNP π (top) or SynDiv (bottom), genome-wide and across seven genomic-feature groups. Diversity values were transformed as ln[1 + value/median(non-zero value)]. Vertical dashed lines delimit the main, tail and extreme-tail ranges, and shading marks the sparsely represented predictor tails. For SNP π, the main (<2), tail (2–3) and extreme-tail (>3) ranges contained 90.12%, 9.39% and 0.49% of the analysis set, respectively. For SynDiv, the main (<1), tail (1–4) and extreme-tail (>4) ranges contained 95.59%, 3.53% and 0.88%, respectively. f, Feature-group estimates from the same models. Red and blue squares show the incremental weighted *R*² attributable to SNP π and SynDiv, respectively (Δweighted *R*² × 10⁻³); horizontal lines are 95% intervals from 1,000 chromosome-stratified 5-Mb block-bootstrap replicates, and arrows indicate intervals truncated by the plotting range. The two right columns show 200-bp partial Spearman correlations; color indicates direction and magnitude, and point area denotes the block-bootstrap false-discovery-rate (FDR) class for feature-group estimates and the block-bootstrap *P* value for genome-wide estimates.

Across 460 haplotypes, projection recovered a median of 2.86 Gb of projected sequence per haplotype (**Extended Data Fig. 1a–c**). Overall, 97.27% of projected intervals remained close to 200 bp, whereas longer intervals showed secondary peaks at lengths characteristic of Alu and L1 elements, consistent with the retention of haplotype-specific inserted sequence (**Extended Data Fig. 1d**). Projected methylation profiles retained expected patterns around transcription start sites, CGIs and candidate cis-regulatory elements (**Extended Data Fig. 2**).

We quantified CpG sharing across haplotypes by defining CpGs mapping to the same position in the pangenome graph as homologous, even if they were absent from CHM13. After excluding CpGs in centromeric and satellite-repeat sequence and CpGs observed on only one haplotype (singletons), we characterized 35,200,252 distinct CpG sites, of which 17,933,898 (50.95%) were present on all 460 haplotypes (**Extended Data Fig. 1e,f**). Novel CpGs did not saturate by 460 haplotypes.

We then asked whether regions with variable methylation were the same regions in which CpG density varied among haplotypes. To address this, we quantified genome-wide variability in methylation and CpG density across haplotypes (Methods; **Fig. 1c** and **Extended Data Fig. 3a**). We identified 24,022 highly variable methylation regions (HVMRs) and 3,869 highly variable CpG-density regions (HVDRs) at a false-discovery rate (FDR) < 0.05 (Methods; **Fig. 1c** and Supplementary Table 2). Only 45 regions were shared between HVMRs and HVDRs. HVMRs were enriched in regulatory sequence, most strongly at CGIs (13.61-fold), CTCF-bound proximal enhancer-like elements (9.48-fold) and CTCF-bound promoter-like elements (8.16-fold), whereas HVDRs were concentrated in structurally variable sequence, including AluY insertions (60.28-fold), SINE-VNTR-Alu (SVA) insertions (25.35-fold) and structural insertions overall (16.20-fold; **Fig. 1d**).

Methylation across HVMRs separated maternal and paternal haplotypes by uniform manifold approximation and projection (UMAP; **Extended Data Fig. 4a,b**). Of the 1,879 intervals overlapping 118 high-confidence parent-of-origin methylation markers, 588 were HVMRs, representing a 153.70-fold enrichment (one-sided hypergeometric *P* = 2.8 × 10⁻¹⁰⁸²; Supplementary Table 3), whereas none was an HVDR. Despite this enrichment, the 588 parent-of-origin HVMRs accounted for only 2.45% of all HVMRs. Superpopulation structure was also visible in the HVMR methylation profiles: maternal haplotypes formed distinct African and East Asian clusters, whereas paternal haplotypes formed a distinct African cluster (**Extended Data Fig. 4b**).

We evaluated whether population-associated methylation differences were concentrated within HVMRs. We defined population differentially methylated regions (population DMRs) by comparing African (AFR), Admixed American (AMR), East Asian (EAS), European (EUR) and South Asian (SAS) haplotypes separately with all remaining haplotypes (Methods). These superpopulation-specific comparisons identified 217,637 population DMRs (Supplementary Table 4). Only 1,596 population DMRs (0.73%) fell within an HVMR, and 1,513 HVMRs (6.30%) overlapped at least one population DMR. Population DMRs were instead broadly enriched in genic regions, CGIs, candidate cis-regulatory elements, SV and mobile-element-insertion (MEI) regions, and segmental duplications (Methods; **Extended Data Fig. 4c,d**).

We tested whether greater genetic diversity was associated with greater methylation variability. Because the number of variants within a region does not capture how divergent the haplotypes are, we used two complementary measures of local genetic diversity: nucleotide diversity (SNP π) to quantify base-level sequence divergence and synteny diversity (SynDiv) to quantify differences in local sequence collinearity, capturing larger structural differences (Methods). We modelled methylation variability jointly as a function of SNP π and SynDiv using generalized additive models (Methods). Genome-wide, SNP π explained little methylation variability (Δ*R*² = 1.16 × 10⁻³; block-bootstrap *P* = 0.0020) and was weakly inversely correlated with methylation variability (partial Spearman ρ = −0.064; block-bootstrap *P* = 0.0020). SynDiv also explained little methylation variability (Δ*R*² = 0.06 × 10⁻³; block-bootstrap *P* = 0.14) and was not correlated with methylation variability genome-wide (ρ = −0.002; block-bootstrap *P* = 0.400; **Fig. 1f**). Thus, methylation variability generally did not increase with local genetic diversity. We define this pattern as epigenetic buffering.

Across 90.12% of genomic regions, methylation variability decreased as SNP π increased. In the 9.88% of genomic regions with the highest SNP π, however, methylation variability increased as SNP π increased, indicating that epigenetic buffering breaks down at extreme nucleotide diversity (**Fig. 1e**). Six of the seven genomic contexts analyzed separately reached their highest methylation variability in regions with the highest SNP π; MEIs were the exception: methylation variability decreased as SNP π increased (**Fig. 1e** and **Supplementary Fig. 1**).

Across 95.59% of genomic regions, methylation variability remained largely stable as SynDiv increased (**Fig. 1e**). Structural insertions showed the largest contribution of SynDiv to methylation variability (Δ*R*² = 4.61 × 10⁻³), 74-fold above the genome-wide value (**Fig. 1f**). SVA elements showed the strongest epigenetic buffering for both measures: methylation variability decreased as SNP π increased (ρ = −0.28) and as SynDiv increased (ρ = −0.14; **Extended Data Fig. 3b** and **Supplementary Fig. 2**).

### Structural variants create local departures from genome-wide epigenetic buffering

Given the pronounced coupling between synteny diversity and methylation variability within insertion regions, we systematically characterized how major SV classes reshape local DNA methylation. We analyzed 107,040 insertions, 77,260 deletions and 126 inversions (Methods). For insertions and deletions, we compared methylation between carrier and non-carrier haplotypes at homologous CpGs flanking each breakpoint and identified DMRs (Methods). We quantified DMR burden in 100-bp windows flanking each breakpoint as the fraction of tested SVs associated with a DMR in each window. DMR burden was highest near breakpoints, with opposite directional biases for insertions and deletions. Immediately adjacent to insertion breakpoints, 7.55% of insertions were next to a hypermethylated DMR, compared with 2.75% with a hypomethylated DMR; for deletions, 8.54% had a hypomethylated DMR and 3.23% a hypermethylated DMR (**Fig. 2a,b**). DMR burden declined rapidly as a function of distance from the breakpoints, reaching 0.67–1.05% at 1–3 kb. Overall, 20,443 insertions (19.10%) and 18,261 deletions (23.64%) were associated with at least one DMR (**Fig. 2f**). Insertion-associated DMRs were nearly evenly divided between hypermethylation (50.77%) and hypomethylation (49.23%), whereas deletion-associated DMRs were predominantly hypomethylated (60.06%; **Fig. 2g**). At the *IFI44L* locus, a 91-bp deletion carried by seven haplotypes was associated with a focal hypermethylated DMR near the promoter, with a methylation difference of 0.372 (**Fig. 2d**). *IFI44L* promoter methylation is a sensitive blood biomarker of systemic lupus erythematosus^32^, and all seven *IFI44L* transcripts detected in matched long-read transcriptome data showed lower abundance on deletion-carrying than non-carrier haplotypes (Supplementary Table 5).

**Fig. 2.**
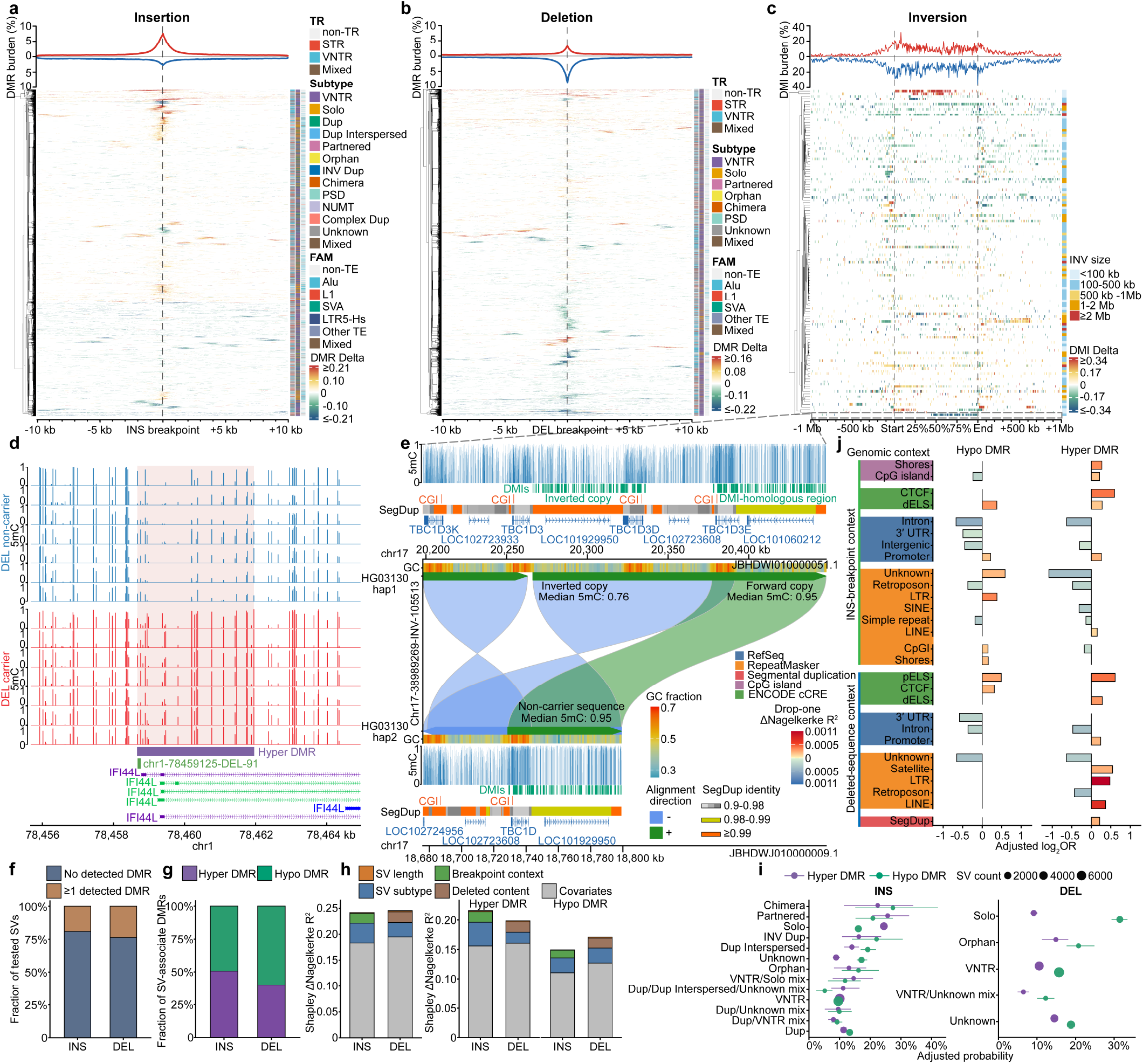
Methylation dynamics across structural-variant architectures and genomic contexts. a,b, Breakpoint-relative burden of hypermethylated and hypomethylated differentially methylated regions (DMRs) for insertions (INS; *n* = 107,040) and deletions (DEL; *n* = 77,260), shown in a and b, respectively. DMR burden is the fraction of structural variants (SVs) tested in each 100-bp window with an overlapping carrier-versus-non-carrier DMR; windows were stepped by 50 bp across 10-kb flanks on either side of each breakpoint. Hypermethylated and hypomethylated burdens are plotted above and below zero, respectively. Rows represent individual insertions (a) or deletions (b), clustered by carrier-minus-non-carrier methylation differences across callable flanking windows; DMR-overlapping windows are colored by methylation difference. The three sidebars, from left to right, show tandem-repeat (TR) class, SV subtype and transposable-element (TE) family (FAM). c, Burden of hypermethylated and hypomethylated differentially methylated intervals (DMIs) across normalized inversion (INV) bodies and physical flanks extending 1 Mb from each breakpoint (*n* = 126 INVs). INV bodies are scaled from start to end, whereas the flanks retain physical distance. Hypermethylated and hypomethylated DMI burdens are plotted above and below zero, respectively. Rows represent individual inversions clustered by carrier-minus-non-carrier methylation differences across the inversion body and flanks; DMI intervals are colored by methylation difference. The right sidebar denotes inversion-size class. d, A 91-bp deletion near the *IFI44L* promoter associated with a focal hypermethylated DMR. The deletion was carried by 7 of 460 haplotypes; tracks show the seven within-individual carrier-non-carrier haplotype comparisons. Pink shading marks the focal DMR. The maximum absolute methylation difference was 0.372. e, Complex inversion-associated duplication in the *TBC1D3* family region (chr17:38,989,269–39,094,781; 105,513 bp) in HG03130, carried by 24 of 460 haplotypes. The inversion-carrying haplotype contains two copies of a segment homologous to the corresponding non-carrier sequence, one in inverted orientation and one in forward orientation. Blue and green ribbons denote reverse- and forward-orientation alignments, respectively. Overlapping carrier-path sequence was excluded before comparison of the non-carrier, forward and inverted segments. The three segments contained 11.56–11.72 assayed CpGs per kb. Median methylation was 0.76 in the inverted copy and 0.95 in both the forward copy and corresponding non-carrier sequence. f, Proportions of tested SVs associated with at least one DMR: 20,443 of 107,040 INS (19.10%) and 18,261 of 77,260 DEL (23.64%). g, Proportions of individual INS- and DEL-associated DMRs that were hypermethylated or hypomethylated (INS, 50.77% and 49.23%; DEL, 39.94% and 60.06%, respectively). h, Shapley decomposition of full-model Nagelkerke pseudo-*R*² for occurrence of at least one DMR (left) and for hypermethylated or hypomethylated DMR occurrence (right). Models contained SV length, SV subtype and the genomic context of the insertion breakpoint for INS or the deleted sequence for DEL; allele frequency and the number of tested CpGs were included as covariates. i, Covariate-adjusted probabilities of hypermethylated or hypomethylated DMR occurrence by SV subtype. Purple and green denote hypermethylated and hypomethylated DMRs, respectively, and point size indicates the number of SVs in each subtype. Points are standardized model estimates and horizontal lines are 95% bootstrap confidence intervals from 200 SV-level resamples. j, Genomic-context effects on DMR occurrence in directional models. Bars show adjusted log₂ odds ratios (ORs) for INS-breakpoint context (top) and deleted-sequence context (bottom), with hypomethylated and hypermethylated DMRs shown in the left and right columns, respectively. Blue and red denote decreased and increased DMR occurrence, respectively, and color intensity gives the unsigned drop-one ΔNagelkerke pseudo-*R*². Only terms with Benjamini–Hochberg-adjusted *q* < 0.05 within SV class, DMR direction and genomic-context role are shown.

Inversions were associated with a broader distribution of methylation differences. We tested projected 200-bp intervals across each inversion body and 1 Mb on either side and identified differentially methylated intervals (DMIs; Methods). The proportion of inversions with a DMI remained elevated across the inversion body and within approximately 25 kb of each breakpoint, before declining in the 250-kb to 1-Mb flanks (**Fig. 2c**). Among these, an inversion at the *TBC1D3* locus on chromosome 17 showed a series of hypomethylated DMIs within the inversion body (**Fig. 2c,e**). The *TBC1D3* gene family spans two segmental-duplication clusters at 17q12 that differ by as many as 20 copies between human haplotypes^29,33^. The 105,513-bp inversion relative to CHM13 was carried by 24 of 460 haplotypes. In HG03130, relative to the non-carrier haplotype, the inversion-carrying haplotype contained two copies of a 56.5-kb segment homologous to the corresponding segment on the non-carrier haplotype, one in forward and one in inverted orientation (**Fig. 2e**). Both copies were highly similar in sequence to the corresponding segment on the non-carrier haplotype (99.71% and 99.72% identity) with similar CpG density (11.56 to 11.72 CpGs per kb). Median methylation was 0.95 in both the forward copy and the corresponding non-carrier copy, but 0.76 in the inverted copy. Thus, highly similar copies of the same homologous segment may display markedly different methylation states associated with a large inversion.

Because SVs differ in variant subtype, size and local genomic context, we asked which of these properties best explained DMR occurrence. We modelled DMR occurrence as a function of variant subtype, SV length and the local genomic context of the SV and used Shapley decomposition^34^ to compare their contributions (Methods). Variant subtype accounted for 65.6% of the explained variation in insertions and 54.7% in deletions and remained the largest contributor when hypermethylated and hypomethylated DMRs were analyzed separately (49.0– 67.0%; **Fig. 2h**). Among the subtypes, solo insertions were more often associated with hypermethylated than hypomethylated DMRs, whereas solo deletions were more often associated with hypomethylated than hypermethylated DMRs (**Fig. 2i**). DMR occurrence also varied with the local genomic context. Insertion breakpoints in regulatory sequence were preferentially associated with hypermethylated DMRs, whereas intronic and intergenic breakpoints were associated with lower DMR occurrence. Deletions removing proximal enhancer-like sequence were associated with increased occurrence of both hypermethylated and hypomethylated DMRs (**Fig. 2j**). Retroposons were consistently associated with reduced DMR occurrence, consistent with the strong epigenetic buffering observed in **Fig. 1f**. SV length accounted for only 2.7% of the explained variation in insertions and 4.7% in deletions. Thus, whether an SV was associated with a DMR depended much more strongly on its variant subtype and local genomic context than on its size (Supplementary Table 6).

We also examined whether DMR occurrence varied with the repeat composition of an SV. Insertions carrying transposable-element (TE) sequence were more likely than other insertions to carry DMRs, with a stronger association with hypermethylation (**Extended Data Fig. 5a,b**). Deletions removing TE sequence showed a different pattern: relative to deletions without TE sequence, they were more likely to carry hypomethylated DMRs and less likely to carry hypermethylated DMRs (**Extended Data Fig. 5a,b**). Insertions and deletions classified as short tandem repeats (STRs) or variable-number tandem repeats (VNTRs) were less likely to carry DMRs than SVs not classified as tandem repeats, in both methylation directions (**Extended Data Fig. 5c,d**). Thus, SV repeat composition was associated with both DMR occurrence and whether DMRs were hypermethylated or hypomethylated.

Beyond DMR occurrence, we examined DMR count and magnitude. We defined DMR count as the number of DMRs associated with an SV, and DMR magnitude as the absolute methylation difference between carrier and non-carrier haplotypes for each SV-DMR pair. Variant subtype again made the largest contribution to DMR count, accounting for 61.4% of the explained variation in insertions and 46.7% in deletions. Repeat composition also contributed substantially, particularly to hypermethylated-DMR counts in insertions (**Extended Data Fig. 5e,f,i,j**). DMR count also differed according to the local genomic context of the SV (**Extended Data Fig. 5h** and Supplementary Table 7). In contrast, once a DMR was present, properties of the DMR itself, including its genomic context, CpG count, length and distance from the nearest SV breakpoint, accounted for 75.7–76.0% of the explained variation in DMR magnitude for insertions and 72.9–84.0% for deletions. DMRs overlapping regulatory sequence or CGIs were associated with larger absolute methylation differences, whereas DMRs overlapping repeat sequence showed smaller differences, particularly at retroposons (**Supplementary Fig. 3** and Supplementary Table 8). By comparison, the repeat composition of the SV contributed little to DMR magnitude (**Extended Data Fig. 5g,k,l**). Thus, variant subtype, repeat composition and genomic context primarily distinguished whether and how many DMRs occurred, whereas properties of the DMR itself more strongly determined DMR magnitude.

Finally, we examined whether methylation differences in the insertion-containing interval extend into the surrounding sequence. For 58,038 insertions, we compared methylation between carrier and non-carrier haplotypes separately in the projected 200-bp insertion-containing interval and the 1-kb flanking sequence (Methods). The methylation difference in the insertion-containing interval was only moderately correlated with the difference in flanking regions (Spearman ρ = 0.297). Among 2,249 insertions showing significant methylation differences in the insertion-containing interval or the 1-kb flanking sequence, 2,220 were significant in the insertion-containing interval, 48 in the flanks and only 19 in both (**Supplementary Fig. 4**). Thus, methylation differences within insertions rarely extended into the flanking sequence.

### CGIs are polymorphic features of the human genome

Because CGIs are defined by sequence^4^, genetic variation frequently creates, removes or repositions CGIs across homologous haplotypes. Differences in genome sequence alter CGI representation and consequently affect the measurement of CGI methylation^35^. A fixed CGI annotation from a single reference genome therefore cannot represent CGI presence, absence and sequence variation across human genomes. We developed the second framework, PanCGI, to integrate CGI calls made independently in each phased assembly into a common pangenome catalogue (**Fig. 3a**).

**Fig. 3.**
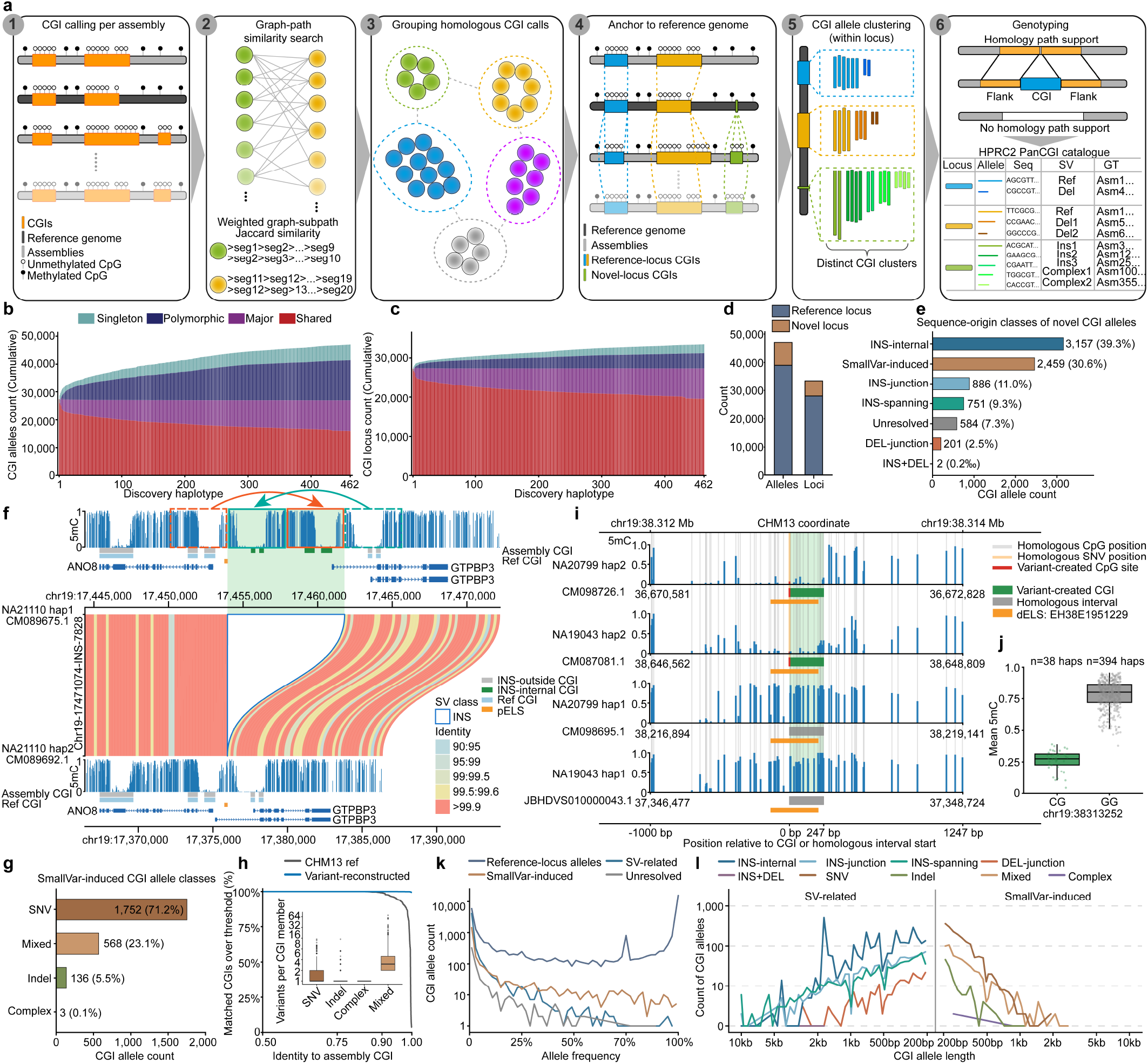
CpG-island polymorphism and its sequence origins in the human pangenome. a, PanCGI workflow. CpG islands (CGIs) were called independently in each phased assembly, represented as paths through the pangenome graph and grouped by graph-path similarity. These groups were anchored to CHM13 and consolidated into reference or novel CGI loci. Each assembly CGI assigned to a locus is termed a member; within each locus, member sequences were clustered by sequence similarity, with one representative member defining each CGI allele, after which locus and allele genotypes were assigned from homology-path support. b,c, Cumulative discovery of CGI alleles (b) and loci (c) across 462 HPRC2 haplotype assemblies, including both haplotypes of HG002. Colors distinguish singleton, polymorphic, major and shared states. At each cumulative haplotype rank *k*, classes were recalculated from carrier count AC and carrier fraction *f* = AC/*k*: singleton, AC = 1 and *f* < 1; polymorphic, AC > 1 and *f* < 0.5; major, AC > 1 and 0.5 ≤ *f* < 1; and shared, *f* = 1. d, The catalogue contains 47,039 alleles at 33,463 loci, including 8,040 alleles at 5,451 loci without an annotated CHM13 CGI. Bars partition allele and locus counts into reference and novel loci. e, Mutually exclusive sequence origins of the 8,040 novel-locus alleles. CGIs contained within inserted sequence (INS-internal), crossing an insertion junction (INS-junction), spanning an inserted segment (INS-spanning), spanning a deletion junction (DEL-junction), or associated with both an insertion and a deletion (INS+DEL) define the structural variant (SV)-related classes; SmallVar-induced alleles are attributable to small variants, whereas unresolved alleles could not be assigned a sequence origin. f, A 7,854-bp haplotype insertion between the adjacent *ANO8* and *GTPBP3* promoters comprising duplicated sequence derived from both sides of the insertion site. The insertion contains four CGI members representing two CGI alleles. Green marks assembly CGIs within the insertion, grey assembly CGIs outside it, light blue reference CGIs and orange proximal enhancer-like signature (pELS); the blue outline marks the insertion, and ribbon color denotes pairwise sequence identity. g, SmallVar-induced alleles classified as single-nucleotide variant (SNV), mixed, insertion or deletion (indel), or complex. h, Reverse cumulative distributions of sequence identity for 132,097 matched member sequences from SmallVar-induced alleles, comparing each assembly CGI with either its homologous CHM13 interval or its variant-reconstructed sequence. Black and blue denote the CHM13 and variant-reconstructed sequences, respectively. The inset shows the number of small variants required to account for the sequence difference for each matched CGI member; boxes show medians and interquartile ranges and whiskers extend to 1.5 times the interquartile range. i, A CpG-creating substitution in a distal enhancer-like signature (dELS). Blue bars show CpG methylation across four representative haplotypes; grey and pale-orange vertical lines mark homologous CpG and SNV positions, respectively. Red marks the variant-created CpG, green the variant-created CGI, grey the homologous interval and orange the dELS. j, Methylation by sequence state at the locus in i. Methylation was averaged across the assembly-defined CGI interval in CGI-positive haplotypes and across the corresponding homologous interval in CGI-negative haplotypes. CGI-positive haplotypes had mean methylation of 0.261 (median, 0.275; *n* = 38), compared with 0.781 (median, 0.800; *n* = 394) for CGI-negative haplotypes. Boxes show medians and interquartile ranges; whiskers extend to 1.5 times the interquartile range and points denote haplotypes. k, Allele-frequency distributions for reference-locus, SV-related, SmallVar-induced and unresolved CGI alleles. l, Allele counts by CGI length, stratified by SV-related sequence-origin class (left) and SmallVar-induced variant class (right).

PanCGI uses the pangenome graph to group homologous CGI calls across assemblies into loci. A locus overlapping an annotated CHM13 CGI was defined as a reference locus, whereas one without an annotated CHM13 CGI was defined as a novel locus. Each CGI call assigned to a locus was termed a member. Within each locus, CGI members may differ in sequence, and we therefore clustered them based on sequence similarity and length. A representative member from each cluster was defined as a CGI allele. A locus can therefore contain multiple CGI alleles, and each allele may contain multiple members. Each assembly was then genotyped at every locus as CGI-positive if it had a member, CGI-negative when the homologous locus was recoverable but no CGI was called, or missing when the homologous locus itself could not be reliably identified. On CHM13, the CGI-calling procedure reproduced the UCSC annotation exactly on unmasked sequence and remained highly concordant after repeat masking (**Extended Data Fig. 6a–d**).

Across 462 HPRC2 haplotypes, including both haplotypes of HG002, PanCGI identified 47,039 CGI alleles at 33,463 loci, with both allele and locus discovery continuing to rise as additional haplotypes were added (**Fig. 3b,c** and Supplementary Table 9). Among these, 8,040 alleles at 5,451 loci were novel to CHM13 (**Fig. 3d**). SVs accounted for 4,997 alleles (62.15%): 3,157 occurred entirely within inserted sequence (INS-internal), 886 formed at insertion junctions (INS-junction), 751 spanned insertion boundaries (INS-spanning), 201 formed at deletion junctions (DEL-junction) and 2 involved a combined insertion and deletion (**Fig. 3e** and **Extended Data Fig. 7a**). Small variants (shorter than 50 bp) accounted for 2,459 alleles (30.58%; SmallVar-induced), leaving 584 unresolved.

We showed a 7.8-kb insertion adjacent to the *ANO8* and *GTPBP3* promoters that duplicated sequence from both sides of the insertion site. Each duplicated segment contained two CGI members representing one CGI allele, resulting in two novel CGI alleles at this locus (**Fig. 3f**). Each inserted CGI showed methylation similar to its corresponding source copy, despite being located at a different genomic position, in contrast to the marked methylation divergence observed after sequence inversion at the *TBC1D3* locus in **Fig. 2e**. Thus, distinct forms of sequence reconfiguration can be associated with markedly different effects on local DNA methylation. More broadly, CGI polymorphism occurred at promoters, including *TERT*, *TGFB1* and *ACTG1* at reference loci and *POLDIP3* and *ESPNL* at novel loci (**Extended Data Fig. 7b**).

We examined the sequence changes underlying the 2,459 SmallVar-induced alleles. Most were attributable to single-nucleotide variants (SNVs; 1,752, 71.25%), followed by alleles involving at least two classes of small variants (mixed; 568, 23.10%), short insertion/deletion (indel) alleles (136, 5.53%) and complex alleles (3, 0.12%) (**Fig. 3g**). To determine how many sequence changes were required to create these alleles, we counted the small-variant differences between each CGI-positive member and its homologous CHM13 sequence. SNV-induced alleles required a median of one small variant (interquartile range, 1–2), indel-induced alleles a median of one, and mixed alleles a median of three (interquartile range, 2–5) (**Fig. 3h**). Thus, only a small number of sequence changes were typically required to alter CGI state. For example, at a distal enhancer-like element, a single substitution created a CpG and changed the homologous sequence from CGI-negative to CGI-positive (**Fig. 3i**). The 38 CGI-positive haplotypes showed substantially lower methylation across the CGI than the 394 CGI-negative haplotypes across the corresponding homologous interval (mean, 0.261 versus 0.781; **Fig. 3j**).

CGI alleles arising through different mechanisms also differed in length and population frequency. INS-internal alleles were generally longer and rarer, with a median length of 530 bp and a median allele frequency of 0.0022, whereas SmallVar-induced alleles were shorter and more common, with corresponding medians of 255 bp and 0.0108 (**Fig. 3k,l** and **Supplementary Fig. 5**). INS-spanning and DEL-junction alleles were more frequent, with median allele frequencies of 0.0130 and 0.0152, respectively.

We next characterized the population distribution of CGI presence-absence variation (PAV). Principal-component analysis based on CGI PAV separated the five superpopulations at both locus and allele resolution (**Extended Data Fig. 7c,d**). Across five CGI sets comprising polymorphic loci, novel loci, novel alleles, SV-related novel alleles and SmallVar-induced novel alleles, 74.4–79.7% of PAV variance reflected differences among individuals within the same population, with the remaining 20.3–25.6% attributable to differences between population groups (**Supplementary Fig. 6**).

Population differentiation also varied with the sequence origin of CGI alleles. SmallVar-induced novel alleles showed higher pairwise Hudson’s *F* ^36^, a measure of allele-frequency differentiation between population groups, than SV-related novel alleles in every superpopulation comparison (**Extended Data Fig. 8a,b**), with the largest contrast between AFR and EAS (0.147 versus 0.097). Among SmallVar-induced alleles, mixed and SNV classes showed the highest differentiation between AFR and EAS, whereas indel alleles showed the highest differentiation between AFR and SAS (**Extended Data Fig. 8b**). These population-frequency differences were also reflected in the alleles carried by each individual: AFR individuals carried a larger fraction of INS-junction, INS-spanning, SNV and mixed alleles than individuals from the other superpopulations (**Extended Data Fig. 7e** and **Supplementary Fig. 7**).

### Local genealogies resolve CGI gain, loss and turnover

Many SmallVar-induced loci differed between CGI-positive and CGI-negative haplotypes by only a few sequence changes, suggesting that CGI state can change around sequence that is already CpG-rich. This presents an opportunity to investigate the recent evolutionary history of CGIs. We thus analyzed the 2,279 SmallVar-induced novel loci using local genealogies reconstructed from phased autosomal SNVs^37^ to infer ancestral CGI states and reconstruct CGI gain and loss. A local genealogy relates the 462 haplotypes through their inferred common ancestry at a genomic position, allowing changes between CGI-positive and CGI-negative states to map onto its branches. We used weighted parsimony^38^ to infer the CGI state of the ancestor and the branches on which state changes occurred (Methods). Of the 2,279 loci, 2,166 had an evaluable local genealogy and 113 lacked a usable genealogy. Of the evaluable loci, 753 had a CGI-positive ancestor, 1,281 had a CGI-negative ancestor and 132 were equivocal (**Fig. 4a** and **Supplementary Fig. 8**).

**Fig. 4.**
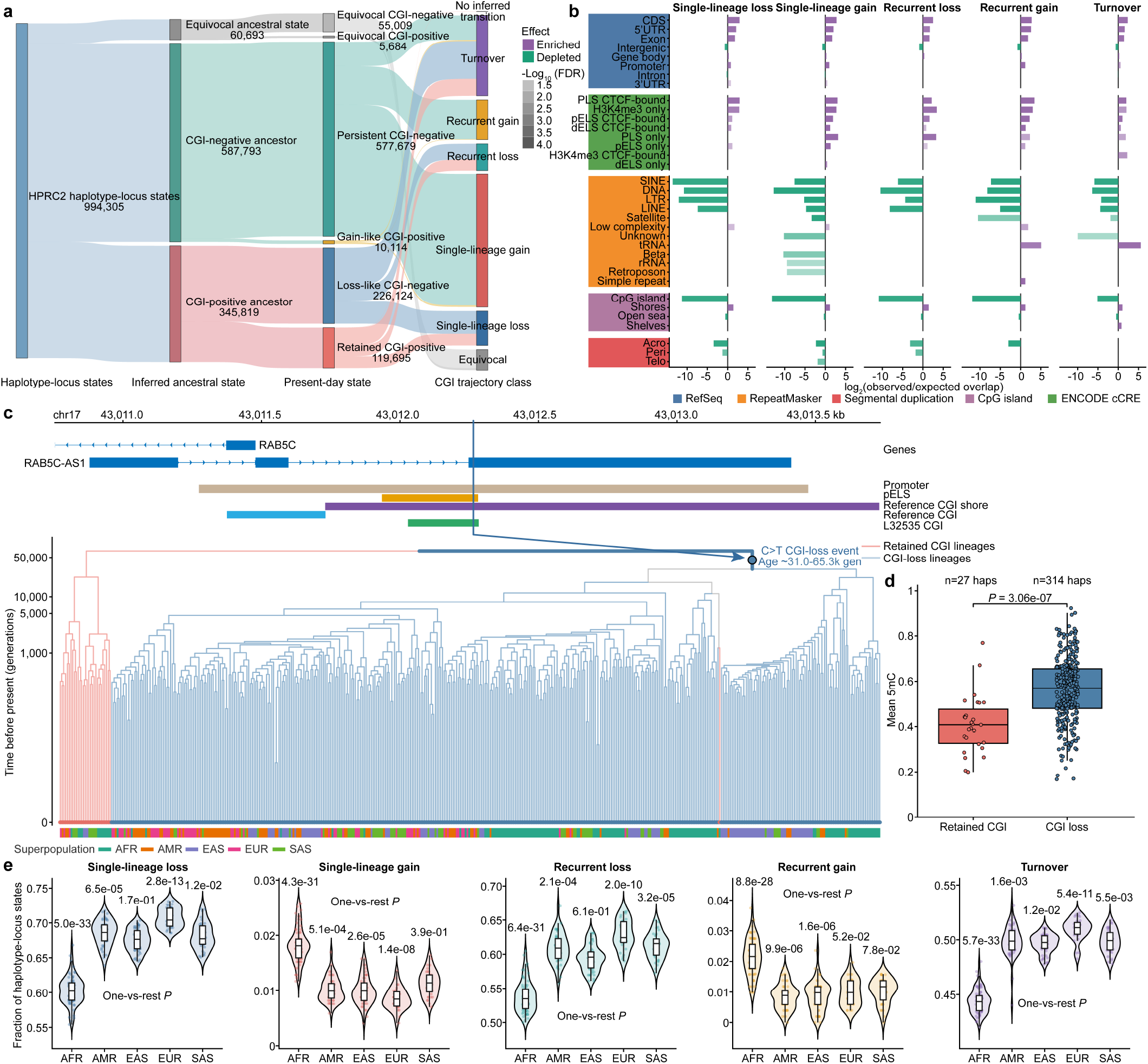
Genealogical trajectories of CpG-island gain, loss and turnover. a, Flow of CpG-island (CGI) states across 994,305 haplotype-locus states at 2,166 loci with an evaluable local genealogy. A haplotype-locus state is the CGI state of one haplotype at one locus. Ribbon width gives the number of haplotype-locus states passing through the inferred CGI-positive, CGI-negative or equivocal ancestral state, present-day state and mutually exclusive trajectory class. For loci assigned a CGI-positive ancestor, present-day states are classified as retained CGI-positive or loss-like CGI-negative; for loci assigned a CGI-negative ancestor, they are classified as gain-like CGI-positive or persistent CGI-negative. The final layer comprises single-lineage loss, single-lineage gain, recurrent loss, recurrent gain, turnover, equivocal loci and loci with no inferred transition. Loci for which the CGI-negative and CGI-positive ancestral-state assignments had equal parsimony costs are shown as equivocal. b, Genomic enrichment and depletion of the five trajectory classes: 222 single-lineage-loss, 857 single-lineage-gain, 172 recurrent-loss, 258 recurrent-gain and 524 turnover loci. Bars show log₂ observed-to-expected nucleotide overlap from Genomic Association Tester (GAT) analyses. Purple and green indicate enrichment and depletion, respectively, and opacity gives −log₁₀(*q*), where *q* is the Benjamini–Hochberg false-discovery-rate (FDR)-adjusted *P* value; only terms with *q* ≤ 0.05 are shown. Genomic features comprise RefSeq gene annotations, ENCODE candidate cis-regulatory elements (cCREs), RepeatMasker classes, reference CGI contexts and segmental duplications. Within cCREs, the labels denote promoter-like signatures (PLS), proximal enhancer-like signatures (pELS) and distal enhancer-like signatures (dELS). c, Local genealogy and genomic features at the *RAB5C* locus L32535. The tree contains 462 haplotypes and supports a CGI-positive ancestor, with 30 retained-CGI and 432 CGI-loss present-day states. Red and blue branches denote reconstructed retained-CGI and CGI-loss lineages, respectively, and terminal dots indicate present-day CGI state. The vertical axis is estimated time before present in generations. The derived C>T allele at chr17:43,012,266 lies on the branch associated with CGI loss, with an inferred child-to-parent interval of 30,990–65,308 generations. The aligned track marks *RAB5C*, its promoter, a pELS, the reference CGI shore, the novel CGI locus L32535 and the focal variant; the lower strip shows superpopulation in tree-leaf order. d, Haplotype methylation at L32535 by present-day CGI state. Mean 5-methylcytosine (5mC) across graph-linked CpGs was estimated for 27 retained-CGI and 314 CGI-loss haplotypes. Median methylation was 40.8% and 57.0%, respectively (raw two-sided Wilcoxon rank-sum test, *P* = 3.06 × 10⁻⁷). Boxes show medians and interquartile ranges; whiskers extend to 1.5 times the interquartile range and points denote haplotypes. e, Fraction of evaluable haplotype-locus states per individual whose present-day CGI state differed from the inferred ancestral state, shown for the five trajectory classes. The numerator is the number of CGI-negative states at loss-oriented loci or CGI-positive states at gain-oriented loci; turnover loci use their locus-specific inferred ancestral state. The denominator comprises all non-missing haplotype-locus states across both haplotypes of that individual within the indicated class. Each point denotes one of 231 diploid individuals. Violins show the distribution and internal boxes the median and interquartile range. Labels give raw two-sided Wilcoxon rank-sum one-versus-rest *P* values comparing each superpopulation with all remaining individuals within that class. Superpopulations are African (AFR), Admixed American (AMR), East Asian (EAS), European (EUR) and South Asian (SAS).

Comparing the inferred ancestral states with present-day haplotypes revealed a pronounced asymmetry. We treated the CGI state of one haplotype at one locus as a haplotype-locus state. Across the 753 loci with a CGI-positive ancestor, 65.4% of present-day haplotype-locus states had lost the CGI. By contrast, across the 1,281 loci with a CGI-negative ancestor, only 1.7% had gained a CGI (**Fig. 4a**). This asymmetry persisted across all tested weighting schemes for CGI gain and loss (Methods).

Using the ancestral-state assignments, we classified five evolutionary trajectories for each locus: single-lineage gain (1) or loss (2), if the CGI state change occurred on a single genealogy branch; recurrent gain (3) or loss (4), if there were multiple independent changes in the same direction; and turnover (5), if there were both gains and losses. Among loci with a CGI-negative ancestor, 857 showed single-lineage gain and 258 recurrent gain; among loci with a CGI-positive ancestor, 222 showed single-lineage loss and 172 recurrent loss. Another 524 loci showed turnover, while one locus required no CGI state change within the human genealogy (**Fig. 4a**, **Supplementary Fig. 9** and Supplementary Tables 10, 11). CGIs were lost at fewer loci than they were gained, but a loss removed the CGI from most haplotypes at that locus, whereas a gain was usually confined to relatively few haplotypes.

The *RAB5C* promoter illustrates a single-lineage CGI loss. Thirty of the 462 HPRC2 haplotypes carried a CGI that was absent from CHM13, whereas the other 432 haplotypes were CGI-negative (**Fig. 4c**). The genealogy supported a CGI-positive ancestor, making the common CGI-negative state a derived loss. A single C>T substitution marked the lineage carrying this loss. All 432 CGI-negative haplotypes (including CHM13) descended from this branch, which was dated to approximately 30,990 to 65,308 generations ago. Haplotypes retaining the CGI had lower methylation than those with CGI loss (median, 40.8% versus 57.0%; *n* = 27 and 314; two-sided Wilcoxon *P* = 3.06 × 10⁻⁷; **Fig. 4d**).

Single-lineage gains and losses, recurrent gains and losses, and turnover loci were enriched in regulatory and transcribed sequence. Promoter-like elements and coding sequence were enriched across all five trajectories, and DNase-H3K4me3 elements, 5′ untranslated regions (UTRs) and exons were also consistently enriched (**Fig. 4b**). Recurrent gains and turnover loci showed an additional strong enrichment at tRNA-derived repeats, but this enrichment was absent from single-lineage gain and loss loci.

How an individual’s present-day CGI states differed from the inferred ancestor varied among superpopulations. At single-lineage and recurrent loss loci, AFR individuals had lower fractions of CGI loss than the remainder of the cohort (median, 0.602 versus 0.686 and 0.535 versus 0.608, respectively). At single-lineage and recurrent gain loci, AFR individuals instead had higher fractions of CGI gain (0.018 versus 0.010 and 0.022 versus 0.010, respectively; **Fig. 4e** and **Supplementary Fig. 10**).

Locus L31021 linked these evolutionary trajectories to the population DMRs described above. This single-lineage-loss locus overlapped an AFR-hypermethylated DMR (**Extended Data Fig. 8c**). Across superpopulations, CGI-positive frequency increased progressively as median methylation decreased, ranging from 20.0% CGI-positive and 48.25% median methylation in AFR to 54.2% and 39.35%, respectively, in SAS. Ancestral-base polarization identified G as ancestral, and the genealogy placed the G>A substitution associated with CGI loss on a branch dating to approximately 14,522 to 25,822 generations ago. CGI-positive haplotypes carrying the ancestral G allele had median methylation of 28.05% (*n* = 153). Haplotypes that retained ancestral G but carried one or both of two additional C>T substitutions no longer met the CGI criterion and had median methylation of 47.12% (*n* = 57), while other CGI-negative haplotypes had median methylation of 53.97% (*n* = 238). Methylation at this locus tracked whether the local sequence retained CGI status, not simply the allele at the G>A site associated with CGI loss.

### Allele-aware mapping resolves distinct and shared cis-regulatory architectures

Building on the precise local genetic effects on methylation at polymorphic CGI loci, we next resolved the genome-wide cis genetic architecture of the methylome and examined how this architecture relates to transcriptional regulation. To measure transcript output in the same individuals, we assembled a long-read transcript catalogue from 202 matched Iso-Seq datasets (HPRC2 catalogue). The HPRC2 catalogue comprised 35,568 genes (29,683 annotated and 5,885 novel) and 437,568 isoforms (123,002 annotated and 314,566 novel). Merging these models with GENCODE v49^39^ produced a combined catalogue of 84,728 genes and 823,720 isoforms (**Fig. 5a**).

**Fig. 5.**
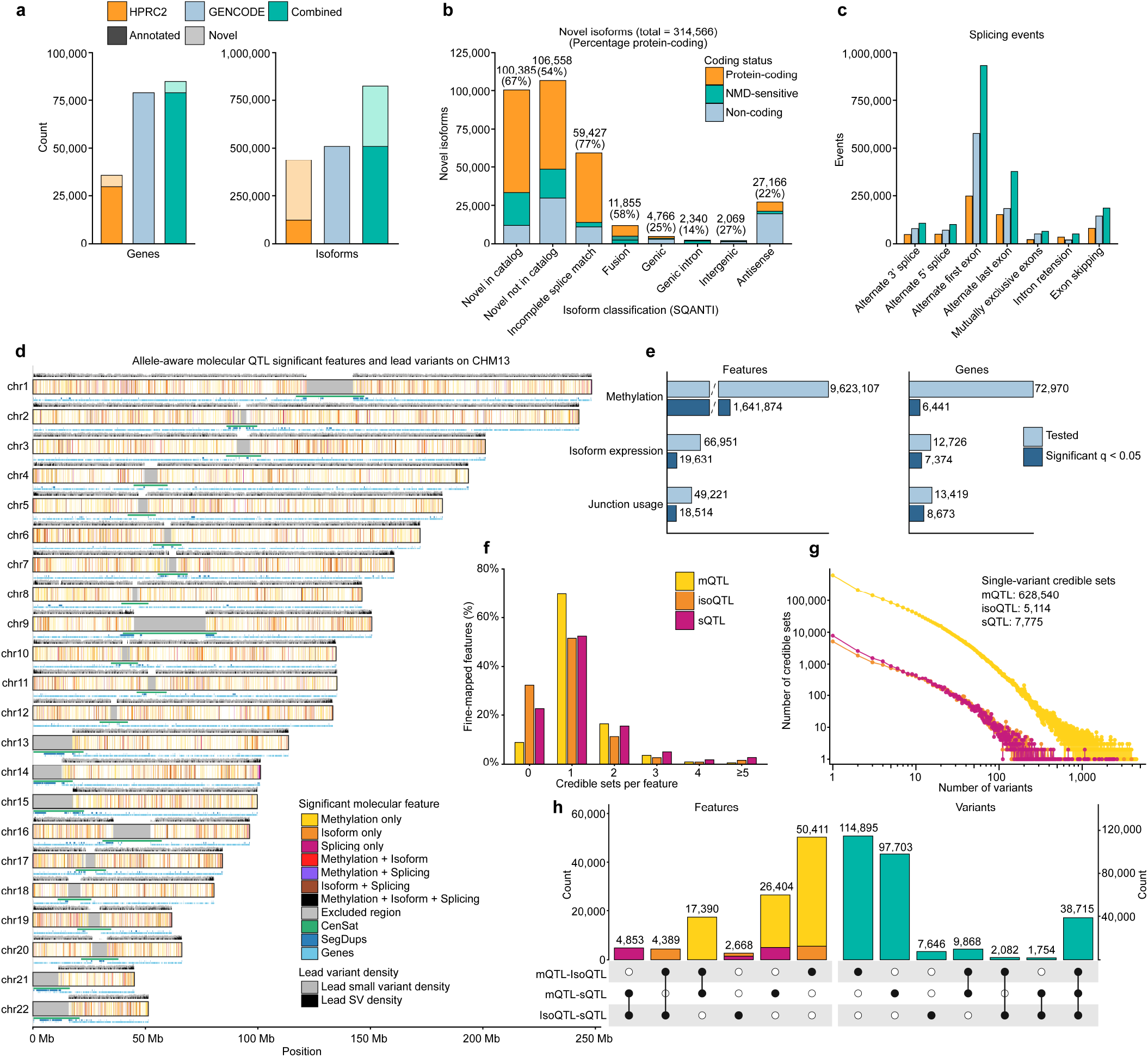
Allele-aware molecular quantitative trait locus (QTL) maps of methylation, isoform abundance and splicing. a, Numbers of gene and isoform models in three GRCh38 catalogues: the HPRC2 catalogue identified from Iso-Seq data for 202 HPRC2 individuals, GENCODE v49 and their combined catalogue. Gene and isoform counts are 35,568 and 437,568 for HPRC2, 78,843 and 509,110 for GENCODE, and 84,728 and 823,720 for the combined catalogue. b, SQANTI3 structural class and predicted coding status of 314,566 novel isoforms, including isoforms predicted to be sensitive to nonsense-mediated decay (NMD). Labels show class totals and percentages predicted to be protein coding. c, Alternative-splicing event counts by class in the three catalogues. d, CHM13 distribution of significant methylation intervals, isoforms and splice junctions and their lead variants. Coloured chromosome tracks show the genomic intervals covered by significant features and overlaps among the three feature classes. Grey and black tracks show the 100-kb densities of unique lead small variants and structural variants (SVs), respectively, pooled across the methylation QTL (mQTL), isoform QTL (isoQTL) and splicing QTL (sQTL) analyses; a lead variant shared among analyses is counted once per window. Lower tracks mark excluded regions, centromere/satellite repeats (CenSat), segmental duplications (SegDups) and genes. e, Numbers of tested and significant (*q* < 0.05) molecular features and associated genes for methylation, isoform abundance and splice-junction usage. f, Percentage of fine-mapped features assigned 0, 1, 2, 3, 4 or ≥5 95% credible sets. g, Distribution of 95% credibl e-set sizes; median sizes were three variants for mQTLs, five for isoQTLs and three for sQTLs. h, UpSet summaries of colocalized molecular features (left) and variants in shared-signal 95% credible sets (right). Filled circles and connecting lines indicate the pairwise QTL colocalization classes represented by each intersection, and bars give the number of distinct features or variants in that intersection. In total, 272,663 unique variants occurred in at least one shared-signal credible set, including 38,715 represented across all three QTL classes.

Of the 314,566 novel isoforms, 185,742 (59.05%) were predicted to be protein coding, 48,182 (15.32%) to be sensitive to nonsense-mediated decay (NMD), and 80,642 (25.64%) to be non-coding. Novel-not-in-catalogue, novel-in-catalogue and incomplete-splice-match isoforms were the largest SQANTI3 structural classes (**Fig. 5b**). The HPRC2 transcript models contained 634,212 alternative-splicing events, which increased to 1,817,914 after merging with GENCODE (**Fig. 5c**). This expansion is consistent with long-read transcriptome studies showing that existing annotations omit many expressed isoforms^40^. Using the combined transcript catalogue, we quantified isoform abundance and splice-junction usage separately on the two phased haplotypes. Expected parent-of-origin expression at established imprinted loci^41,42^ further supported the haplotype assignment of the Iso-Seq data, including maternal enrichment at *KCNQ1* and paternal enrichment at *KCNQ1OT1*, *NAP1L5*, *FAM50B* and *L3MBTL1* (**Supplementary Fig. 11**).

Combining the phased isoform abundance and splice-junction usage with methylation phenotypes defined by our framework for direct comparison across homologous regions, we mapped cis molecular quantitative trait loci (QTLs) in an allele-aware manner^43^. Allele-aware mapping combines genotype-phenotype association across individuals with allelic imbalance between the two haplotypes within each individual. For methylation and junction usage, we developed coverage-weighted ratio-based formulations (Methods). Methylation testing included 9,623,107 projected methylation intervals spanning 66.35% of autosomal CHM13, with 97.78 Mb in segmental duplications, 127.42 Mb in centromeric and satellite (CenSat) sequence and 9.36 Mb of CHM13 sequence absent from GRCh38. Isoform testing included 66,951 transcripts across 12,726 genes, including 34,412 novel isoforms, and splicing testing included 49,221 junctions across 13,419 genes (**Supplementary Fig. 12**). The mQTL analysis tested 19,306,360 variants, including 105,814 insertions, 77,038 deletions and 294 inversions. Isoform and splicing QTL analyses used the corresponding 201-individual panel of 19,304,602 variants after cohort-specific missingness filtering.

We detected significant cis associations for 1,641,874 methylation intervals (17.06% of those tested), 19,631 isoforms (29.32%), including 8,697 novel isoforms, and 18,514 splice junctions (37.61%; **Fig. 5d,e** and Supplementary Tables 12–14). Significant methylation intervals extended broadly into structurally complex and non-reference sequence, with strong enrichment in insertion regions (16,804; 48.2% of those tested in these regions; 2.79-fold), particularly at young MEIs (3.22–3.64-fold), and enrichment in segmental duplications (118,790 intervals; 1.39–1.77-fold) and CHM13 sequence absent from GRCh38 (11,660; 1.44-fold); 116,563 significant intervals also mapped within CenSat sequence (Supplementary Table 15). Selecting the most strongly associated variants for each significant feature yielded 1,288,781 mQTL lead variants, 15,770 isoform QTL (isoQTL) lead variants and 13,892 splicing QTL (sQTL) lead variants. When QTLs were summarized by gene, 6,441 genes had a promoter mQTL, 7,374 an isoQTL and 8,673 an sQTL, with 376 genes having all three types (**Fig. 5e**). Lead variants were closest to methylation features: the median distance was 3,128 bp to a methylation interval, 22,219 bp to an isoform transcription start site and 14,302 bp to the midpoint of a splice-junction interval (**Extended Data Fig. 9a**). For 461,607 of the 1,641,874 significant methylation intervals (28.11%), the mQTL lead variant fell within the associated interval itself. Array-based mQTL studies can miss a subset of such proximal genetic effects because probes with variants at the assayed CpG or single-base extension site are typically excluded^8^.

To resolve candidate causal variants, we performed fine-mapping^44^. Allele-aware fine-mapping identified 1,975,769 credible sets for 1,496,452 methylation intervals, 19,003 credible sets for 13,248 isoforms and 23,269 credible sets for 14,327 splice junctions (**Fig. 5f,g**). Median credible-set sizes were three variants for mQTLs, five for isoQTLs and three for sQTLs. Of these, 628,540 mQTL, 5,114 isoQTL and 7,775 sQTL credible sets contained a single variant; 51.50 to 60.06% of credible sets across the three QTL classes contained no more than five variants.

We assessed the functional context of fine-mapped variants through enrichment across functional annotations and genomic features. Fine-mapped mQTLs and isoQTLs were both enriched for variants with coding consequences, with stronger enrichment for isoQTLs (1.36-fold and 4.62-fold, respectively), whereas fine-mapped sQTLs were enriched for variants with splice-donor-region consequences (4.71-fold; **Extended Data Fig. 9b**), consistent with previous studies of genetic regulation of splicing^45^.

Chromatin-state enrichments also differed among QTL classes. In the GM12878 15-state ChromHMM annotation (Roadmap E116)^46^, fine-mapped mQTLs were enriched in Polycomb-repressed and bivalent chromatin and depleted from strongly transcribed chromatin, whereas fine-mapped isoQTLs were enriched in transcribed chromatin and genic enhancers. Fine-mapped sQTLs were most enriched at active transcription start sites and their flanking regions (**Extended Data Fig. 9c**). Among the CGI trajectory classes defined above, fine-mapped sQTLs were enriched at recurrent-loss CGI loci (2.66-fold; FDR = 0.0233; **Fig. 6a**).

**Fig. 6.**
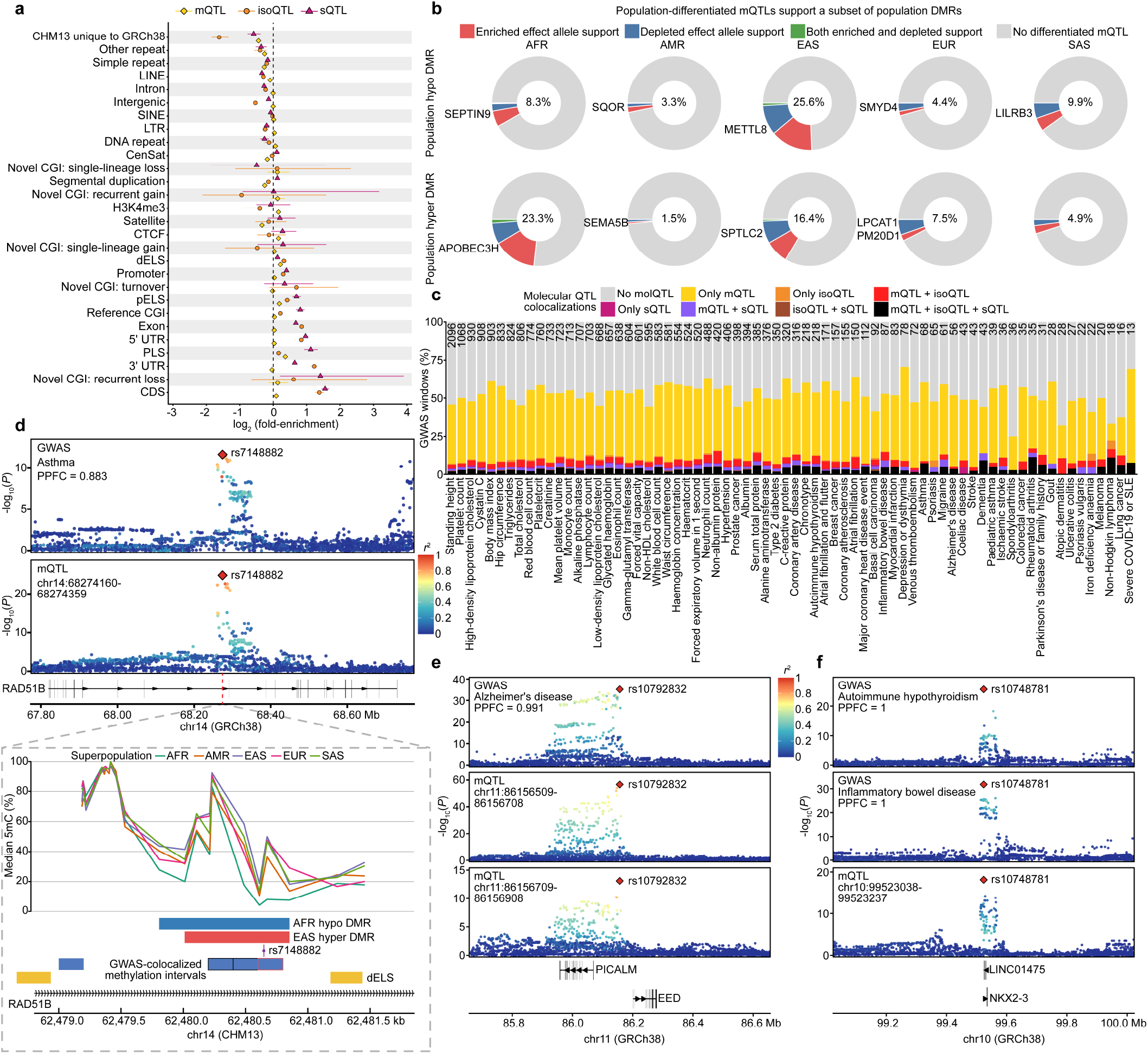
Functional, population and complex-trait contexts of fine-mapped molecular quantitative trait loci (QTLs). a, Empirical enrichment of the variant with the highest posterior inclusion probability (PIP) from each independent methylation QTL (mQTL), isoform QTL (isoQTL) or splicing QTL (sQTL) credible set across genomic features. Yellow diamonds, orange circles and magenta triangles denote mQTLs, isoQTLs and sQTLs, respectively. Points show log₂ observed-to-expected overlap; horizontal lines are empirical 95% intervals from 1,000 control sets matched for allele frequency, distance to the molecular feature, variant class and, for structural variants, length. The dashed vertical line denotes no enrichment. The labels denote centromere/satellite repeats (CenSat), long interspersed nuclear element (LINE), short interspersed nuclear element (SINE), long terminal repeat (LTR), promoter-like signature (PLS), proximal enhancer-like signature (pELS), distal enhancer-like signature (dELS), untranslated region (UTR) and coding sequence (CDS). CpG-island (CGI) trajectory classes use the genealogy-based classification. b, Population-differentiated mQTL support for 217,637 population differentially methylated region (DMR) calls. Columns correspond to African (AFR), Admixed American (AMR), East Asian (EAS), European (EUR) and South Asian (SAS) superpopulations; the top and bottom rows show hypomethylated and hypermethylated population DMRs, respectively. Donuts partition DMR calls into support from an effect allele enriched in the focal superpopulation, an effect allele depleted from it, both, or no qualifying differentiated mQTL. Support required concordance between the methylation shift predicted from the population allele-frequency difference and mQTL effect and the observed DMR direction. Both denotes support for the same DMR call by at least one enriched and at least one depleted effect allele. In total, 17,580 population DMR calls (8.08%) were supported. Centres give the supported fraction and gene labels mark representative DMRs containing a transcription start site. c, Molecular-QTL colocalization outcomes for 26,237 1-Mb windows centred on independent lead variants across 70 genome-wide association study (GWAS) traits and successfully evaluated by HyPrColoc. Stacked bars show the fraction of analyzed windows with no molecular-QTL colocalization, colocalization involving one QTL class, each pair of classes or all three classes; numbers above the bars give the number of analyzed windows per trait. d, Asthma association at rs7148882 (HyPrColoc posterior probability of full colocalization (PPFC) = 0.883). The representative methylation-interval mQTL has β = −0.431 and *P* = 1.44 × 10⁻²⁵. The lower CHM13 view shows median methylation profiles for each superpopulation, the AFR-hypomethylated and EAS-hypermethylated DMRs, GWAS-colocalized methylation intervals, a dELS and *RAD51B*. Blue boxes mark the colocalized methylation intervals, and the red outline marks the representative interval plotted above. The mQTL effect allele is enriched in AFR (0.900 versus 0.206 in non-AFR haplotypes) and depleted in EAS (0.078 versus 0.514), concordant with its negative mQTL effect and reciprocal population-DMR directions (*n* = 363 haplotypes). e, Alzheimer’s disease association at rs10792832 (PPFC = 0.991), colocalizing with two adjacent methylation intervals (β = −0.635, *P* = 2.63 × 10⁻⁵⁷; β = −0.333, *P* = 7.96 × 10⁻¹⁴); *PICALM* and *EED* provide genomic context. f, Autoimmune hypothyroidism and inflammatory bowel disease associations at rs10748781. Each GWAS signal colocalizes with the same methylation interval (PPFC = 1.0 for each; β = 0.343, *P* = 7.30 × 10⁻¹⁹). In d–f, association panels use GRCh38, the coordinate system used to harmonize GWAS and molecular-QTL variants; lower methylation and genomic-feature tracks, where shown, use CHM13. Filled red diamonds mark the HyPrColoc candidate variant, point colour gives phased-haplotype *r*² to that variant, and mQTL β values are reported relative to the CHM13 alternate allele. Gene labels provide genomic context and do not assign molecular targets.

Finally, we tested whether QTLs affecting different molecular phenotypes at the same locus were consistent with the same underlying causal signal. We therefore performed pairwise colocalization among mQTLs, isoQTLs and sQTLs using coloc^47^, which distinguishes evidence for a shared causal variant from evidence for distinct nearby causal variants. Colocalized features most often involved methylation: 50,411 occurred only in methylation-isoform colocalizations and 26,404 only in methylation-splicing colocalizations, compared with 2,668 occurring only in isoform-splicing colocalizations (**Fig. 5h**). Across all pairwise comparisons, 106,115 molecular features participated in at least one colocalized signal. At the variant level, 272,663 variants were included in at least one 95% credible set for a shared molecular-QTL signal, including 38,715 represented across all three QTL classes (**Fig. 5h**). The shared-signal credible sets were dominated by small variants, whereas 1,259 were SVs.

### Fine-mapped mQTLs link population methylation differences to complex-trait loci

We next asked whether mQTLs could link population DMRs to complex-trait associations. We first used the fine-mapped mQTL map to quantify how much of the population methylation difference is under cis genetic control. We defined a population-differentiated mQTL as one whose effect allele differed in frequency by at least 0.1 and was at least twofold enriched or depleted in the focal superpopulation relative to all remaining haplotypes. To link such an mQTL to a population DMR, we further required the methylation shift predicted from the allele-frequency difference and mQTL effect to agree with the observed methylation change direction across the DMR (Methods). An enriched methylation-increasing allele and a depleted methylation-decreasing allele therefore both supported a hypermethylated DMR, whereas the reverse combinations supported a hypomethylated DMR. Of 217,637 population DMR calls, 17,580 (8.08%) were supported by population-differentiated mQTLs: 10,693 by enriched effect alleles, 6,281 by depleted effect alleles and 606 by both enriched and depleted supporting alleles (**Fig. 6b**). The supported fraction ranged from 3.3% in AMR to 25.6% in EAS among hypomethylated DMRs and from 1.5% in AMR to 23.3% in AFR among hypermethylated DMRs.

Among these genetically supported DMRs, 967 contained at least one transcription start site, encompassing 2,762 transcripts from 882 genes; 854 of these transcripts were novel isoforms. For example, at the *LPCAT1* promoter, median methylation of the EUR-hypermethylated DMR was 30.2% in EUR compared with 12.2 to 18.5% in the other superpopulations, and the methylation-increasing allele at rs36978 was 3.7-fold more frequent in EUR than in non-EUR haplotypes (**Extended Data Fig. 9d,e**). The DMR at the *LPCAT1* promoter was spanned by eight consecutive methylation intervals and encompassed the transcription start sites of five novel isoforms. Pairwise molecular-QTL colocalization at this locus linked 29 methylation intervals, including the eight spanning the DMR, to two isoforms and two splice junctions in a shared signal, with rs36978 consistently identified as the shared leading variant (**Extended Data Fig. 9f** and **Supplementary Fig. 13**). Both isoQTLs showed lower isoform abundance with the methylation-increasing allele, while junction usage shifted reciprocally between two alternative junctions of the same splicing cluster (**Extended Data Fig. 9f** and **Supplementary Fig. 14**). Thus, a single shared genetic signal at *LPCAT1* was associated with coordinated differences across all three molecular phenotypes.

The AFR-hypermethylated *APOBEC3H* promoter DMR illustrated a different architecture, with more than one fine-mapped mQTL signal. Median DMR methylation was 60.8% in AFR compared with 35.0 to 42.7% in the other superpopulations, and the DMR encompassed the transcription start sites of 14 transcripts, eight of them novel (**Extended Data Fig. 9g,h**). Two methylation-increasing alleles, rs41281251 and rs55989856, each the top fine-mapped variant for a separate methylation interval, were 6.7- and 7.0-fold more frequent in AFR than in non-AFR haplotypes (**Supplementary Fig. 15**). Together, *LPCAT1* and *APOBEC3H* illustrate how population methylation differences can arise from either a shared multi-phenotype genetic signal or multiple local mQTL signals.

We next extended the molecular-QTL analysis to complex-trait associations. We examined 26,237 1-Mb windows centered on independent lead variants from genome-wide association studies (GWAS) across 70 diseases and quantitative traits from the GWAS Catalog, Pan-UK Biobank and FinnGen^48–50^ (Supplementary Table 16). Previous long-read studies have used molecular-QTL colocalization to resolve disease-associated isoforms and splicing events^40,51^. We used HyPrColoc^52^ to test whether GWAS and molecular-QTL associations within each window were consistent with a shared causal variant. More than half of the GWAS windows (13,894 of 26,237; 52.96%) colocalized with at least one molecular QTL (**Fig. 6c**). Methylation provided the broadest molecular annotation of colocalized trait loci: mQTLs were present in 13,533 of 13,894 windows (97.40%), including 11,307 (83.55%) with no isoQTL or sQTL colocalization (Supplementary Table 17).

These colocalizations included major disease loci. At the *PICALM-EED* locus, the Alzheimer’s disease association at rs10792832^53^ colocalized with mQTLs for two adjacent methylation intervals, with a posterior probability of full colocalization (PPFC) of 0.991 (**Fig. 6e**). At *LINC01475-NKX2-3*, the GWAS associations for autoimmune hypothyroidism and inflammatory bowel disease^50^ both colocalized with the same mQTL signal for a single methylation interval at rs10748781 (β = 0.343; *P* = 7.30 × 10^−19^; PPFC = 1.0 for each; **Fig. 6f**). A single methylation-regulatory signal was therefore shared across two related immune disease associations.

Population allele-frequency differences further connected these trait-associated mQTLs to the population DMRs described earlier. At rs7148882, the asthma GWAS association^54^ colocalized with an mQTL (PPFC = 0.883) whose effect allele was strongly enriched in AFR (frequency 0.900 versus 0.206 in non-AFR haplotypes) but depleted in EAS (0.078 versus 0.514 in non-EAS haplotypes). The allele decreased methylation (β = −0.431), matching the reciprocal population pattern of an AFR-hypomethylated DMR and an EAS-hypermethylated DMR (**Fig. 6d**). A similar relationship was found at rs2710323, where the body mass index GWAS association^55^ colocalized with an mQTL (PPFC = 0.999). The methylation-decreasing effect allele (β = −0.362) had frequencies of 0.743 in AFR and 0.236 in SAS, concordant with the AFR-hypomethylated and SAS-hypermethylated DMRs at the *ITIH* locus (**Supplementary Fig. 16**).

We systematically quantified how often population-differentiated mQTL variants were also represented in GWAS-colocalized signals across all 70 traits. Across the five superpopulations, we identified 17,990 unique population-differentiated mQTL variants; 202 (1.12%) were also identified by HyPrColoc as the candidate shared variant for at least one GWAS signal across 48 traits (**Extended Data Fig. 10a**). At the GWAS-window level, 277 of the 13,533 GWAS windows with an mQTL colocalization (2.05%) contained the same variant-methylation interval pair that supported a population DMR. Of these windows, 105 were supported by an allele enriched in the focal superpopulation, 120 by a depleted allele and 52 by both enriched and depleted supporting alleles (**Extended Data Fig. 10b**). The matches spanned hematological, metabolic, cardiovascular and immune phenotypes, including platelet traits, hypertension, glycated hemoglobin and monocyte count (**Extended Data Fig. 10c–f**). Together, these results identify a defined set of loci at which population allele-frequency differences, methylation divergence and complex-trait associations converge on the same genetic signal.

## Discussion

Human genomics has moved from a single linear reference to complete genomes and pangenomes that explicitly represent sequence diversity^56,57^. Our results extend this transition from genome representation to epigenomic analysis. The central problem is no longer simply whether non-reference sequence can be measured^29,30^, but how epigenomic states can be compared across homologous genomes with different sequence and CpG content. By defining regional homology without requiring identical sequence content, our projection framework allows each haplotype to contribute the sequence and CpGs it carries within a common population-scale comparison. This comparison revealed that CpG-density variability and methylation variability occupied largely distinct genomic regions: genetic variation can alter CpG content without proportionally increasing methylation variability. A pangenome view therefore changes both the observable genomic space and what constitutes a comparable methylation phenotype.

Comparative epigenomic studies suggested that epigenetic states can remain conserved despite sequence divergence^58,59^, but whether such buffering operates genome-wide within humans remained untested. Here we provide the first direct genome-wide quantification of this relationship across hundreds of diverse human haplotypes, revealing that methylation variability did not increase with local genetic diversity across more than 90% of the genome, whereas the buffering effect broke down only at extreme nucleotide diversity. SVs nevertheless created localized departures from this genome-wide stability. Previous studies documented DNA methylation changes associated with SV breakpoints and polymorphic TE insertions^20,21,28^, but did not define the factors governing their occurrence and magnitude. Variant subtype, repeat composition and the genomic context of the SV determined DMR occurrence and count, whereas DMR magnitude depended mainly on the genomic context of the DMR. Thus, population-scale analysis moves the field from documenting SV-associated methylation to defining the rules governing when, where and how sequence reconfiguration perturbs the methylome.

CGIs reveal a second consequence of sequence variation: genetic variants can change not only how a CGI is methylated, but also whether the CGI itself exists. CGIs are sequence-defined, yet human methylome analyses generally rely on reference-genome CGI annotations, whereas CGI gain and loss have largely been examined across species^4,5^. PanCGI resolves CGI variation across human haplotypes into homologous loci and sequence-resolved alleles, making it possible to directly genotype CGI states. SVs and even a few sequence changes can create or remove a CGI. Local genealogies further place these polymorphisms in evolutionary context, reconstructing ancestral states and CGI gain, loss and turnover within modern humans.

Comparative studies linked CGI turnover between species to regulatory evolution^5^; our results extend this paradigm within humans, where loci undergoing CGI gain, loss or turnover are concentrated in regulatory and transcribed sequence. Together, these findings bring CGI evolution into human population genetics, establishing CGI presence as a sequence-resolved polymorphic epigenetic feature rather than a fixed reference annotation.

Pangenome representation becomes especially consequential for association genetics. Long-read and pangenome studies have made non-reference methylation measurable, but population-scale methylation phenotypes remain largely confined to reference-defined genomic space^29,30^. By coupling homologous regional phenotypes to allele-aware QTL mapping, our framework brings methylation variation within non-reference sequence itself into cis-mQTL analysis. Methylation intervals under cis genetic control were disproportionately enriched in these newly accessible contexts, especially insertion regions, including young MEIs, and other structurally complex regions. The resulting mQTL map added a broad methylation-based molecular layer to complex-trait loci, with mQTLs colocalizing with more than half of tested GWAS windows. Fine-mapped population-differentiated mQTLs provided a high-resolution quantitative assessment of cis genetic support for population methylation differences. Strikingly, these mQTLs also colocalized with complex-trait associations, linking population genetic differentiation and methylation divergence to trait-associated genetic architecture. Together with genome-wide epigenetic buffering, these results resolve genetic-epigenetic interaction at two complementary scales: broad methylation robustness to genetic divergence and precise local cis control where coupling occurs.

The current cohort size cannot fully represent global human genetic diversity, and lymphoblastoid cell lines capture a limited cellular context. A key next question is how sequence-resolved CGI alleles shape context-specific methylation and regulatory function across tissues, developmental stages and disease states. Larger pangenome-based methylomes and perturbation studies will test how broadly the genetic-epigenetic interactions resolved here extend across human biology. More broadly, pangenomes should change not only where functional data are aligned, but how epigenomic phenotypes themselves are defined. This work therefore establishes a general strategy for population epigenomics and recasts the human methylome as a polymorphic regulatory layer that is genetically robust yet locally responsive, accessible to population genetics, evolutionary reconstruction and complex-trait interpretation.

## Methods

### Study samples and population definitions

Analyses used 230 HPRC2 individuals with matched diploid assemblies and Oxford Nanopore methylation data (460 haplotypes; 115 female and 115 male)^29^. Superpopulation assignments followed HPRC2 metadata and the 1000 Genomes Project Phase 3 classification: 70 individuals were assigned to AFR, 44 to AMR, 51 to EAS, 29 to EUR and 36 to SAS^31^. HG06807 and NA21309 were included in AFR and HG005 in EAS (Supplementary Table 1).

Analyses requiring matched methylation data used the 230 HPRC2 individuals described above (460 haplotypes). PanCGI was constructed from 462 HPRC2 haplotype assemblies representing 231 individuals, including HG002, together with T2T-CHM13 and GRCh38. Transcript discovery used 202 individuals, and isoQTL and sQTL analyses used 201 after excluding HG00272 because of chromosome X data quality.

### Projection of CHM13 intervals to haplotype assemblies

We partitioned the autosomal T2T-CHM13 v2.0 genome^56^ into non-overlapping 200-bp intervals. Each CHM13 interval was projected independently onto each of the 460 assemblies using halLiftover (v.2.2)^60^ and the Minigraph-Cactus hierarchical alignment^61^. We used --bedType 4 --outPSLWithName to retain the CHM13 interval identifier in the PSL output. The sequence recovered by projection on each haplotype was termed the haplotype-specific interval. A CHM13 interval can project onto sequences longer or shorter than its own length, so the corresponding haplotype-specific interval can be longer or shorter than 200 bp, or absent. When a CHM13 interval yielded more than one alignment record on a haplotype, the records were pooled: projected lengths were summed, CpGs from all records were combined, and the record count was retained as a multiple-mapping flag.

Masks were applied after projection. We removed intervals on chromosomes X and Y and on the mitochondrial genome, and intervals overlapping the complex satellite annotation, the telomeric regions, or regions defined in chm13v2.cen-mask.bed^62^. The chm13v2.cen-mask.bed mask covers centromeric satellite repeats and acrocentric short arms. It was curated from the T2T-CHM13 satellite annotation, DNA-BRNN satellite calls and the HPRC year-1 minigraph graph, and identifies regions in which alignment is substantially less reliable than elsewhere in the genome.

### Interval-level methylation and CpG density

For every haplotype-specific interval we recorded its length, its CpG content and its methylation. Only 5-methylcytosine calls at CpG dinucleotides were used to quantify methylation. CpG methylation measurements were retained only when read coverage on the corresponding haplotype was at least five; measurements below this threshold were set to missing. Interval methylation was the mean methylation percentage across CpGs with retained measurements in the interval.

CpG density was the number of detected CpGs divided by the projected interval length in kilobases. Because coverage determines the reliability of a methylation estimate and not whether a CpG is present, no coverage threshold was applied to CpG density. Methylation and CpG density were measured on the same interval using distinct inclusion criteria.

### Graph coordinates for CpG dinucleotides

Projected intervals allow methylation to be compared across homologous regions by averaging methylation over the sequence carried by each haplotype. For analyses requiring single-CpG resolution, we mapped CpG dinucleotides to pangenome graph coordinates.

CpG sites were identified in the linear coordinates of each assembly and mapped to the corresponding haplotype path in the Minigraph-Cactus graph. Each site was assigned a graph identifier comprising the graph segment and the offset within that segment^63,64^. Homologous CpGs on different haplotypes were assigned the same identifier, whether or not the surrounding sequence was present in CHM13. CpGs in non-reference sequence were also assigned graph identifiers.

For each haplotype, we removed CpGs overlapping the union of the assembly-specific satellite and active centromere annotations. Graph identifiers were used for population differential methylation, SV differential methylation and the CGI analyses.

### CpG discovery and core curves

Assembly-based CpG discovery and core curves were computed by sequentially adding the 460 haplotypes in their input order. At each step, the discovery curve recorded the number of distinct graph CpGs across the included haplotypes, and the core curve recorded the number shared by all included haplotypes. CpGs observed on only one of the 460 haplotypes (singletons) were excluded from these curves.

Across the 460 haplotypes, we identified 35,200,252 distinct CpG sites, of which 17,933,898 (50.9%) were present on every haplotype. CHM13 contained 33,839,747 CpG sites.

### Covariate-adjusted variance and scaled variability

The unit of analysis was defined by a 200-bp CHM13 interval and its corresponding haplotype-specific projections. Interval methylation was arcsine-transformed as *y* = arcsin(2*m* − 1), where *m* is mean methylation in the haplotype-specific interval. For each interval, we fitted an ordinary least-squares model across haplotypes with methylation values for that interval. The design matrix contained sequencing chemistry, sex, the z-score of log₂ mean coverage, the z-score of read N50 and 30 methylation factors estimated using probabilistic estimation of expression residuals (PEER; v.1.3)^65^. Residual variance was estimated from this model, and the interval mean was calculated from the transformed methylation values. Intervals with methylation values for at least 20% of haplotypes were included in the analysis, and the subset used for mean-variance fitting required methylation values for at least 50% of haplotypes.

Both methylation and CpG density show a strong mean-variance relationship, so a single genome-wide variance threshold would reflect this mean dependence rather than identify intervals with greater variance than expected at a given mean. We therefore modelled the expected residual variance as a smooth function of the interval mean using mgcv (v.1.9-4)^66^ with bam(log(s2) ∼ s(mu, k = 30, bs = “cs”), method = “fREML”, discrete = TRUE), then back-transformed the fitted values to the variance scale. Scaled variability was the ratio of observed to expected variance. A value of one denotes covariate-adjusted variance equal to the expectation at the same mean, and values above one denote greater variance than expected. CpG-density variability was calculated in the same way using a separate mean-variance model with k = 20.

This mean-variance correction follows established approaches for calling highly variable features in single-cell transcriptomics^67^ and hypervariable signals in chromatin profiling^68^.

### Empirical null estimation and identification of HVMRs and HVDRs

HVMRs and HVDRs were identified separately. Of the 13,460,232 tested autosomal intervals, 9,742,607 had methylation values and 9,749,472 had CpG-density values for at least 20% of haplotypes. Requiring methylation or CpG-density values for at least half of the haplotypes retained 9,483,032 intervals for methylation and 9,575,790 intervals for CpG density; these sets were used for mean-variance fitting and identification of HVMRs and HVDRs, respectively.

For each measure, the raw observed-to-expected variance ratio was evaluated against a robust scaled-*F* empirical null distribution. Each interval was assigned to a single annotation group according to a prespecified order of precedence. For methylation, the 15 groups were, in order of precedence, RefSeq promoters, masked CGIs, their shores and their shelves, ENCODE candidate cis-regulatory elements, SVs and MEIs, seven RepeatMasker classes (DNA, long interspersed nuclear element (LINE), low complexity, long terminal repeat (LTR), satellite, simple repeat and short interspersed nuclear element (SINE)), other repeats, and CGI open sea. For CpG density, the 14 groups replaced RefSeq promoters and ENCODE elements with segmental duplications and were otherwise identical. Intervals were stratified jointly by annotation group and by ten quantile bins of the interval mean. Quantile boundaries were calculated within each annotation group for methylation and genome-wide for CpG density. Within each stratum, intervals at or below the 70th percentile of the variance-ratio distribution were eligible for sampling. Sampling was balanced across chromosomes and capped at 30,000 intervals per stratum. The methylation null comprised 2,825,863 intervals.

We estimated the scale parameter and prior degrees of freedom of the scaled-*F* distribution using limma (v.3.66.0) fitFDistRobustly^69,70^ with lower- and upper-tail winsorization probabilities of 0.05 and 0.10. Upper-tail *P* values were adjusted genome-wide using the Benjamini– Hochberg procedure, and HVMRs and HVDRs were identified at an FDR of 0.05.

Genome-browser tracks show the log₂ observed-to-expected ratio, and 1-Mb display windows show the median of the underlying interval ratios on the same log₂ scale.

### Parent-of-origin DMRs

Maternal and paternal labels were known for 127 of the 230 individuals. Assemblies for the remaining 103 individuals were Hi-C-phased. The hap1 and hap2 labels were consistent within each chromosome, but their correspondence to maternal and paternal origin could differ between chromosomes. We inferred parental origin from methylation at imprinted regions before analyses requiring maternal and paternal labels.

Parent-of-origin DMRs were identified in the 127 individuals with known labels. At each CpG, we arcsine-transformed methylation and used the within-individual paternal-minus-maternal difference as the response. Paired methylation values were included only when both haplotypes had read coverage of at least five, and paired differences were weighted by the harmonic combination of the two read coverages. CpGs with paired methylation values from at least 20 individuals were analyzed with an intercept-only limma (v.3.66.0) model^69^ using empirical-Bayes moderation with robust = TRUE. Regions were called from the site-level statistics using callDMR in DSS (v.2.58.0)^71^ at a per-CpG *P* threshold of 0.01, with other region criteria at their defaults. We identified 2,121 autosomal regions.

### Marker selection and inference of parental origin

We lifted published imprinted regions^72,73^ to CHM13 coordinates and used their annotated methylation directions to select markers from the parent-of-origin DMRs identified in our cohort. A region was retained when at least one overlapping published annotation agreed in methylation direction and at least 80% of overlapping annotations were concordant. Markers also required an absolute mean paired methylation difference of at least 15%, a consistent direction of difference in at least 75% of informative individuals, at least 20 informative individuals and at least three CpGs. These criteria identified 118 high-confidence parent-of-origin methylation markers across the 22 autosomes.

For each of the 103 individuals, we inferred parental origin independently on each autosome (2,266 comparisons in total). Within each marker, methylation was averaged across the observed 200-bp intervals separately for hap1 and hap2. We calculated hap1-minus-hap2 methylation differences, reversing the sign for markers with higher maternal methylation so that positive values supported hap1 as paternal. The chromosome-level score was the weighted mean of the marker differences. Marker weights were estimated from the 127 individuals with known parental origin and then fixed. Each weight was the product of the absolute mean paternal-minus-maternal methylation difference, directional consistency, the log-transformed CpG count ln(1 + *n*_CpG), and an evidence multiplier. Evidence multipliers were three for markers supported by whole-genome bisulfite sequencing data or previously known parent-of-origin regions, two for novel parent-of-origin regions and one for array-only support.

Parental origin was assigned when at least one marker had methylation values for both haplotypes, the absolute weighted score was at least 0.05, and at least 65% of the total informative marker weight supported one parental orientation. The sign of the weighted score determined the parental assignment. In total, 2,092 of the 2,266 comparisons (92.3%) met these criteria, and the corresponding haplotypes were relabelled as maternal or paternal. Of the remaining 174, 69 lacked marker methylation values for both haplotypes, 82 had an absolute weighted score below 0.05 and 23 did not meet the weighted directional concordance threshold. For these chromosomes, values in the maternal and paternal columns of the methylation matrix were set to missing. Known maternal and paternal labels were retained, and the same assignments were applied to the batch-corrected matrix.

### Batch correction of interval methylation

The methylation matrix for the 200-bp CHM13 intervals was batch-corrected using ComBat-met (v.0.99.3)^74^, with sequencing chemistry as the batch variable while preserving methylation variation associated with sex and the first ten genetic principal components. An interval was corrected only when methylation values were available for at least ten haplotypes in each sequencing chemistry group. The corrected matrix was used for haplotype clustering and SV analyses based on projected intervals. For HVMR and HVDR identification, the per-interval regression models included sequencing chemistry, sex, the z-score of log₂ mean coverage, the z-score of read N50 and 30 methylation PEER factors.

### Haplotype clustering

We clustered maternal and paternal haplotypes using batch-corrected methylation across the 24,022 HVMRs. Methylation values were arcsine-transformed as *y* = arcsin(2*m* − 1), and intervals with methylation values for fewer than two haplotypes were removed. Because sequence presence and absence vary among haplotypes, imputing missing methylation values before principal-component analysis could cause structural differences to influence the embedding. We therefore calculated pairwise Euclidean distances with daisy in cluster (v.2.1.8.2)^75^, using intervals with methylation values in both haplotypes of each pair.

We performed principal coordinate analysis of the distance matrix with the Lingoes correction using pcoa in ape (v.5.8-1)^76^. The first 20 corrected axes were used as input to UMAP, implemented with umap in uwot (v.0.2.4)^77^ using Euclidean distances, 50 neighbours and a minimum distance of 0.2.

### Population differences in methylation

We tested population differences in methylation at CHM13 reference CpGs, with methylation values for each haplotype retrieved by exact matches to graph CpG identifiers. We compared haplotypes from each of the five superpopulations with all remaining haplotypes. Only 5-methylcytosine calls with read coverage of at least five were retained, and a CpG was tested when methylation values were available for at least 20 haplotypes in the focal superpopulation and at least 20 in the remaining cohort.

We analyzed differential methylation using two DSS (v.2.58.0)^71^ models, both with a smoothing span of 500 bp. A two-group model estimated mean methylation in each group and the direction of the difference. A multifactor model including group and sequencing chemistry as covariates provided the *P* values used to call DMRs.

Population DMRs were called at a per-CpG *P-value* threshold of 1 × 10^−3^, with remaining parameters set at defaults. Methylation differences were calculated from the two-group model as mean methylation in the focal superpopulation minus mean methylation in the remaining cohort.

### Genomic annotation sets

We compiled 91 genomic annotation sets in seven categories. Gene features were obtained from the CHM13 RefSeq annotation and comprised eight sets: promoters, 5′ UTRs, coding sequence, exons, introns, 3′ UTRs, gene bodies and intergenic sequence. The CHM13 repeat annotation^78,79^ provided 17 repeat classes. ENCODE candidate cis-regulatory elements were lifted from GRCh38 to CHM13 coordinates^80^ and classified into nine groups comprising promoter-like, proximal enhancer-like, distal enhancer-like, DNase-H3K4me3 and CTCF-only elements, distinguished by CTCF-binding status. CGI annotations comprised islands called after repeat masking, their shores and shelves, and the intervening open sea. Shores extended 2 kb on each side of an island, and shelves covered the next 2 kb. Segmental duplications were classified as pericentromeric, telomeric or acrocentric. The remaining 50 sets comprised regions occupied by variants in the HPRC2 SV call set^29^ with minor allele frequency above 0.005. Insertions, deletions and inversions defined three sets, and a further 47 were defined by mobile-element family and insertion or deletion status relative to CHM13. For mQTL enrichment, the 52 genomic annotation sets used in **Fig. 1d** were supplemented with CHM13 sequence absent from GRCh38 and CenSat sequence.

### Genomic enrichment

Three enrichment analyses were used for sets of 200-bp intervals, population DMRs and CGI loci, respectively.

HVMRs, HVDRs and intervals shared between the two sets were tested for over-representation on the 200-bp analysis grid using the compareCluster and enricher functions in clusterProfiler (v.4.18.4)^81^. For the analysis shown in **Fig. 1d**, 52 retained annotation sets were used. Of the 13,460,232 masked autosomal intervals supplied as the background, 10,880,074 overlapped at least one of these annotation sets and formed the effective background. Of the 24,022 HVMRs, 22,106 overlapped at least one retained annotation set. One-sided hypergeometric *P* values were adjusted across annotation sets within each query using the Benjamini–Hochberg procedure^82^. Enrichment of methylation intervals with significant cis-mQTLs was tested with enricher using all intervals tested for mQTLs as the supplied background, with annotation sets restricted to those containing 50–4,000,000 tested intervals (52 sets).

To assess parent-of-origin enrichment, we added the 1,879 autosomal intervals overlapping the high-confidence parent-of-origin methylation markers as a 53rd annotation set for HVMR and HVDR enrichment. The effective background for this analysis comprised 10,880,268 intervals, of which 22,152 were HVMRs.

Enrichment of population DMRs was evaluated using their genomic coordinates and lengths, with two separate methods. Testing was restricted to CHM13 positions tested for differential methylation on chromosomes 1–22, stratified by GC-content octiles in 10-kb windows. Query sets comprised population DMRs called at a per-CpG *P* threshold of 1 × 10^−3^. We performed 1,000 GC-stratified interval randomizations using regioneR (v.1.42.0)^83^. Genomic Association Tester (GAT; v.1.3.6)^84^ was run with 10,000 samples and nucleotide overlap as the test statistic. For each query set and each method, *P* values were adjusted across the 91 annotation sets. An annotation set was retained only when both methods yielded *q* ≤ 0.05 and concordant enrichment or depletion. GAT log₂ fold enrichment and *q* values were reported.

Enrichment of CGI loci was evaluated separately for five evolutionary trajectories: single-lineage loss, single-lineage gain, recurrent loss, recurrent gain and turnover. We used GAT with 10,000 samples and nucleotide overlap as the test statistic, including chromosomes 1–22, X and Y and stratifying by GC-content octiles in 10-kb windows.

### Sequence diversity measures

We calculated SNP π with VCFtools (v.0.1.17)^85^ from small variants in the HPRC2 PanGenie variant set^86^. This set comprises variants from the HPRC2 Minigraph-Cactus graph that were decomposed into biallelic records and normalized using the PanGenie pipeline, with CHM13 as the reference. Variants were restricted to biallelic SNPs using --min-alleles 2 --max-alleles 2 --remove-indels. Nucleotide diversity was estimated in non-overlapping 200-bp windows using --window-pi 200 --window-pi-step 200, with 1-Mb estimates generated for visualization.

We calculated synteny diversity across the haplotype assemblies with SynDiv (v.1.1.5)^87^, which uses MUMmer (v.4.0.1)^88^ for assembly alignment. Centromeric and pericentromeric sequence was excluded. Values were summarized in non-overlapping 200-bp and 1-Mb CHM13 intervals.

### Generalized additive models of methylation variability

We modelled scaled methylation variability as a function of CpG-density variability, SNP π and synteny diversity. The response was log₂-transformed scaled methylation variability.

CpG-density variability was zero in a substantial fraction of intervals. We represented CpG-density variability with an indicator distinguishing zero from non-zero values and a cubic regression spline fitted to non-zero values. Positive values were winsorized at the 0.1st and 99.9th percentiles. SNP π was transformed as log1p(π / median positive π) and modelled with a single smooth term because zero values were rare. Synteny diversity was also represented by an indicator distinguishing zero from non-zero values and a spline fitted to non-zero values. Mean methylation and mean CpG density were included as smooth covariate terms, and chromosome was included as a random effect.

We fitted the model using the bam function in mgcv (v.1.9-4)^66^ with a Gaussian family, cubic regression splines, fast restricted maximum likelihood (fREML), discrete fitting and term selection. Basis dimensions were 96 for CpG-density variability, 16 for SNP π, 8 for synteny diversity and 20 each for mean methylation and mean CpG density. Observations were weighted by the precision of the interval variance estimate, with weights truncated to the range 0.5–2. Weighted *R*² was one minus the weighted residual sum of squares divided by the weighted total sum of squares.

The contribution of each predictor was quantified as the difference in weighted *R*² between the full model and a nested model omitting that predictor. Differences in weighted *R*² do not indicate the direction of an association. The direction of each association was assessed separately by partial Spearman correlation and by partial-residual curves after adjustment for the remaining model terms. Uncertainty was estimated using chromosome-stratified block bootstrap. Each chromosome was divided into non-overlapping 5-Mb blocks, which were resampled with replacement within each chromosome. Weighted *R*² and the contribution of each predictor were recalculated in each of 1,000 bootstrap replicates. Models fitted within individual genomic annotation categories used the same transformations and were retained when at least 1,000 eligible intervals were available.

### SVs and quality control of flanking regions

SVs were obtained from the HPRC2 SV call set; 116,358 insertions and 84,429 deletions passed a minor allele frequency threshold of 0.005. We compared methylation between carrier and non-carrier haplotypes at T2T-CHM13 reference CpGs within 10 kb of either breakpoint. Because inserted and deleted sequence is present in only one of the two groups, we selected flanking CpGs by their CHM13 coordinates and retrieved methylation values for each haplotype by exact matches to graph CpG identifiers. This allowed homologous CpGs to be compared across haplotypes irrespective of their assembly coordinates.

The consensus breakpoint coordinates in the HPRC2 SV call set can differ from those of individual carrier haplotypes, shifting the 10-kb flanking windows and compromising the comparison of homologous flanking sequence. We therefore used each carrier haplotype’s own breakpoint coordinates and required at least 80% overlap between its 10-kb flanks and the corresponding flanks defined by the HPRC2 SV call set on both sides. A variant was retained when at least 80% of its carrier haplotypes met this criterion. All 3,315,813 flanks from insertion-carrying haplotypes passed the overlap criterion, whereas 9,424 of 3,348,216 flanks failed for deletions, excluding 876 deletions and retaining 83,553.

Carrier and non-carrier groups were defined from genotypes in the HPRC2 SV call set. CpG methylation values were retained only when read coverage was at least five. A CpG was tested when methylation values were available for at least three carrier and three non-carrier haplotypes. Requiring at least three eligible haplotypes per group and available reference CpGs retained 111,804 insertions and 80,760 deletions. Further requiring at least three tested CpGs and a completed DSS model fit adjusted for sequencing chemistry retained 107,040 insertions and 77,260 deletions, which were used in all subsequent models.

### Insertion- and deletion-associated differential methylation

We tested differential methylation at graph CpGs using two DSS (v.2.58.0)^71^ models with smoothing. A two-group model estimated the methylation difference at each CpG as mean methylation in carrier haplotypes minus mean methylation in non-carrier haplotypes. A multifactor model including carrier status and sequencing chemistry as covariates provided the *P* values used to call DMRs. DMRs were called at a per-CpG *P-value* threshold of 1 × 10^−3^, with remaining parameters set at defaults. DMR direction was determined from the sign of the DSS area statistic (areaStat). We tested 184,300 SVs and identified 76,977 SV-DMR pairs. DMRs were called separately for each SV, so the same genomic region could contribute to multiple SV-DMR pairs.

We summarized the distribution of DMRs relative to each breakpoint using 100-bp sliding windows with a 50-bp step size along the two 10-kb flanks, analysing each flank separately. A window was considered tested for a given SV if DSS returned a test result for at least one CpG within that window. At each window position, DMR burden was the fraction of tested SVs associated with an overlapping DMR of the specified direction.

### Outcome definitions and predictors

We defined three outcomes: DMR occurrence, whether an SV was associated with at least one DMR; DMR count, the number of DMRs associated with an SV; and DMR magnitude, the absolute methylation difference between carrier and non-carrier haplotypes for each SV-DMR pair. DMR occurrence and count were modelled at the SV level, and DMR magnitude at the SV-DMR pair level. Models were fitted separately for insertions and deletions, with separate analyses of all DMRs, hypermethylated DMRs and hypomethylated DMRs.

We considered five groups of predictors: variant subtype, repeat composition, SV length, the genomic context of insertion breakpoints and the genomic content of deleted sequence. Allele frequency and the number of CpGs tested in the flanks were included as covariates to account for differences in ascertainment. SV length and the number of tested flanking CpGs were log-transformed and standardized, and genomic-context terms were represented by indicator variables. Models of DMR magnitude included four additional groups of predictors: the genomic context of the DMR, its CpG count, its length and its distance to the nearest SV breakpoint. The three continuous DMR predictors were also log-transformed and standardized.

Variant subtypes were obtained from the annotations in the HPRC2 SV call set^29^. Subtype and repeat composition were annotated for individual SV events in each carrier haplotype, whereas models were fitted to merged SVs. A merged SV was assigned a subtype or repeat class when at least 80% of its carrier haplotypes shared that annotation. When agreement was below 80%, the distinct annotations from the individual SVs contributing to the merged SV were combined into a single label.

Three alternative SV classifications were evaluated in separate models: variant subtype, tandem-repeat class and the presence of TE sequence. A class was analyzed only when it included at least 30 SVs, or 100 for tandem-repeat classes, with at least 20 SVs associated with a DMR and at least 20 without a DMR. Genomic context was defined using the annotation sets described above. Genomic-context terms were analyzed when at least 80 SVs had the annotation and at least 80 did not. The corresponding threshold for DMR-level annotations was 50 SV-DMR pairs with the annotation and 50 without.

### Model fitting and contribution decomposition

We modelled DMR occurrence using logistic regression, DMR count using negative binomial regression with the logarithm of the number of tested 200-bp windows as an offset, and DMR magnitude using linear regression weighted by the number of CpGs in each DMR. For this offset, we divided the 10-kb flanks into 200-bp windows relative to each SV breakpoint: both sides of the insertion site for insertions, and upstream of the left breakpoint and downstream of the right breakpoint for deletions. Only windows containing at least one tested CpG were counted, with at most 100 windows per SV and 13,777,413 window records summed across all SVs. The offset accounts for differences in the amount of tested flanking sequence. A single dispersion parameter was estimated from the full count model and reused in all subset models.

To compare the contributions of the predictor groups, we partitioned explained variation using exact Shapley decomposition. We refitted the model for every subset of predictor groups, making each group’s contribution independent of the order in which groups were included. We decomposed Nagelkerke pseudo-*R*² for DMR occurrence, McFadden pseudo-*R*² for DMR count and CpG-weighted *R*² for DMR magnitude, each calculated relative to the corresponding null model.

Effects of individual annotations were estimated from the full model as adjusted log₂ odds ratios for DMR occurrence, adjusted log₂ incidence-rate ratios for DMR count and adjusted differences in DMR magnitude. Because an SV could contribute multiple SV-DMR pairs, standard errors in the magnitude models were computed using a sandwich estimator with clustering by SV. *P* values were adjusted using the Benjamini–Hochberg procedure separately within each combination of SV type, DMR direction and annotation location (insertion breakpoint, deleted sequence or DMR).

We estimated class effects using model-standardized predictions. Each class was assigned in turn to every SV while all other covariates were held at their observed values, and predictions from the full model were averaged across SVs. The resulting estimates represent the probability of DMR occurrence or the DMR count per 100 tested windows for each class, standardized to the observed covariate distribution.

Bootstrap confidence intervals were estimated from 200 replicates. We resampled merged SVs with replacement, separately for insertions and deletions, retaining all SV-DMR pairs associated with each resampled SV. This preserved dependence among pairs associated with the same SV. Class levels and the set of eligible model terms were determined from the observed data and held fixed across bootstrap replicates. Confidence intervals were defined by the 2.5th and 97.5th percentiles of the bootstrap estimates.

### Methylation in insertion-containing intervals and flanking sequence

We compared methylation between carrier and non-carrier haplotypes in the projected 200-bp insertion-containing interval and in the second through sixth 200-bp intervals on each side, comprising ten flanking intervals. One insertion-containing interval was used per insertion. The immediately adjacent intervals were excluded because individual insertion breakpoints can differ slightly from the breakpoint in the HPRC2 SV call set.

Among the 107,040 tested insertions, we excluded those on chromosomes X and Y and in the mitochondrial genome. We required at least three carrier and three non-carrier haplotypes with methylation values in the insertion-containing interval, and methylation values in at least three of the ten flanking intervals per haplotype. Both flanks were required to yield valid test results. These criteria retained 58,038 insertions.

Methylation in each flank was compared between carrier and non-carrier haplotypes using a two-sided Wilcoxon rank-sum test. Results from the two flanks were combined using directional Stouffer meta-analysis, with weights proportional to the square root of the harmonic sample size. Significant methylation differences required a Benjamini–Hochberg FDR of at most 0.05 and an absolute log₂ fold change of at least one in either the insertion-containing interval or the combined flanking analysis. For directional classification, flanking methylation differences were considered concordant when both flanks had absolute log₂ fold changes ≥ 0.5 with the same sign, opposite when both were ≥ 0.5 in absolute value with opposite signs, and intermediate otherwise.

### Inversion-associated differential methylation

Autosomal inversions were analyzed using a separate interval projection. We considered 294 candidate inversions with a minor allele frequency greater than 0.005. We aligned CHM13 to each haplotype assembly with minimap2 (v.2.29-r1283)^89^ using -x asm5 -c --secondary=no, then projected 200-bp CHM13 intervals onto each assembly with paftools.js liftover -q 5 -l 0 -d 1. After removing duplicate projections, we retained uniquely projected CHM13 intervals and excluded projections spanning at least 100 kb on the target assembly.

We obtained the inner breakpoint coordinates in the reference genome for each carrier haplotype from the HPRC2 SV call set. Inversions frequently occur within segmental duplications, so the recorded inversion span and the inner breakpoint coordinates delimit the regions containing the true breakpoints. Carrier and non-carrier status was defined using the genotypes in the HPRC2 SV call set.

A haplotype was retained when at least 80% of CHM13 intervals were projected within the inversion and within each 1-Mb flank. Within the inversion body, the proportion of intervals projected in reverse orientation was required to be at least 80% in carrier haplotypes and below 20% in non-carrier haplotypes. An inversion was retained when at least three carrier and three non-carrier haplotypes passed these criteria, retaining 141 of the 294 candidate inversions. Methylation was then tested across the merged inversion interval and 1 Mb on either side. Five inversions lying entirely within masked complex satellite sequence had no intervals available for testing, leaving 136 inversions.

Methylation was calculated using CpGs with read coverage of at least five. We tested differential methylation at the interval level using two DSS (v.2.58.0)^71^ models with smoothing. A two-group model estimated the methylation difference as mean methylation in carrier haplotypes minus mean methylation in non-carrier haplotypes. A multifactor model including inversion genotype and sequencing chemistry as covariates provided the *P* values. *P* values were adjusted globally across all tested inversion-interval combinations. DMIs were defined by an FDR below 0.05 and an absolute methylation difference greater than 10%.

We normalized positions within inversion bodies to account for differences in inversion length when summarizing DMIs. For each inversion, we used either the boundaries in the HPRC2 SV call set or the median inner breakpoint coordinates across carrier haplotypes, selecting the estimate closest to the boundaries supported by assembly alignments. The median inner interval was used for 84 inversions and the boundaries from the HPRC2 SV call set for 52. We then generated overlapping 10-kb windows with a 5-kb step size across each inversion body and its 1-Mb flanks. Window midpoints within the inversion body were assigned to 200 normalized bins, each representing 0.5% of the inversion length, whereas flanking windows retained physical coordinates. At each position, the proportion of inversions with a DMI of the specified direction was calculated among inversions tested at that position. Each inversion was counted once per normalized bin, giving equal weight to each inversion. Ten inversions with tested intervals in fewer than 20% of the 200 bins within the inversion body were excluded, leaving 126.

### CGI calling

CGIs were called separately in each assembly with the UCSC cpg_lh algorithm. The algorithm identifies maximally scoring segments using a weight of +17 for CG dinucleotides and −1 at all other positions, and retains candidates with GC content of at least 50%, length greater than 200 bp and an observed-to-expected CpG ratio above 0.6^2^.

We called CGIs separately on unmasked and repeat-masked sequence from each assembly. The unmasked call set was generated from the full assembly sequence. For the masked call set, we hard-masked the union of RepeatMasker annotations^78^ for that assembly and merged tandem repeats^90^ with a repeat unit length of at most 12 bp before calling CGIs.

We benchmarked this procedure on CHM13 by comparing both CGI call sets with the UCSC CGI annotation, measuring concordance as the proportion of calls with identical coordinates, with reciprocal overlap of at least 90%, and with any overlap.

### Construction of the PanCGI catalogue

PanCGI integrates CGI calls made independently in each assembly into a sequence-resolved catalogue. We used path-coordinate maps generated with GrAnnoT^91^ to represent each CGI call as an ordered walk of oriented segments in the pangenome graph. Each walk was represented by overlapping subpaths of three consecutive segments. We compared CGI calls using weighted Jaccard similarity across these subpaths. Each subpath was weighted by the inverse of its frequency across the catalogue, giving greater weight to less frequent subpaths. CGI calls with similar graph walks were grouped together.

We assigned each group a corresponding position in CHM13, using the insertion site for CGI calls located within an insertion. Groups overlapping an annotated CHM13 CGI defined a reference locus. The remaining groups were combined into novel loci when neighbouring groups mapped within 100 bp of one another in CHM13. For each locus, the start and end coordinates in CHM13 were defined as the median start and median end coordinates, respectively, of all CGI calls assigned to that locus. Each CGI call assigned to a locus was termed a member and retained its own assembly sequence.

Within each locus, the member sequences were compared pairwise by semi-global alignment after orienting all sequences consistently. Members were clustered by sequence similarity and length, requiring at least 80% sequence identity and a length ratio of at least 0.8 for every pair within a cluster. Each candidate member had to meet both criteria with every member already in the cluster. The member with the smallest average sequence distance to the other members was selected as the representative of each cluster. Its sequence defined a CGI allele.

Each assembly was then genotyped at every locus as CGI-positive if it contributed a member. For assemblies without a CGI member, we assessed whether the homologous locus could be recovered by projecting the graph segments of the locus onto the corresponding assembly path. The homologous locus was considered recoverable if its flanking segments were identified on the same chromosome, in the same orientation, in the expected left-to-right order and within a plausible genomic span, or if the segments of the locus itself were recovered as a coherent chain. Assemblies with a recoverable homologous locus but no CGI member were genotyped CGI-negative. Genotypes were recorded as missing when the homologous locus could not be reliably identified, because CGI absence could not be distinguished from absence of the homologous sequence.

The catalogue was generated using Python (v.3.12.13), with pairwise sequence alignments performed using parasail (v.1.3.4)^92^.

### SV origins of novel CGI alleles

For each novel CGI allele associated with an SV from the HPRC2 SV call set, we classified its sequence origin according to the positional relationship between the CGI and the matched SV on the carrier assembly. An allele was classified as INS-internal when the CGI was fully contained within an insertion, INS-junction when the CGI and insertion overlapped but neither contained the other, and INS-spanning when the CGI contained the insertion. For deletions, an allele was classified as DEL-junction when the deletion junction fell within the CGI, with CGI sequence on both sides. Alleles meeting both an insertion criterion and the deletion-junction criterion were assigned to INS+DEL. These five sequence-origin classes were mutually exclusive; where an allele overlapped multiple insertions, INS-internal took precedence over INS-junction and INS-spanning. Alleles not assigned to one of these SV-related classes were evaluated by the reconstruction analysis described below.

### Reconstruction of novel CGI alleles

We examined the sequence changes underlying novel CGI alleles. Candidate members were selected from novel CGI alleles with CHM13 coordinates and no assigned SV-related origin, and we tested whether small variants alone could account for each member’s CGI sequence. For each member, we extended the homologous CHM13 interval by 2 kb on each side. The flanking sequence was included to capture variants near the boundaries of the homologous interval that could contribute to CGI formation.

Phased small variants from the HPRC2 PanGenie variant set were applied to the CHM13 sequence separately for each haplotype, after checking each allele against the reference. Sequences projected to CHM13 in reverse orientation were reverse-complemented before comparison.

We aligned each assembly CGI sequence to both the extended CHM13 reference sequence and the reconstructed haplotype sequence using edlib (v.1.3.9)^93^ in infix mode. Sequence identity was recorded for each alignment.

To identify small variants contributing to sequence reconstruction, we omitted each variant in turn while retaining all others and repeated the alignment. A variant was counted if its inclusion reduced the edit distance or increased sequence identity relative to the reconstruction without that variant. We recorded the number of these variants for each CGI member. For sequence-origin classification, a member provided small-variant support when reconstruction using small variants alone achieved a sequence identity of at least 0.99. A novel CGI allele was classified as SmallVar-induced when at least one member met this criterion; alleles for which no member met the criterion were classified as unresolved. The unresolved category indicates that the sequence origin could not be assigned. This analysis classified 2,459 CGI alleles at 2,279 novel loci as SmallVar-induced and 584 CGI alleles as unresolved.

### CGI population genetics

We assessed population structure in CGI PAV separately at locus and allele resolution in 231 HPRC2 individuals (462 haplotypes, including HG002). At each locus, haplotype genotypes were encoded as 1 for CGI-positive and 0 for CGI-negative. At allele resolution, presence or absence was encoded separately for each CGI allele. Haplotype genotypes that could not be determined were recorded as missing. The two haplotype values for each individual were summed when both genotypes were available, yielding diploid dosages of 0, 1 or 2; the diploid genotype was recorded as missing when either haplotype genotype was missing. At multiallelic loci, each CGI allele was retained as a separate feature.

We retained autosomal polymorphic CGI loci and alleles with a cohort-wide haplotype call rate of at least 0.95, yielding 12,118 loci and 28,687 alleles. We performed principal-component analysis of the dosage matrices using PLINK2 (v.2.0.0-a.6.9LM)^94^ with --maf 0.01 --geno 0.05 --pca 20 allele-wts. After PLINK2 filtering, 5,559 CGI loci and 15,501 CGI alleles remained. **Extended Data Fig. 7c,d** show principal components 1 and 2 (PC1 and PC2) from the locus- and allele-level analyses, respectively.

We partitioned CGI PAV using a nested analysis of variance of individual CGI dosages into components among superpopulations, among populations within superpopulations, and among individuals within populations. A CGI locus or allele was included if dosage values were available for at least three individuals and varied among those individuals. Missing dosages were excluded separately for each locus or allele. Variance components were estimated for five CGI sets: polymorphic loci, novel loci, novel alleles, SV-related novel alleles and SmallVar-induced novel alleles.

We assessed CGI allele-frequency differentiation for individual alleles and for alleles grouped by sequence origin. For per-allele analyses, we retained autosomal polymorphic CGI alleles with a cohort-wide call rate of at least 0.90 and a minor allele frequency of at least 0.01, and calculated Hudson’s *F* ^36^ for each superpopulation against the remainder of the cohort. For each sequence-origin class, pairwise differentiation between superpopulations was calculated as the sum of Hudson numerator terms across alleles divided by the sum of the corresponding denominator terms.

### Local genealogy inference

We inferred local genealogies with Relate, which requires ancestral and derived alleles to be distinguished at each variant site. Human ancestral-base estimates were obtained from the Ensembl multi-primate Enredo–Pecan–Ortheus (EPO) alignment^95^ in GRCh38 coordinates. Because our variants and CGI loci were defined in CHM13 coordinates, we mapped the ancestral bases from GRCh38 to CHM13 before genealogy inference.

We used the HPRC2 PanGenie variant set for 231 HPRC2 individuals (462 haplotypes). Using PLINK2 (v.2.0.0-a.6.9LM)^94^, we retained autosomal biallelic SNVs with --snps-only just-acgt and required non-missing genotypes in all individuals with --geno 0, yielding 31,865,692 variants.

At CHM13 positions with multiple retained variant records, we excluded all records because Relate requires a single variant per position. This removed 394,982 records at 196,612 coordinates. We then mapped each remaining position from CHM13 to GRCh38 and back to CHM13. We required a unique CHM13-to-GRCh38 mapping and a return mapping that recovered the original single-base CHM13 interval on the positive strand, excluding 262,192 positions that failed these criteria. The EPO ancestral base was retrieved at the corresponding GRCh38 position and complemented when the CHM13-to-GRCh38 mapping was on the negative strand. Because ancestral bases were reported on the GRCh38 forward strand, complementation ensured consistent orientation with the CHM13 reference and alternate alleles. We excluded 1,648,358 positions without an ancestral base and 163,205 at which the ancestral base matched neither the reference nor the alternate allele. The remaining 29,396,955 ancestral bases were used to construct a CHM13 ancestral sequence, with all other positions encoded as N.

During Relate input preparation, we excluded a further 18,327 sites that were monomorphic in the cohort and 9,634 sites in regions masked as described above, leaving 29,368,994 variants. Haplotype genotypes were recoded at the 2,918,751 sites where the ancestral base matched the alternate allele, so that 0 represented the ancestral allele and 1 the derived allele. This allele coding was retained throughout subsequent analyses. We ran Relate (v.1.2.4)^37^ separately for chromosomes 1–22 with --mode All, using recombination maps constructed directly in CHM13 coordinates. We set a mutation rate of 1.25 × 10^−8^ per base per generation and an initial effective population size of 30,000. Effective population size was then re-estimated jointly across the autosomes over five iterations, assuming 28 years per generation. Each iteration sampled branch lengths under the current coalescence rates and re-estimated those rates from the sampled trees. Final branch lengths were re-estimated on the original genealogies under the converged rates. The genealogies were then converted to tree sequences, each sample node corresponding to one of the 462 haplotypes.

### Assigning CGI loci to local genealogies

To infer whether a novel CGI had been gained or lost along a lineage, we assigned each locus to the local genealogy of the haplotypes carrying the surrounding sequence. A local genealogy relates the 462 haplotypes through their inferred common ancestry at a genomic position: its leaves represent the sampled haplotypes, its internal nodes their inferred common ancestors, and its root the inferred common ancestor of all 462 haplotypes. Genealogies and CGI loci were both defined in CHM13 coordinates.

We analyzed the 2,279 SmallVar-induced novel loci. Each locus was assigned to the local genealogy at its CHM13 position, with one tree used to represent each locus. A haplotype was classified as CGI-positive when a CGI was called at the locus in its assembly, CGI-negative when the locus could be genotyped but no CGI was called, and missing when the locus could not be genotyped. Across the 2,279 loci, this yielded 141,343 CGI-positive, 896,289 CGI-negative and 15,266 missing haplotype-locus states. Missing haplotype-locus states contributed no evidence to the subsequent ancestral-state and trajectory analyses.

Local genealogies were available for 2,166 loci. Of the remaining 113 loci, 57 were on sex chromosomes, for which genealogies were not inferred, and 56 fell outside the genomic range covered by the inferred local genealogies. All 113 loci were recorded as lacking a usable genealogy.

### Ancestral state reconstruction

We reconstructed the history of CGI gain and loss from present-day haplotype states using weighted parsimony. This assigns CGI states to ancestral nodes to minimize the total cost of state changes across the genealogy. CGI loss and gain were assigned costs of one and three, respectively, imposing a higher penalty on repeated independent gains than on repeated losses. Missing haplotype states were assigned equal costs for both CGI states and provided no evidence for either state. A locus was excluded if its genealogy lacked a single root.

For each locus, we calculated the minimum parsimony cost twice, with the root fixed as CGI-negative or CGI-positive. The root state with the lower cost was assigned as the inferred ancestral CGI state; loci with equal costs were classified as equivocal. A cost difference of at least two indicated a supported root-state assignment, and a smaller difference indicated weak support. Both the ancestral state and trajectory class were determined by the lower-cost reconstruction regardless of this support category, which was used only for the confidence criteria described below.

Because multiple reconstructions can attain the same minimum parsimony cost, we enumerated the ancestral states and transition branches across all optimal reconstructions. Across these reconstructions, we determined the minimum and maximum numbers of CGI gains and losses. A transition on a given branch was required if it occurred in all optimal reconstructions and possible if it occurred in only some. A locus was classified as recurrent only if no optimal reconstruction included a transition in the opposite direction.

The gain cost of three was specified for parsimony analysis and did not represent an estimate of the relative rates of CGI gain and loss. We repeated the reconstruction with gain costs of one, two and five to assess sensitivity to this choice. Parsimony was used to analyse discrete CGI states defined by assembly calls on genealogies of human haplotypes. We lacked the information needed to specify a likelihood model: a calibrated transition matrix for CGI gain and loss, a model of differential detection of gains and losses, and an outgroup CGI state. The inferred ancestral CGI state therefore refers to the root of the sampled human genealogy.

### Trajectory classification

Under a single-lineage CGI gain or loss, haplotypes whose CGI state differs from the inferred ancestral state are expected to belong predominantly to one clade in the local genealogy. We used 300,000 autosomal variants outside the candidate CGI loci to calibrate a criterion for consistency with a single clade. Within six bins of derived-allele count, we compared the carrier haplotypes for each variant with the descendants of the branch on which the variant was inferred to have arisen. The resulting distributions of Jaccard similarity, precision and recall defined the empirical range of values expected for haplotypes descending from a single branch.

For each locus, we compared haplotypes whose CGI state differed from the inferred ancestral state with the descendant haplotypes of every branch in the local genealogy. The branch with the highest Jaccard similarity was selected; ties were resolved sequentially by fewer discordant haplotypes, more haplotypes shared between the two sets, higher precision and higher recall. A locus was considered consistent with a single clade when all three measures for the selected branch met or exceeded their fifth-percentile calibration thresholds in the corresponding bin. The tenth- and first-percentile thresholds were used as stricter and more permissive criteria, respectively, to assess classification robustness and identify loci near class boundaries.

Loci were classified in a fixed order, with each locus assigned to the first category whose criteria it met. A locus was recorded as lacking a usable genealogy if either the genealogy or its CHM13 coordinate was unavailable. Loci with equal minimum parsimony costs for CGI-positive and CGI-negative roots were classified as equivocal. A locus whose optimal reconstruction required no CGI state change was recorded as having no transition within the human genealogy. Among the remaining loci, those meeting the single-clade criterion were classified as single-lineage loss under a CGI-positive root or single-lineage gain under a CGI-negative root. A locus that did not meet the single-clade criterion was classified as recurrent gain or loss if at least two transitions in the direction implied by the root state were required and no optimal reconstruction included a transition in the opposite direction. All remaining loci were classified as turnover.

Of the 2,279 candidate loci, 857 showed single-lineage gain, 222 single-lineage loss, 258 recurrent gain, 172 recurrent loss and 524 turnover. Among the turnover loci, 166 had a CGI-negative inferred ancestral state and 358 had a CGI-positive inferred ancestral state. A further 132 loci had equivocal ancestral states, one required no CGI state change within the human genealogy and 113 lacked a usable genealogy.

### Long-read transcript catalogue

We constructed transcript models from 202 Iso-Seq datasets and subsequently lifted the transcript annotation to CHM13. Full-length non-concatemer (FLNC) reads from each sample were assembled into transcript models using IsoQuant (v.3.6.3)^96^ with GRCh38 as the reference genome and GENCODE v49 as the reference gene annotation. We assessed the transcript models from each sample using SQANTI3 (v.5.4)^97^, which compares transcript structures with the reference annotation and evaluates support for transcript ends and splice junctions. Supporting evidence comprised cap analysis of gene expression (CAGE) peaks from refTSS^98^, polyadenylation motifs, polyadenylation site clusters^99^ and splice-junction coverage from Intropolis^100^.

We filtered transcript models using the following criteria. A full-splice-match model was retained if the adenine content of the genomic sequence downstream of its 3′ end was below 60%, thereby excluding models with evidence of internal priming. Models in all other SQANTI3 structural classes were required to meet this criterion, show no evidence of reverse-transcriptase template switching, and have either only canonical splice junctions or junction coverage of at least ten in Intropolis. Models excluded by these criteria were re-evaluated using the SQANTI3 rescue step, with rescue restricted to full-splice-match models.

We merged transcript models across samples and with GENCODE v49^39^ using TAMA merge^101^. Models with the same splice structure were merged when their 5′ and 3′ ends were within 50 bp; internal splice-junction coordinates were required to match exactly. GENCODE v49 was designated as the source of gene names and coding sequences. The merged annotation was assessed using the SQANTI3 quality-control module, without additional filtering or rescue. Transcript models on chrEBV were removed.

Genes detected only in the Iso-Seq data and absent from GENCODE v49 were required to meet at least one of two support criteria. A gene was retained if at least one transcript had abundance above 0.1 transcripts per million and a full-length read count of at least five in at least ten of the 201 quantified samples. Alternatively, a gene was retained if the TAMA merge report showed support from at least ten distinct Iso-Seq samples. Genes present in GENCODE v49 were retained without these additional criteria. Annotation attributes were preserved for all retained records.

Transcript classification and coding potential were assigned from the SQANTI3 annotation of the final filtered GRCh38 transcript catalogue. Full-splice-match isoforms were classified as annotated, whereas all other SQANTI3 structural classes were classified as novel. Novel isoforms with a coding prediction were further classified as protein coding or sensitive to NMD according to the SQANTI3 NMD prediction; isoforms without a coding prediction were classified as non-coding.

Alternative-splicing events in the HPRC2, GENCODE and combined transcript catalogues were identified using SUPPA (v.2.4)^102^ with strict transcript-boundary matching. Event counts were summarized by SUPPA event class for each catalogue.

We lifted the annotation from GRCh38 to CHM13 using Liftoff (v.1.6.2)^103^ with minimap2 for alignment, allowing additional gene copies at a sequence-identity threshold of 0.95. We re-predicted coding sequences using CHM13 sequence to account for potential reading-frame differences between assemblies. The lifted gene, transcript and exon coordinates were held fixed, and SQANTI3 (v.5.5.4)^97^ with TransDecoder2^104^ was run separately for each chromosome to predict coding sequences against CHM13. We mapped the predicted coding segments to the lifted exons and ordered them from 5′ to 3′ according to transcript strand. Each coding segment was assigned the phase recorded in GFF3, defined as the number of bases from its 5′ end to the start of the next complete codon. When the combined length of preceding coding segments was divisible by three, the phase was zero. Otherwise, the phase was three minus the remainder after division of that length by three.

A predicted coding sequence was retained only if every coding segment lay within a lifted exon and no two segments overlapped. The chromosome and strand of each segment were required to match those of the transcript. The total coding-sequence length had to be divisible by three, and translation had to yield no internal stop codons. Transcripts failing any of these criteria retained their exon structures and were left without coding-sequence annotations.

### Variant calling and haplotype phasing

We called small variants from high-fidelity (HiFi) reads aligned to CHM13 using DeepVariant (v.1.6.1)^105^ with run_deepvariant --model_type PACBIO. For male samples, we additionally specified --haploid_contigs “chrX,chrY” and provided a BED file defining the pseudoautosomal regions. An assembly-based variant call set was generated for each individual using dipcall^106^ on the corresponding pair of haplotype assemblies, with run-dipcall -W and a repetitive 19-mer list. For male samples, dipcall was run with -x and the same pseudoautosomal-region BED file. Variants passing the respective caller’s filters were normalized against CHM13 using BCFtools (v.1.21)^107^ norm -f CHM13 --check-ref ws -m -any -d both.

We combined assembly-based and read-based phasing information. dipcall provided chromosome-wide phase information from the haplotype assemblies. WhatsHap (v.2.8)^108^ used read evidence to phase variants into internally consistent phase blocks, whose haplotype orientations were independent across blocks.

We phased the DeepVariant calls with WhatsHap using both the HiFi read alignments to CHM13 and the phased dipcall variants as evidence. We then oriented each resulting phase block to match the dipcall haplotype assignments and merged the oriented DeepVariant calls with the dipcall calls into a single phased variant set per individual.

### Iso-Seq read phasing

For variant-based read phasing, FLNC reads from each individual were aligned to CHM13 using pbmm2 (v.1.17.0)^89^ with --preset ISOSEQ. We also aligned the reads to each of the individual’s two haplotype assemblies separately and to a reference containing both haplotype assemblies, using the same preset.

We assigned Iso-Seq reads to haplotypes using variant-based and assembly-based approaches. WhatsHap haplotag used the phased variant set to assign CHM13-aligned reads to haplotypes, recording the haplotype and phase block for each assigned read. Independently, we compared each read’s alignments to the two haplotype assemblies. Reads were assigned to the haplotype with the higher gap-compressed sequence identity; ties were resolved by the lower edit distance. When both measures were equal, the assignment was based on the read’s alignment to the reference containing both haplotype assemblies.

For each read, we used the WhatsHap haplotype assignment when available and the assembly-based assignment otherwise. CHM13 alignments of assigned reads were extracted by read name to generate one alignment set per haplotype.

### Isoform, splicing and methylation phenotypes

We quantified isoform abundance against the combined transcript catalogue lifted to CHM13 using Bambu (v.3.12.0)^109^ with bambu(annotations, genome, discovery = FALSE, quant = TRUE). Abundance was estimated for the supplied transcript models without additional transcript discovery. For each individual, quantification was performed separately using all FLNC alignments to CHM13 and the alignments assigned to each haplotype, yielding total and haplotype-specific abundance estimates for each transcript.

Transcripts were included in QTL analysis if their abundance exceeded 0.1 transcripts per million in at least 20% of samples and their full-length read count was at least five in at least 20% of samples.

We extracted splice junctions from the CHM13 FLNC alignments with RegTools (v.1.0.0)^110^ using junctions extract -a 8 -m 50 -M 2500000 -s XS. We clustered the junctions across all 606 alignment sets with LeafCutter (v.0.2.9) using leafcutter_cluster_regtools.py^111^ -l 2500000. Each junction was assigned to an intron cluster, and junction read counts were reported. Within-cluster junction usage fractions were calculated with prepare_phenotype_table.py.

Introns were filtered by read support and annotation context. An intron was retained if its read count exceeded 0.1 counts per million in at least 20% of samples and it had at least five supporting reads in at least 20% of samples. Clusters with fewer than two retained introns were excluded. Retained introns were further required to overlap the combined transcript annotation on CHM13 and to lie on an autosome. The splicing phenotype for each retained intron was its within-cluster usage. Methylation phenotypes were derived from the haplotype-specific intervals projected from the 200-bp CHM13 intervals described above. For each individual and interval, we summed methylated read counts and valid read coverage across CpGs with valid read coverage of at least five on the corresponding haplotype. Sample-level methylation was calculated as the total methylated read count across both haplotypes divided by the total valid read coverage. Allele-specific methylation phenotypes were calculated separately for the two haplotypes.

### Covariates

Covariate matrices for QTL mapping were constructed separately for methylation, isoform abundance and splicing phenotypes. Thirty PEER factors were estimated separately for each phenotype class from its transformed phenotype matrix using peertool -n 30 -i 2000. All three covariate matrices included sex and the first ten genetic principal components. Methylation covariates additionally included sequencing chemistry, the z-score of log₂ mean nanopore coverage weighted by the number of CpG sites, and the z-score of read N50. Isoform and splicing covariates additionally included Kinnex sequencing pool indicators for the 32 libraries. Splicing covariates also included log_10_-transformed junction library size.

Genetic principal components were computed from variants pruned for linkage disequilibrium within each superpopulation. Autosomal biallelic SNVs from the HPRC2 PanGenie variant set were imported into PLINK2 using --vcf-half-call missing --autosome --snps-only just-acgt. Within each of the five superpopulations, we applied variant quality control with --maf 0.01 --geno 0.10 --hwe 1e-6 midp and performed linkage disequilibrium pruning with --indep-pairwise 50 10 0.2. The --bad-ld option was used for superpopulations with fewer than 50 samples. To reduce the influence of unequal superpopulation sample sizes on variant selection, we retained variants selected in at least four of the five pruned sets. We then applied quality control in the combined cohort with --maf 0.01 --geno 0.10 and computed principal components from the resulting variant set. The first ten components were included as covariates.

### Variant panels for molecular QTL mapping

Small variants were obtained from the HPRC2 PanGenie variant set^61,86^ and were retained at minor allele frequency greater than 0.01. SVs were obtained from the HPRC2 SV call set^29^ and retained at minor allele frequency greater than 0.005. Variants were required to have genotype missingness of at most 0.20. Missingness was recalculated after subsetting to each molecular cohort, and the same threshold was reapplied.

The mQTL panel comprised 19,306,360 autosomal variants. isoQTL and sQTL analyses used 19,304,602 variants after missingness filtering in the RNA cohort.

### Allele-aware QTL mapping

Molecular QTLs were mapped with mixQTL (v.0.2)^43^, which estimates a shared variant effect using two approximately independent regressions: one models allelic imbalance between an individual’s two haplotypes, and the other models the total phenotype across both haplotypes. The effect estimates are combined by inverse-variance weighted meta-analysis. Isoform abundance was analyzed using the published count-based formulation. We developed both regressions for methylation and splice-junction usage, which are measured as proportions.

Because the log ratio is undefined for zero proportions and its variance depends on the proportion, we replaced the logarithm with the arcsine transformation described above and derived the variance from read coverage. For individual *i*, let 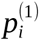 and 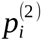 be the two haplotype proportions, *p_i_* the proportion across both haplotypes, 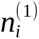 and 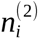 the corresponding haplotype read coverages, and 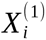 and 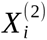 the two haplotype genotypes. The allele-specific phenotype (ASP) and total phenotype (TP) regressions were defined as follows:

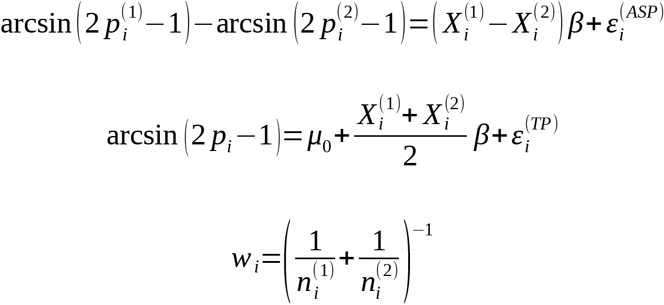

Here, β is the shared per-allele effect, *μ*_0_ is the intercept, and 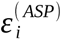 and 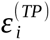 are the residuals of the ASP and TP regressions, respectively.

The ASP regression was fitted by weighted least squares using 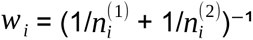, equal to half the harmonic mean of the two haplotype coverages. These inverse-variance weights replace the count-based weights. To limit the influence of individuals with very high coverage, weights were capped at the minimum weight for the feature multiplied by one tenth of the sample size. For methylation, *p_i_* was the total methylated read count across both haplotypes and all CpGs in the interval divided by the total valid read coverage. For splicing, the corresponding proportions were calculated from splice-junction read counts.

Variants were tested within 1 Mb of a methylation interval or an isoform transcription start site, and within 500 kb of a splicing-cluster midpoint. Association significance was assessed using 1,000 permutations of individuals for each feature in mixQTL, with the meta-analysis statistic recalculated for each permutation. Because 1,000 permutations provide an empirical *P*-value resolution of approximately 0.001, we fitted a beta distribution by maximum likelihood to the minimum nominal *P* value from each permutation, following FastQTL^112^. The fitted distribution was used to estimate the permutation *P* value for the observed lead variant. Storey^113^ *q* values were calculated across features within each phenotype class, and features with *q* < 0.05 were considered significant. For each significant feature, the association lead variant was defined as the tested cis variant with the smallest nominal association *P* value.

### Fine-mapping

We fine-mapped each significant molecular feature with mixFine, the SuSiE-based counterpart of mixQTL^43^, to identify candidate causal variants. mixFine jointly fits the allelic-imbalance and total-phenotype data using a single sum-of-single-effects model. For a feature measured in *N* individuals and tested against *P* variants, the two regression design matrices were stacked to form a 2*N* × *P* matrix. In the model below, *y^(ASP)^* is the allelic-imbalance phenotype 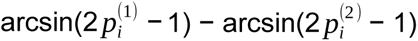 and *y^(TP)^* is the total phenotype arcsin(2 *p_i_* − 1):

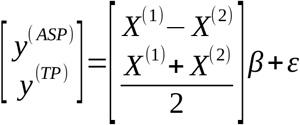

Here, *y^(ASP)^* and *y^(TP)^* are the *N*-element ASP and TP phenotype vectors; *X*^(1)^ and *X*^(2)^ are the *N* × *P* phased haplotype-genotype matrices; β is the *P*-element vector of shared variant effects; and ε is the 2*N*-element residual vector.

ASP observations were weighted by *w_i_*, and TP observations were assigned unit weights. Each regression component was then scaled by its estimated residual standard deviation. The TP phenotype vector and genotype matrix were mean-centred, and SuSiE (v.0.14.2)^44^ was fitted without further standardization or an intercept.

The ASP and TP phenotypes were defined as described above. Methylation and splice-junction usage were analyzed using arcsine-transformed proportions. Isoform abundance was analyzed using the published count-based formulation, with covariate effects regressed out before model fitting.

We allowed up to ten single effects per feature and constructed 95% credible sets. Credible sets were retained if the minimum absolute pairwise correlation among their variants was at least 0.5. The cis windows matched those used for QTL mapping: 1 Mb around a methylation interval or an isoform transcription start site and 500 kb around a splicing-cluster midpoint. For each feature, we reported the credible sets, the number of variants in each set and the posterior inclusion probability (PIP) of each variant.

### Colocalization among molecular QTL classes

We performed pairwise colocalization with coloc (v.6.0.1)^47^ to distinguish evidence for a shared causal variant from evidence for distinct nearby causal variants.

For each phenotype, we obtained the log Bayes factor matrix from mixFine, with one row per single effect and one column per variant across the full cis window. The two matrices were supplied to coloc.bf_bf^114^ with prior probabilities p1 = 1 × 10^−4^, p2 = 1 × 10^−4^ and p12 = 1 × 10^−5^. A pair of single effects was evaluated only when variants shared between the two datasets captured at least 50% of the non-null posterior support for each effect. We analyzed all three pairwise combinations of methylation, isoform abundance and splice-junction usage.

For each pair of single effects, coloc estimated the posterior probabilities of the colocalization hypotheses. PP3 and PP4 denote the posterior probabilities that the two phenotypes have distinct causal variants or share a causal variant, respectively. A pair was considered colocalized if PP3 and PP4 summed to at least 0.9, supporting an association for both phenotypes at the locus, and PP4 was at least three times PP3. For each colocalized pair, variants were ranked by their posterior probability of being the shared causal variant under H4. The 95% credible set comprised the highest-ranked variants whose cumulative posterior probability first reached or exceeded 0.95.

### Harmonization of QTL and GWAS statistics and joint colocalization

We tested whether molecular QTL and GWAS associations shared a causal variant. GWAS summary statistics for 70 non-redundant diseases and quantitative traits were obtained from FinnGen release 12^50^, the NHGRI-EBI GWAS Catalog^48^ and Pan-UK Biobank^49^. Autosomal summary statistics were harmonized to a common format. Odds ratios were converted to log odds ratios, missing standard errors were calculated from the effect size and *P* value, and records with incomplete coordinates, alleles or association statistics were excluded.

GWAS summary statistics reported in GRCh37 coordinates were lifted to GRCh38 using Picard LiftoverVcf (v.3.4.0)^115^ with RECOVER_SWAPPED_REF_ALT=true, after assigning alleles using the GRCh37 reference and normalizing variants with bcftools. When the source effect allele corresponded to the GRCh38 reference allele after liftover, the sign of the effect size was reversed and the effect-allele frequency was replaced by one minus its original value.

For each GWAS trait, variants with *P* < 5 × 10^−8^ were ranked by ascending *P* value. We iteratively selected the remaining variant with the smallest *P* value as a lead variant and excluded all other genome-wide significant variants within 500 kb on the same chromosome from subsequent lead-variant selection. Each lead variant defined a 1-Mb window extending 500 kb in either direction.

QTL variants were lifted from CHM13 to GRCh38 using LiftoverVcf with RECOVER_SWAPPED_REF_ALT=true and the allele-handling procedure described above. QTL and GWAS records were then matched by chromosome and position in GRCh38 coordinates. When the effect and non-effect alleles matched between the two records, the QTL effect size was retained. When those alleles were interchanged, the sign of the QTL effect size was reversed. All other allele combinations were excluded. All retained QTL effect sizes were aligned to the GWAS effect allele, with GWAS statistics unchanged.

HyPrColoc requires the same variants across all traits in a window, so molecular features with limited variant coverage can reduce the number of shared variants. Within each window, we used beam search to identify feature subsets whose removal maximized the number of shared variants. No more than 15% of features were removed, and removal required an increase of at least 500 variants and at least 10% relative to the original shared-variant count. Among subsets that achieved at least 80% of the maximum increase, we selected the smallest subset whose removal left at least 1,000 shared variants. If all features of a QTL class were removed, the class was restored when it alone retained at least 1,000 shared variants. Pruning removed 1.22% of feature-window pairs.

For each window, we constructed effect-size and standard-error matrices containing the GWAS trait and all retained methylation, isoform and splicing features. The matrices required complete effect-size estimates and positive standard errors for every variant-trait combination. We analyzed the matrices with HyPrColoc (v.1.0)^52^ using hyprcoloc(prior.1 = 1e-4, prior.c = 0.02, uniform.priors = FALSE, reg.thresh = “default”, align.thresh = “default”, snpscores = TRUE). For each cluster of colocalized traits, we reported the PPFC, the regional association probability and the proportion of the colocalization posterior probability explained by the candidate variant.

### Annotation enrichment of fine-mapped variants

We assessed enrichment across three annotation categories: genomic intervals, predicted variant consequences and chromatin states. The genomic interval annotations were those defined above, supplemented with the novel CGI loci identified in this study. We predicted variant consequences with the Ensembl Variant Effect Predictor (release 116)^116^, using the CHM13 assembly and the combined transcript catalogue described above so that consequences refer to the same transcript models. We used the GM12878 15-state ChromHMM annotation (Roadmap E116)^46^ for chromatin states. Annotations for this lymphoblastoid cell line were lifted from GRCh38 to CHM13, and intervals assigned to the same state were merged.

To account for QTL ascertainment in annotation-enrichment analyses, query variants were compared with matched background variants. Query variants were defined as the variant with the highest PIP in each independent credible set, with duplicate variants removed within each QTL class and no PIP threshold applied. The background comprised all unique variants tested in the cis windows of significant features included in fine-mapping. These cis windows were the same as those used for QTL mapping. Each variant was represented by its genomic interval in CHM13 coordinates.

Each query variant was matched to background control variants by QTL class, minor allele frequency, distance to the nearest feature position used for QTL mapping, variant class and variant length.

For each query variant and annotation, we calculated the proportion of eligible matched controls with that annotation. Using this proportion as the success probability, we sampled one Bernoulli outcome per query variant and repeated the procedure 1,000 times. Enrichment was calculated as the proportion of annotated query variants divided by the mean annotated proportion across control replicates and was reported on a log₂ scale. A pseudocount of 0.5 was applied when calculating each proportion to avoid undefined ratios when the annotation was absent from either the query or control variants. The interval for the enrichment estimate was defined by the 2.5th and 97.5th percentiles of the replicate values. Statistical significance was assessed using a one-sided empirical *P* value for enrichment, with Benjamini–Hochberg correction within each QTL class and annotation category.

### Population-differentiated mQTL support for population DMRs

We used mQTLs to relate the population methylation differences described above to individual variants. We assessed how often a population DMR was supported by an mQTL whose effect allele differed in frequency between populations and whose predicted methylation shift agreed in direction with the observed regional difference.

For this population analysis, effect-allele frequencies and mQTL effect sizes were defined relative to the CHM13 alternate allele. For each mQTL credible set, we selected the variant with the highest PIP and required PIP ≥ 0.5. For each of the five superpopulations, we compared the effect-allele frequency in the focal superpopulation with that in all remaining haplotypes. A variant was considered population-differentiated if the absolute frequency difference was at least 0.1 and the effect allele was at least twofold enriched or depleted in the focal superpopulation. For enriched alleles, the frequency ratio was the frequency in the focal superpopulation divided by that in all remaining haplotypes; for depleted alleles, the reciprocal was used.

Support for a population DMR was evaluated using its constituent 200-bp methylation intervals. An interval was assigned to a DMR if its midpoint fell within the DMR. The predicted methylation shift was calculated as the mQTL effect size multiplied by the effect-allele frequency difference. An interval supported the DMR when the predicted shift agreed in sign with the observed methylation difference for that interval between the focal superpopulation and all remaining haplotypes. For example, an AFR-hypermethylated DMR could be supported by an allele enriched in AFR that increased methylation or by an allele depleted in AFR that decreased methylation. Allele enrichment and depletion were defined relative to the focal superpopulation. Both types of allele could support the same DMR because support depended on the product of the effect size and the allele-frequency difference. A DMR was considered supported when at least 30% of its intervals agreed in sign and fewer than 20% disagreed. Support was assessed separately for enriched and depleted effect alleles, so a DMR could be supported by either or both. Each unique combination of superpopulation, variant and DMR direction was counted once, regardless of the number of supporting intervals.

We compared the population-differentiated mQTL support with the joint colocalization results. A match required the same variant-methylation interval pair in a completed HyPrColoc analysis that included a GWAS trait. We summarized the overlap at two levels: the supported combinations defined above and GWAS lead windows in which an mQTL was included in the joint analysis. Each window was counted once.

## Data availability

HPRC2 data, including assemblies and long-read sequencing data, are available at https://github.com/human-pangenomics/hprc_intermediate_assembly. The HPRC2 SV call set analyzed in this study is available through the 1KGP long-read structural-variation resource at https://github.com/jiadong324/1KG_LongRead_SV. GENCODE v49 annotations are available at https://www.gencodegenes.org/human/release_49.html. T2T-CHM13v2.0 and its reference annotations are available at https://github.com/marbl/CHM13. The CHM13-native scaled recombination maps used for autosomal local genealogy inference are available at https://github.com/JosephLalli/phasing_T2T/tree/main/resources/recombination_maps/t2t_native_scaled_maps. ENCODE candidate cis-regulatory element annotations are available from the ENCODE Portal under accessions ENCFF036NSJ, ENCFF196MIP, ENCFF262LCI, ENCFF379UDA and ENCFF535MKS. Processed DNA methylome and molecular-QTL resources generated in this study, including genome-wide methylation- and CpG-density-variability tracks, HVMR and HVDR calls, population DMRs, the combined transcript annotation, molecular-QTL lead associations and fine-mapping results, and molecular-QTL and GWAS colocalization results, are available at https://github.com/tianjie16/HPRC2_methylation_framework. Sequence-resolved PanCGI resources are available at https://github.com/tianjie16/PanCGI.

## Code availability

Code for the pangenome-based DNA methylation framework is available at https://github.com/tianjie16/HPRC2_methylation_framework. Code for PanCGI is available at https://github.com/tianjie16/PanCGI. Code used for figure generation is available at https://github.com/tianjie16/HPRC2_DNA_methylome_variation.

## Acknowledgements

We thank colleagues in the Human Pangenome Reference Consortium for helpful discussions and feedback. This work was supported by NIH grants U41HG010972, U41HG010971, U24HG012070, U24NS132103, and UM1DA058219.

## Author contributions

T.W. and T.L. conceived and designed the study. J.L. and E.E.E. generated and curated the insertion and deletion call sets; D.P. and E.E.E. generated and curated the inversion call set; and X.Z. generated and curated the MEI call set. C.T., E.B. and J.F.M.-V. performed data preprocessing and curation. W.Z. developed the graph-based CpG genotyping method and software. T.L. designed the analytical frameworks, including T2T-CHM13 interval projection for haplotype-specific methylation and CpG-density measurement, HVMR and HVDR identification, models of genetic diversity and SV-associated methylation, PanCGI, CGI genealogy analysis, transcript identification, allele-aware molecular-QTL mapping and fine-mapping, molecular-QTL colocalization and colocalization with GWAS signals, and performed the formal analyses. T.L. generated the visualizations and validated and interpreted the results. The Human Pangenome Reference Consortium generated and provided the HPRC2 haplotype-resolved assemblies and matched long-read molecular datasets used in this study. T.L. wrote the original draft. T.W., T.L., J.F.M.-V., D.L., Z.D., S.D. and E.E.E. reviewed and edited the manuscript. T.W. supervised and administered the project and acquired funding. All authors reviewed and approved the manuscript.

## Human Pangenome Reference Consortium

Julian K. Lucas^7^, Prajna Hebbar^7^, Wen-Wei Liao^8,9^, Juan F. Macias-Velasco^1,2,3^, Adam M. Novak^7^, Mobin Asri^7^, Jennifer R. Balacco^10^, Andrew P. Blair^7^, Davide Bolognini^11^, Jana Ebler^12,13^, Joshua M.V. Gardner^7^, Margarita Geleta^14,15^, Cristian Groza^16^, Andrea Guarracino^17,18^, Peter Heringer^12,19,20^, Glenn Hickey^7^, Sergey Koren^21^, Shuangjia Lu^8,9^, Maximillian G. Marin^22,23^, Christopher Markovic^2^, Mira Mastoras^7^, Capucine Mayoud^24^, Brandy McNulty^7^, Julian M. Menendez^7^, Anna Minkina^25^, Saswat K. Mohanty^26^, Jean Monlong^27^, Katherine M. Munson^4^, Keisuke K. Oshima^28^, David Porubsky^4,6^, T. Rhyker Ranallo-Benavidez^17^, Alessandro Raveane^11^, William E. Seligmann^29^, Ruhollah Shemirani^30^, Yoshihiko Suzuki^31^, Jack A.S. Tierney^32^, Ivo Violich^7^, DongAhn Yoo^4^, Xiaoyu Zhuo^1^, Derek Albracht^2^, Ivan A. Alexandrov^33^, Jamie Allen^32^, Alawi A. Alsheikh-Ali^34^, Casey Andrews^1^, Dmitry Antipov^21^, Lucinda Antonacci-Fulton^2^, Alexander Arguello^35^, Marcelo Ayllon^4^, Edward A. Belter Jr^2^, Halle D. Bender^7^, Katherine E. Bonini^30^, Silvia Buonaiuto^18^, Shuo Cao^18^, Ann M. Mc Cartney^36^, Pi-Chuan Chang^37^, Xian Chang^7^, Jitender Cheema^32^, Claudio Ciofi^38^, Hiram Clawson^7^, Sarah Cody^2^, Vincenza Colonna^18^, Holland C. Conwell^29^, Mark Diekhans^7^, Maria Angela Diroma^38^, Zheng Dong^1^, Danilo Dubocanin^39^, Jordan M. Eizenga^7^, Parsa Eskandar^7^, Eddie Ferro^40^, Sarah M. Ford^29^, Willard W. Ford^41^, Adam Frankish^32^, Mallory A. Freeberg^32^, Qichen Fu^1^, Shenghan Gao^28^, Yan Gao^22^, Gage H. Garcia^4^, Obed A. Garcia^42^, John E. Garza^2^, Mohammadmersad Ghorbani^43^, Tina A. Graves-Lindsay^2^, Bida Gu^44^, Leanne Haggerty^32^, Nancy F. Hansen^21^, Yue Hao^17^, Todd L. Hillaker^7^, S. Nakib Hossain^32^, Neng Huang^22,23^, Sarah E. Hunt^32^, Toby Hunt^32^, Nafiseh Jafarzadeh^7^, Nivesh Jain^10^, Maryam Jehangir^17^, Juan Jiang^1^, Juhyun Kim^21^, Bonhwang Koo^10^, Milinn Kremitzki^1,2^, Daofeng Li^1^, Ronghan Li^1^, Jiadong Lin^4^, Tianjie Liu^1^, Ryan Lorig-Roach^7^, Hailey Loucks^7^, Jane E. Loveland^32^, Jianguo Lu^45^, Walfred Ma^44^, Franco L. Marsico^18^, Jack A. Medico^10^, Younes Mokrab^43,46,47^, Shabir Moosa^43^, Avelina Moreno-Ochando^48,49^, Shinichi Morishita^31^, Jonathan M. Mudge^32^, Njagi Mwaniki^50^, Nasna Nassir^34^, Chiara Natali^38^, Shloka Negi^7^, Lingbin Ni^4^, Faith Okamoto^7^, Chie Owa^31^, Sadye Paez^10^, Clelia Peano^11,51^, Brandon D. Pickett^21^, Laura Pignata^18^, Timofey Prodanov^12,13^, Anandi Radhakrishnan^7^, Brian J. Raney^7^, Andreas Rechtsteiner^52^, Luyao Ren^4,53^, Fedor Ryabov^54,55^, Samuel Sacco^29^, Farnaz Salehi^18^, Aarushi Sehgal^41^, Mahsa Shabani^56^, Shadi Shahatit^24^, Vikram S. Shivakumar^57^, Swati Sinha^32^, Linnéa Smeds^26^, Steven J. Solar^21^, Marco Sollitto^10,38^, Nicole Soranzo^11,58,59^, Marie-Marthe Suner^32^, Arda Söylev^12,13^, Chad Tomlinson^2^, Francesca Floriana Tricomi^32^, Matteo Tommaso Ungaro^29,60^, Rahul Varki^40^, Brian P. Walenz^21^, Charles Wang^61^, Lisa E. Wang^30^, Aaron M. Wenger^62^, Conor V. Whelan^10^, Zilan Xin^1^, Zheng Xu^1^, Wenjin Zhang^1^, Ying Zhou^22^, Giulia Zunino^11^, Nicolas Altemose^39^, Floris P. Barthel^17^, Christina Boucher^40^, Guillaume Bourque^63,64,65^, Andrew Carroll^37^, Monika Cechova^7^, Mark J.P. Chaisson^44^, Haoyu Cheng^66^, Robert Cook-Deegan^67^, Daniel Doerr^12,19,20^, Richard Durbin^58,68^, Anna-Sophie Fiston-Lavier^24,69^, Giulio Formenti^10^, Stephanie M. Fullerton^70^, Robert S. Fulton^2^, Shilpa Garg^71^, Nanibaa’ A. Garrison^72,73,74^, Richard E. Green^29^, Carol W. Greider^52^, Melissa Gymrek^75^, Maximilian Haeussler^7^, Mohammad Amiruddin Hashmi^34^, David Haussler^7^, Alexander G. Ioannidis^7,15,39^, Charles H. Langley^76^, Ben Langmead^57^, Heather A. Lawson^1^, Glennis A. Logsdon^28^, Kateryna D. Makova^77^, Fergal J. Martin^32^, Matthew W. Mitchell^78^, Pilar N. Ossorio^79,80^, Nadia Pisanti^50^, Pjotr Prins^18^, Mikko Rautiainen^81^, Arang Rhie^21^, Michael C. Schatz^57,82^, Laura B. Scheinfeldt^83^, Kishwar Shafin^37^, Jouni Sirén^7^, Andrew B. Stergachis^4,25^, Ahmad Abou Tayoun^84,85^, Mohammed Uddin^34,86^, Flavia Villani^18^, Mitchell R. Vollger^87^, Kai Ye^88^, Evan E. Eichler^4,53^, Erik Garrison^18^, Ira M. Hall^8,9^, Erich D. Jarvis^10,53^, Eimear E. Kenny^30^, Heng Li^22,23^, Jonathan LoTempio Jr^36,89^, Tobias Marschall^12,13^, Karen H. Miga^7^, Adam M. Phillippy^21,57,90,91^, Ting Wang^1,2,3^, Benedict Paten^7^.

^7^UC Santa Cruz Genomics Institute, University of California, Santa Cruz, CA 95060, USA

^8^Center for Genomic Health, Yale University School of Medicine, New Haven, CT 06510, USA

^9^Department of Genetics, Yale University School of Medicine, New Haven, CT 06510, USA

^10^The Vertebrate Genome Laboratory, The Rockefeller University, New York, NY 10065, USA

^11^Human Technopole, Milan, Italy

^12^Center for Digital Medicine, Heinrich Heine University Düsseldorf, Düsseldorf, NRW, DE

^13^Institute for Medical Biometry and Bioinformatics, Medical Faculty and University Hospital Düsseldorf, Heinrich Heine University, Düsseldorf, NRW, DE

^14^Department of Electrical Engineering and Computer Science, University of California, Berkeley, Berkeley, CA 94720, USA

^15^Department of Biomedical Data Science, Stanford University School of Medicine, Stanford, CA 94305, USA

^16^Montreal Heart Institute, Montréal, QC, Canada

^17^Bioinnovation and Genome Sciences, The Translational Genomics Research Institute (TGen), Phoenix, AZ 85004, USA

^18^Department of Genetics, Genomics and Informatics, University of Tennessee Health Science Center, Memphis, TN 38163, USA

^19^Department for Endocrinology and Diabetology at the Medical Faculty and University Hospital Düsseldorf, Heinrich Heine University Düsseldorf, Düsseldorf, NRW, DE

^20^Paul-Langerhans-Group Computational Diabetology, German Diabetes Center (DDZ) and Leibniz Institute for Diabetes Research, Düsseldorf, NRW, DE

^21^Genome Informatics Section, Center for Genomics and Data Science Research, National Human Genome Research Institute, National Institutes of Health, Bethesda, MD 20892, USA

^22^Department of Data Science, Dana-Farber Cancer Institute, Boston, MA 02215, USA

^23^Department of Biomedical Informatics, Harvard Medical School, Boston, MA 02115, USA

^24^ISEM, Univ Montpellier, CNRS, IRD, Montpellier, FR

^25^Division of Medical Genetics, Department of Medicine, University of Washington School of Medicine, Seattle, WA 98195, USA

^26^Department of Biology, Penn State University, University Park, PA 16802, USA

^27^IRSD - Digestive Health Research Institute, University of Toulouse, INSERM, INRAE, ENVT, UPS, Toulouse, FR

^28^Department of Genetics, Epigenetics Institute, Perelman School of Medicine, University of Pennsylvania, Philadelphia, PA 19104, USA

^29^Department of Ecology and Evolutionary Biology, University of California, Santa Cruz, CA 95060, USA

^30^Institute for Genomic Health, Icahn School of Medicine at Mount Sinai, New York, NY 10029, USA

^31^Department of Computational Biology and Medical Sciences, The University of Tokyo, Kashiwa, Chiba 277-8561, Japan

^32^European Molecular Biology Laboratory, European Bioinformatics Institute (EMBL-EBI), Wellcome Genome Campus, Hinxton, Cambridge CB10 1SD, UK

^33^Department of Human Molecular Genetics and Biochemistry, Faculty of Medical and Health Sciences, Tel Aviv University, Tel Aviv 69978, Israel

^34^Center for Applied and Translational Genomics (CATG), Mohammed Bin Rashid University of Medicine and Health Sciences, Dubai Health, Dubai, UAE

^35^Division of Genome Sciences, National Human Genome Research Institute, Bethesda, MD 20871, USA

^36^Institute of Clinical and Translational Sciences, University of California, Irvine, CA 92697, USA

^37^Google LLC, Mountain View, CA 94043, USA

^38^Department of Biology, University of Florence, Sesto Fiorentino, FI 50019, Italy

^39^Department of Genetics, Stanford University, Palo Alto, CA 94304, USA

^40^Department of Computer and Information Science and Engineering, University of Florida, Gainesville, FL 32611, USA

^41^Department of Computer Science and Engineering, University of California San Diego, La Jolla, CA 92093, USA

^42^Department of Anthropology, University of Kansas, Lawrence, KS 66045, USA

^43^Medical and Population Genomics Lab, Sidra Medicine, Doha, Qatar

^44^Quantitative and Computational Biology, University of Southern California, Los Angeles, CA 90089, USA

^45^Sun Yat-sen University, Guangzhou, China

^46^Department of Biomedical Science, College of Health Sciences, Qatar University, Doha, Qatar

^47^Department of Genetic Medicine, Weill Cornell Medicine-Qatar, Doha, Qatar

^48^MATCH biosystems, S.L., Elche, Spain

^49^Universidad Miguel Hernández de Elche, Elche, Spain

^50^Department of Computer Science, University of Pisa, Pisa, Italy

^51^Institute of Genetics and Biomedical Research, UoS of Milan, National Research Council, Milan, Italy

^52^Department of Molecular Cell and Developmental Biology, University of California, Santa Cruz, CA, USA

^53^Howard Hughes Medical Institute, Chevy Chase, MD 20815, USA

^54^The Center for Bio- and Medical Technologies, Moscow, RUS

^55^Centre for Biomedical Research and Technology, HSE University, Moscow, RUS

^56^University of Amsterdam, Amsterdam, Netherlands

^57^Department of Computer Science, Johns Hopkins University, Baltimore, MD 21218, USA

^58^Wellcome Sanger Institute, Genome Campus, Hinxton, CB10 1RQ, UK

^59^School of Clinical Medicine, University of Cambridge, Cambridge, CB2 0SP, UK 60Department of Biology and Biotechnologies “Charles Darwin”, University of Rome “La Sapienza”, Rome 00185, IT

^61^Center for Genomics, Loma Linda University School of Medicine, Loma Linda, CA 92350, USA

^62^PacBio, Menlo Park, CA 94025, USA

^63^Canadian Center for Computational Genomics, McGill University, Montréal, QC H3A 0G1, Canada

^64^Department of Human Genetics, McGill University, Montréal, QC H3A 0G1, Canada

^65^Victor Phillip Dahdaleh Institute of Genomic Medicine, Montréal, QC H3A 0G1, Canada

^66^Department of Biomedical Informatics and Data Science, Yale School of Medicine, New Haven, CT 06510, USA

^67^Arizona State University, Consortium for Science, Policy & Outcomes, Washington, DC 20006, USA

^68^Department of Genetics, University of Cambridge, Cambridge, CB2 3EH, UK

^69^Institut Universitaire de France, Paris, FR

^70^Department of Bioethics & Humanities, University of Washington School of Medicine, Seattle, WA 98195, USA

^71^School of Health Sciences, University of Manchester, Manchester M13 9PL, UK

^72^Traditional, ancestral and unceded territory of the Gabrielino/Tongva peoples, Institute for Society & Genetics, University of California, Los Angeles, Los Angeles, CA 90095, USA

^73^Traditional, ancestral and unceded territory of the Gabrielino/Tongva peoples, Institute for Precision Health, David Geffen School of Medicine, University of California, Los Angeles, Los Angeles, CA 90095, USA

^74^Traditional, ancestral and unceded territory of the Gabrielino/Tongva peoples, Division of General Internal Medicine & Health Services Research, David Geffen School of Medicine, University of California, Los Angeles, Los Angeles, CA 90095, USA

^75^Department of Pediatrics, University of California San Diego, La Jolla, CA 92093, USA

^76^Department of Evolution and Ecology and the Center for Population Biology, University of California, One Shields, Davis, CA 95616, USA

^77^Department of Biology and Center for Medical Genomics, Penn State University, University Park, PA 16802, USA

^78^The Jackson Laboratory for Genomic Medicine, Farmington, CT 06032, USA

^79^Law School, University of Wisconsin-Madison, Madison, WI 53706, USA

^80^Morgridge Institute for Research, Madison, WI 53715, USA

^81^Institute for Molecular Medicine Finland, Helsinki Institute of Life Science, University of Helsinki, Helsinki, Finland

^82^Department of Biology, Johns Hopkins University, Baltimore, MD 21218, USA

^83^Coriell Institute for Medical Research, Camden, NJ 08103, USA

^84^Center for Genomic Discovery, Mohammed Bin Rashid University, Dubai Health, UAE

^85^Dubai Health Genomic Medicine Center, Dubai Health, UAE

^86^GenomeArc Inc, Mississauga, ON, Canada

^87^Department of Human Genetics and Utah Center for Genetic Discovery, University of Utah, Salt Lake City, UT, USA

^88^The first affiliated hospital of Xi’an Jiaotong University, Xi’an Jiaotong University, Xi’an, Shaanxi, 710049, China

^89^Department of Pediatrics, Division of Genetics, School of Medicine, University of California, Irvine, CA 92697, USA

^90^Department of Biomedical Engineering, Johns Hopkins University, Baltimore, MD 21218, USA

^91^Department of Genetic Medicine, Johns Hopkins University School of Medicine, Baltimore, MD 21205, USA

