## Extended Data Figures for "The genetic architecture of human DNA methylome variation"

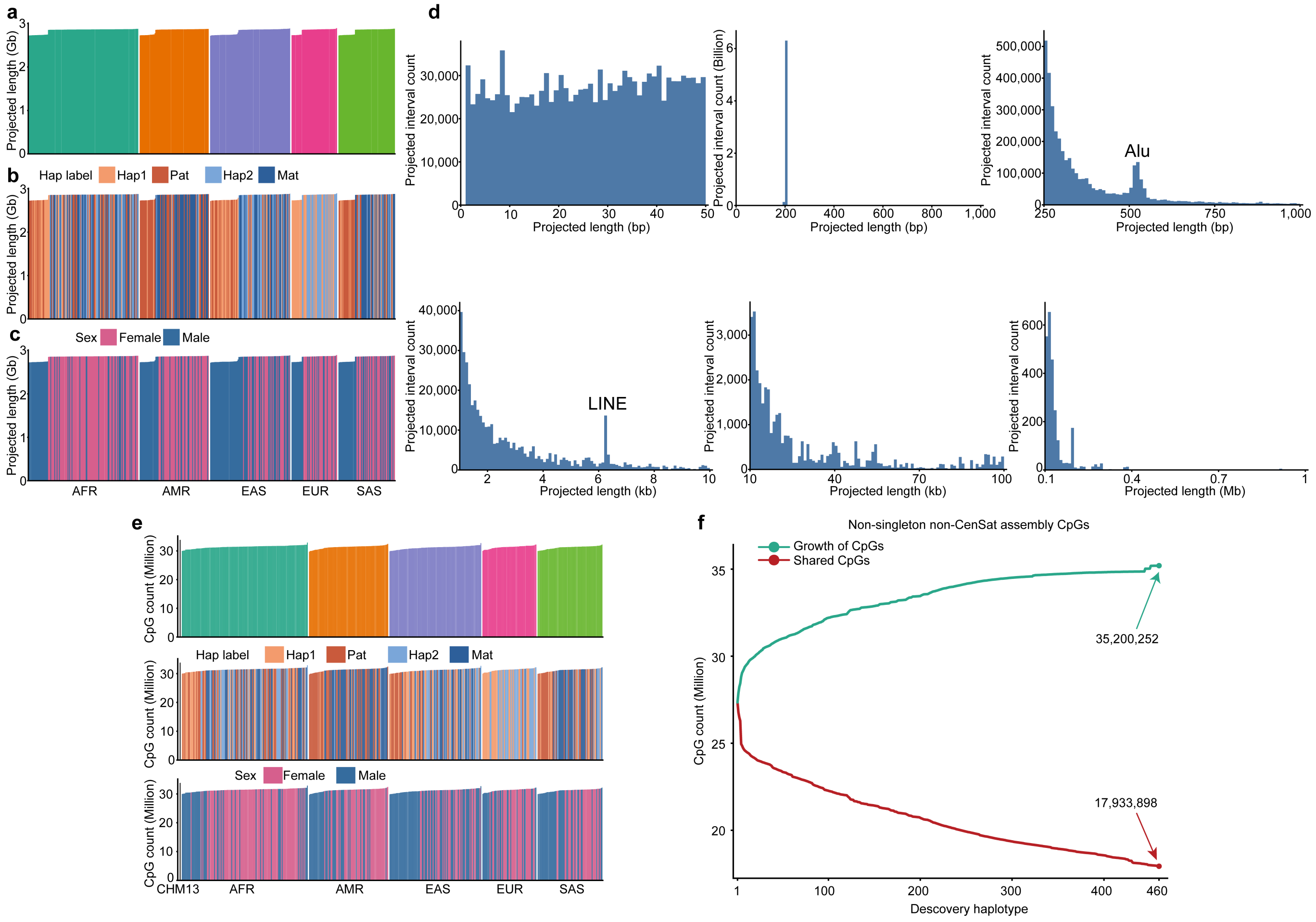


**Extended Data Fig. 1. Pangenome projection performance and CpG sharing across HPRC2 haplotypes.**

a–c, Total T2T-CHM13 sequence recovered by projection from each of 460 haplotypes, ordered by African (AFR), Admixed American (AMR), East Asian (EAS), European (EUR) and South Asian (SAS) superpopulation, and coloured by superpopulation (a), haplotype label (b) or donor sex (c). Each bar denotes one haplotype; median recovery was 2.86 Gb. d, Projected-interval length across six displayed ranges: 0–50 bp, 0–1 kb, 250–1,000 bp, 1–10 kb, 10–100 kb and 0.1–1 Mb. The ranges have different bin widths and their counts are not directly comparable. Overall, 97.27% of projected intervals remained close to 200 bp, whereas longer intervals showed secondary peaks at lengths characteristic of Alu and L1 elements, consistent with the retention of haplotype-specific inserted sequence. e, Graph-CpG counts for T2T-CHM13 and 462 haplotype assemblies, including both HG002 haplotypes, ordered and coloured as in a–c. f, Cumulative graph-CpG discovery across the 460 HPRC2 haplotypes in the same fixed haplotype order. CpGs within each haplotype's own centromere/satellite repeat annotation (CenSat) and graph-CpG identifiers observed on only one HPRC2 haplotype (singletons) were excluded. The *x* axis gives the cumulative number of haplotypes included; green shows cumulative distinct CpGs and red shows CpGs shared by all haplotypes included at each cohort size. Across the 460 HPRC2 haplotypes, 35,200,252 distinct CpGs were observed, of which 17,933,898 were present on all haplotypes; T2T-CHM13 contains 33,839,747 CpGs.

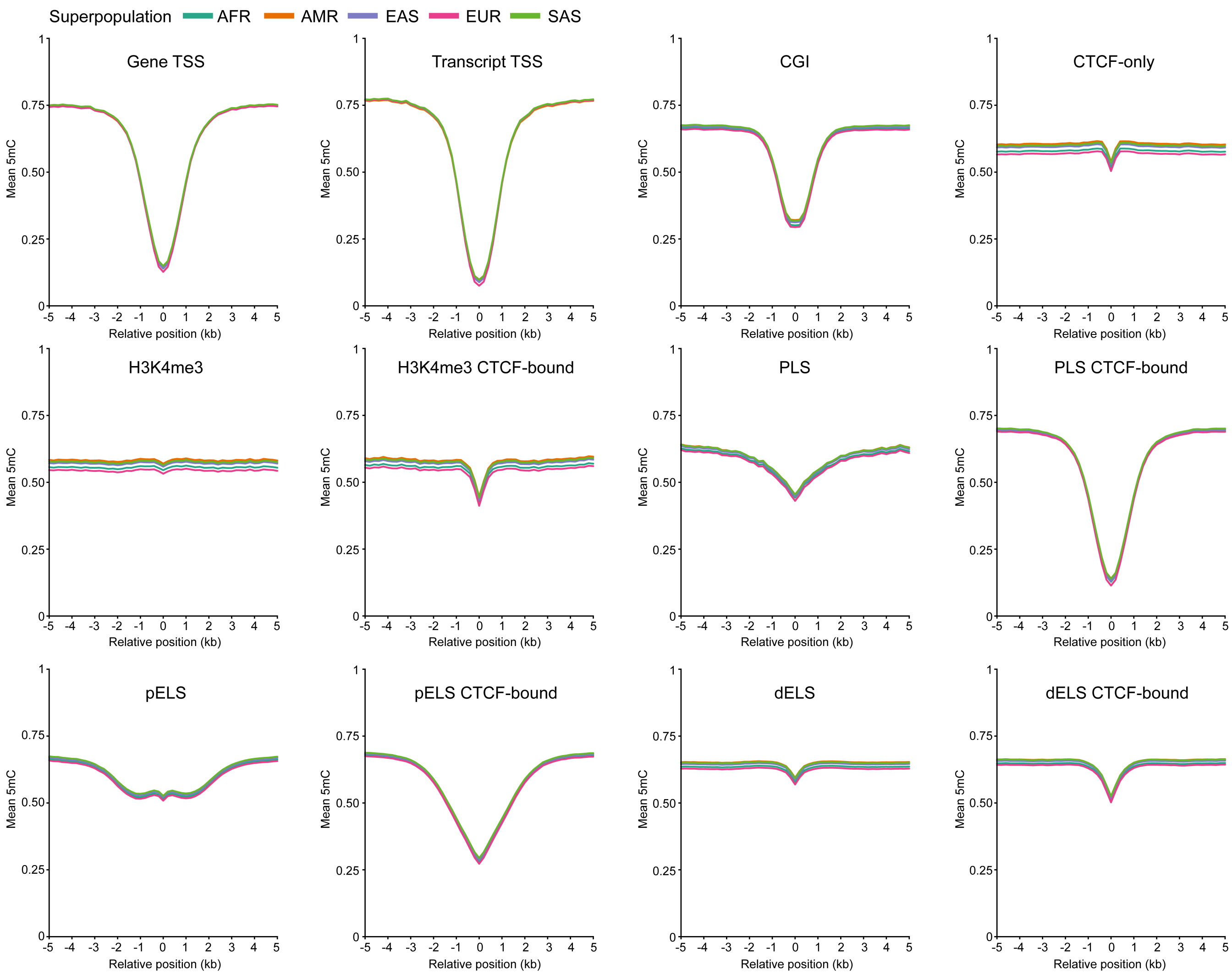


**Extended Data Fig. 2. Superpopulation methylation profiles at regulatory features.**

Mean 5-methylcytosine (5mC) levels in haplotype-specific intervals projected from non-overlapping 200-bp T2T-CHM13 intervals spanning 5 kb on either side of each feature reference point. Gene and transcript panels are aligned to transcription start sites (TSSs), whereas CpG-island (CGI) and candidate cis-regulatory element (cCRE) panels are aligned to feature midpoints. For cCRE panels, position 0 denotes the 200-bp T2T-CHM13 interval containing the cCRE midpoint. The *x* axis gives relative position in 200-bp steps and the *y* axis mean 5mC. At each relative position, methylation was first averaged across the corresponding feature-linked intervals within each haplotype and then across haplotypes within each superpopulation. Lines show superpopulation means from 140 African (AFR), 88 Admixed American (AMR), 102 East Asian (EAS), 58 European (EUR) and 72 South Asian (SAS) haplotypes. Feature classes are not mutually exclusive. The twelve panels are arranged from left to right and top to bottom as follows: expressed-gene TSSs ($n$ = 12,744), expressed-transcript TSSs ($n$ = 66,951), CGI midpoints ($n$ = 29,441), CTCF-only cCREs ($n$ = 54,733), DNase-H3K4me3 cCREs without ($n$ = 16,293) or with ($n$ = 8,587) CTCF binding, promoter-like signatures (PLSs) without ($n$ = 7,298) or with ($n$ = 26,313) CTCF binding, proximal enhancer-like signatures (pELSs) without ($n$ = 62,541) or with ($n$ = 75,537) CTCF binding, and distal enhancer-like signatures (dELSs) without ($n$ = 435,587) or with ($n$ = 213,996) CTCF binding.

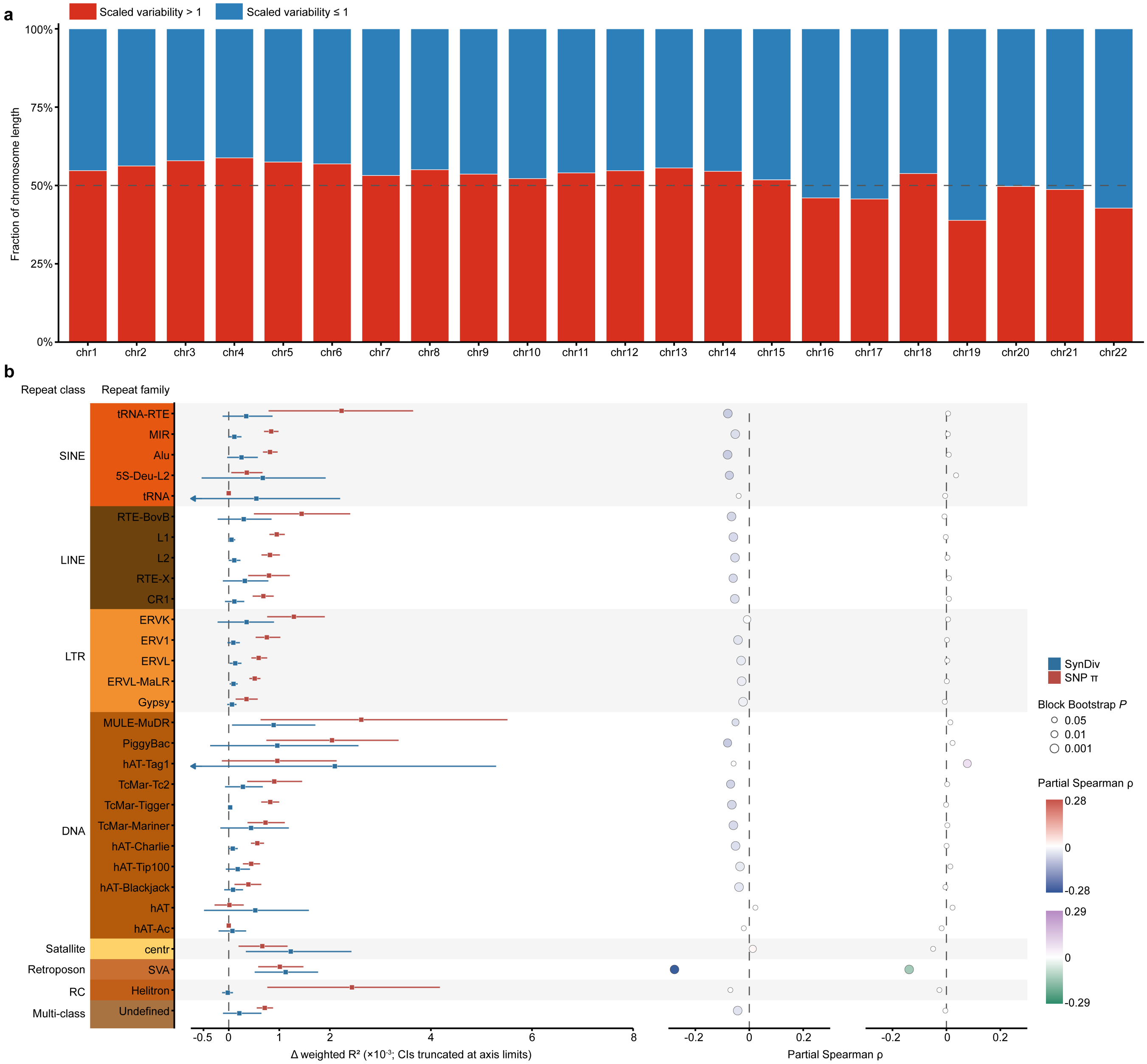


**Extended Data Fig. 3. Chromosomal and repeat-family heterogeneity of methylation variability.**

a, Fraction of analyzed autosomal sequence in 200-bp intervals with scaled methylation variability above 1 (red) or at most 1 (blue); the dashed line marks 50%. Scaled variability is the observed residual variance divided by its mean-dependent expected value. The 9,742,607 analyzed intervals, each with methylation estimates in at least 20% of haplotypes, span

1,948,521,400 bp. The fraction above 1 ranged from 38.9% on chromosome 19 to 58.8% on chromosome 4. This descriptive threshold is not the highly variable methylation region (HVMR) calling criterion. b, Relationships between methylation variability and nucleotide diversity (SNP π) or synteny diversity (SynDiv), which quantifies variation in collinearity among homologous projections, across RepeatMasker families grouped by repeat class. Red and blue squares show the incremental weighted $R^2$ attributable to SNP π and SynDiv, respectively (Δweighted $R^2 \times 10^{-3}$); horizontal lines are 95% intervals from 1,000 chromosome-stratified 5-Mb block-bootstrap replicates and arrows indicate intervals truncated by the plotting range. The middle and right columns show 200-bp partial Spearman correlations; colour gives direction and magnitude and point area the two-sided block-bootstrap *P*. SINE-VNTR-Alu (SVA) showed the strongest inverse correlations among the displayed families for both SNP π (ρ = −0.28) and SynDiv (ρ = −0.14).

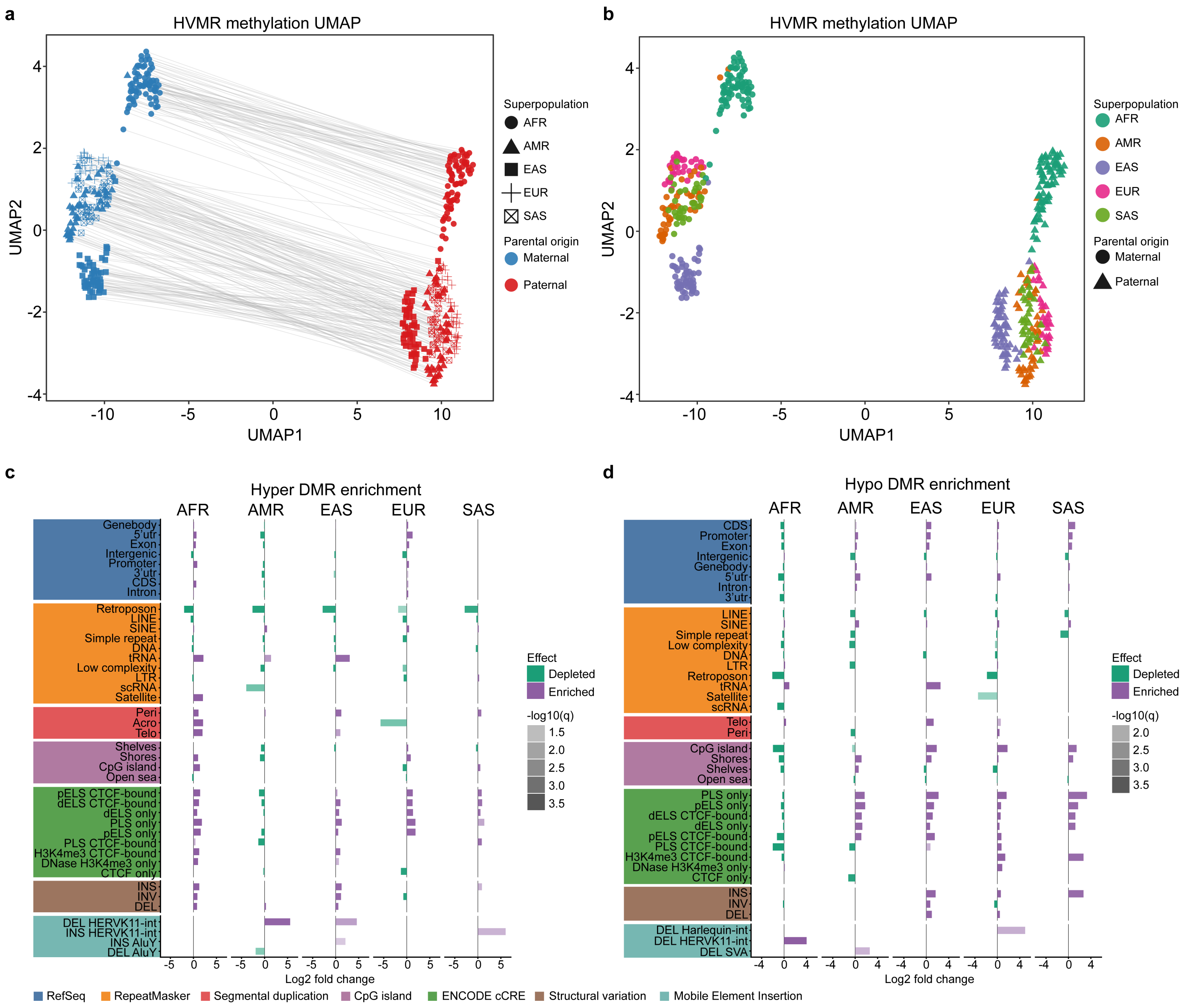


**Extended Data Fig. 4. Haplotype methylation structure and genomic enrichment of population differentially methylated regions (DMRs).**

a,b, Uniform manifold approximation and projection (UMAP) of 460 haplotypes from Euclidean methylation distances across 24,022 highly variable methylation regions (HVMRs). Each point denotes one haplotype, and the axes show UMAP embedding coordinates. In a, colour denotes maternal or paternal assignment, shape denotes superpopulation and grey lines join the two haplotypes of each donor. In b, colour denotes superpopulation and shape parental assignment. Superpopulations are African (AFR), Admixed American (AMR), East Asian (EAS), European

(EUR) and South Asian (SAS). c,d, Genomic-feature enrichment and depletion for hypermethylated (c) and hypomethylated (d) population DMRs. Five superpopulation-versus-remainder comparisons across both methylation directions yielded 217,637 comparison-specific calls: 146,396 for AFR, 30,427 for AMR, 11,970 for EAS, 23,499 for EUR and 5,345 for SAS. Bar length and direction give signed $\log_2$ observed-to-expected nucleotide overlap from Genomic Association Tester (GAT); rightward purple bars indicate enrichment, leftward green bars depletion, and shading gives $-\log_{10}(q)$, where $q$ is the Benjamini–Hochberg-adjusted GAT $P$ value. Displayed terms had $q < 0.05$ and concordant direction in both GAT and GC-stratified regioneR analyses.

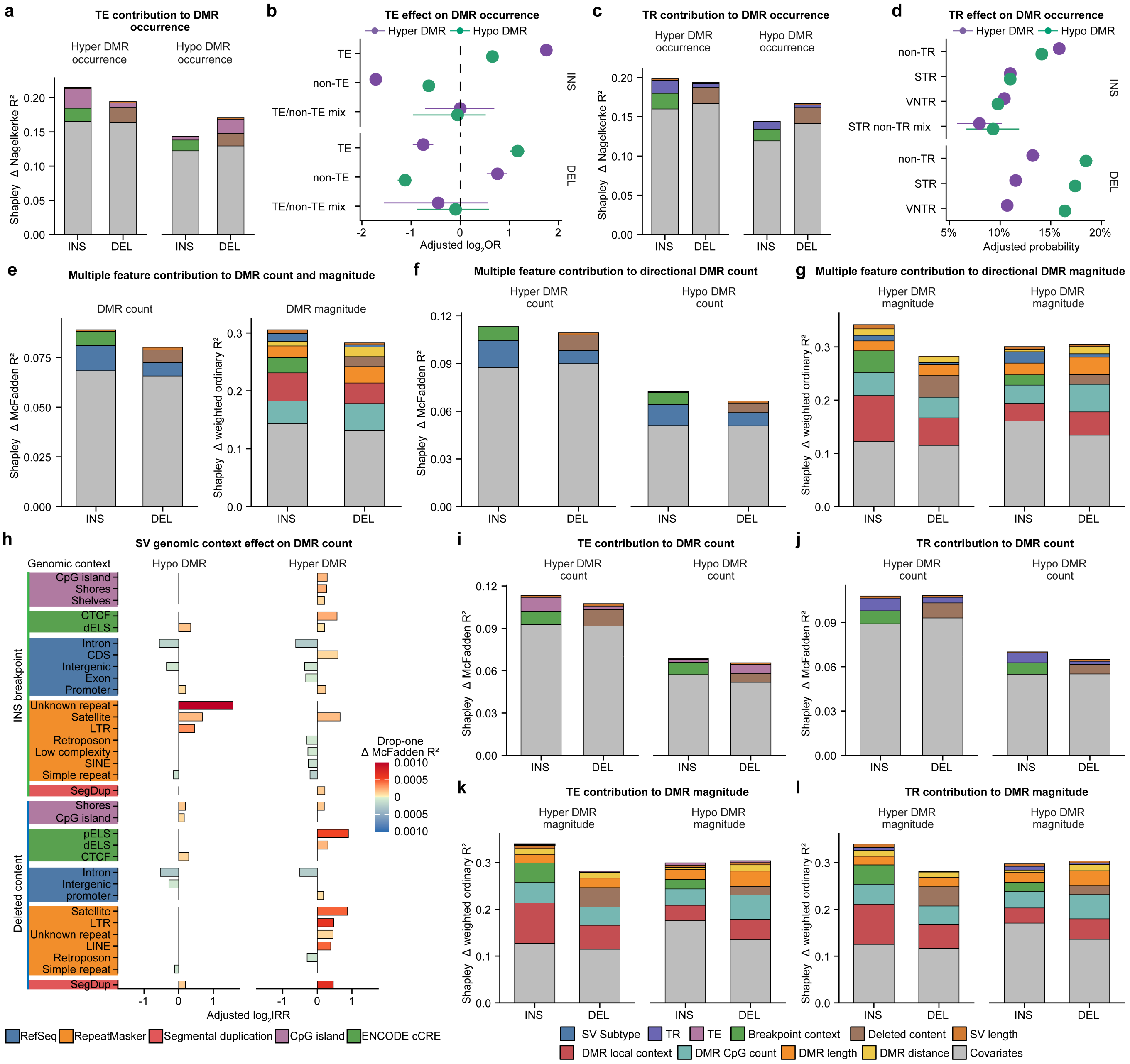


**Extended Data Fig. 5. Relative contributions of structural-variant features to differentially methylated region (DMR) occurrence, count and magnitude.**

a, Shapley decomposition of directional DMR-occurrence models for insertions (INS) and deletions (DEL), using transposable-element (TE) classification. Bars show each predictor group's contribution to the full-model Nagelkerke pseudo-$R^2$. b, Covariate-adjusted TE effects on hypermethylated and hypomethylated DMR occurrence, shown as $\log_2$ odds ratios for TE, non-TE and TE/non-TE mix classes. Purple and green denote hypermethylated and hypomethylated DMRs, respectively. c, Corresponding Shapley decomposition using tandem-repeat (TR) classification. d, Adjusted occurrence probabilities for non-TR, short tandem repeat (STR) and variable-number tandem repeat (VNTR) classes; the STR/non-TR mix is additionally shown for INS. e, Shapley decomposition for DMR count (left) and DMR magnitude (right) in models including DMRs of either methylation direction. DMR count is the number of DMRs associated with a structural variant (SV), whereas DMR magnitude is the absolute carrier-minus-non-carrier methylation difference for each SV-DMR pair. Shapley contributions partition McFadden pseudo-$R^2$ for count and CpG-weighted $R^2$ for DMR magnitude. f,g, Direction-specific Shapley decompositions for DMR count (f) and DMR magnitude (g), using SV subtype as the classification term. Contributions partition McFadden pseudo-$R^2$ for count and CpG-weighted $R^2$ for DMR magnitude. Models included SV subtype, SV length, the genomic context of the insertion breakpoint for INS or the deleted sequence for DEL, and covariates. DMR-magnitude models additionally included DMR-local genomic context, DMR CpG count, DMR length and distance to the nearest breakpoint. h, Genomic-context effects on directional DMR count, shown as adjusted $\log_2$ incidence-rate ratios for insertion-breakpoint context (top) and deleted-sequence context (bottom). Red and blue indicate increased and decreased DMR count, respectively, and colour intensity gives the drop-one ΔMcFadden pseudo-$R^2$. Only terms with Benjamini–Hochberg $q < 0.05$ within an SV class, DMR direction and context role are shown. i,j, Corresponding direction-specific DMR-count decompositions using TE (i) or TR (j) classification in place of SV subtype. k,l, Corresponding DMR-magnitude decompositions using TE (k) or TR (l) classification. Allele frequency and the number of tested CpGs were included as covariates and combined in the grey segment of each stacked decomposition. In b,d, points show bootstrap medians and horizontal lines show 95% intervals from 200 SV-level bootstrap replicates.

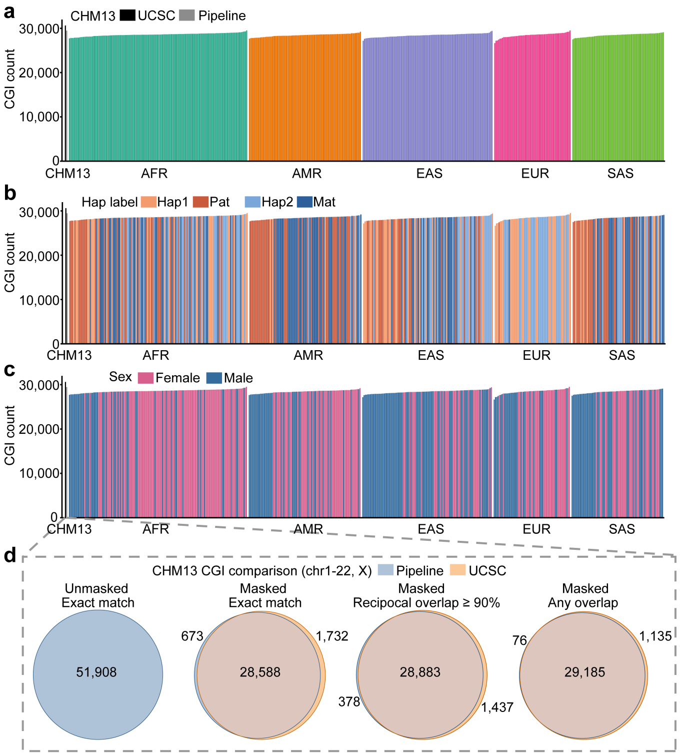


**Extended Data Fig. 6. Benchmarking the pangenome CpG-island catalogue.**

a–c, CpG-island (CGI) counts for 462 HPRC2 haplotype assemblies, including both haplotypes of HG002, grouped by superpopulation and ordered by CGI count within each group. Bars are coloured by superpopulation (a), haplotype label (b) or donor sex (c). The two CHM13 reference bars at left show UCSC and study-pipeline CGI calls. d, CHM13 benchmark restricted to chromosomes 1–22 and X, comparing pipeline calls with the UCSC CGI set under four criteria. The unmasked exact-coordinate comparison contained 51,908 shared calls and no calls unique to either set. After repeat masking, calls were compared by exact coordinates, at least 90% reciprocal overlap and any overlap; numbers indicate shared and method-specific call counts. Exact matching requires identical start and end coordinates; reciprocal overlap requires at least 90% of both intervals to overlap.

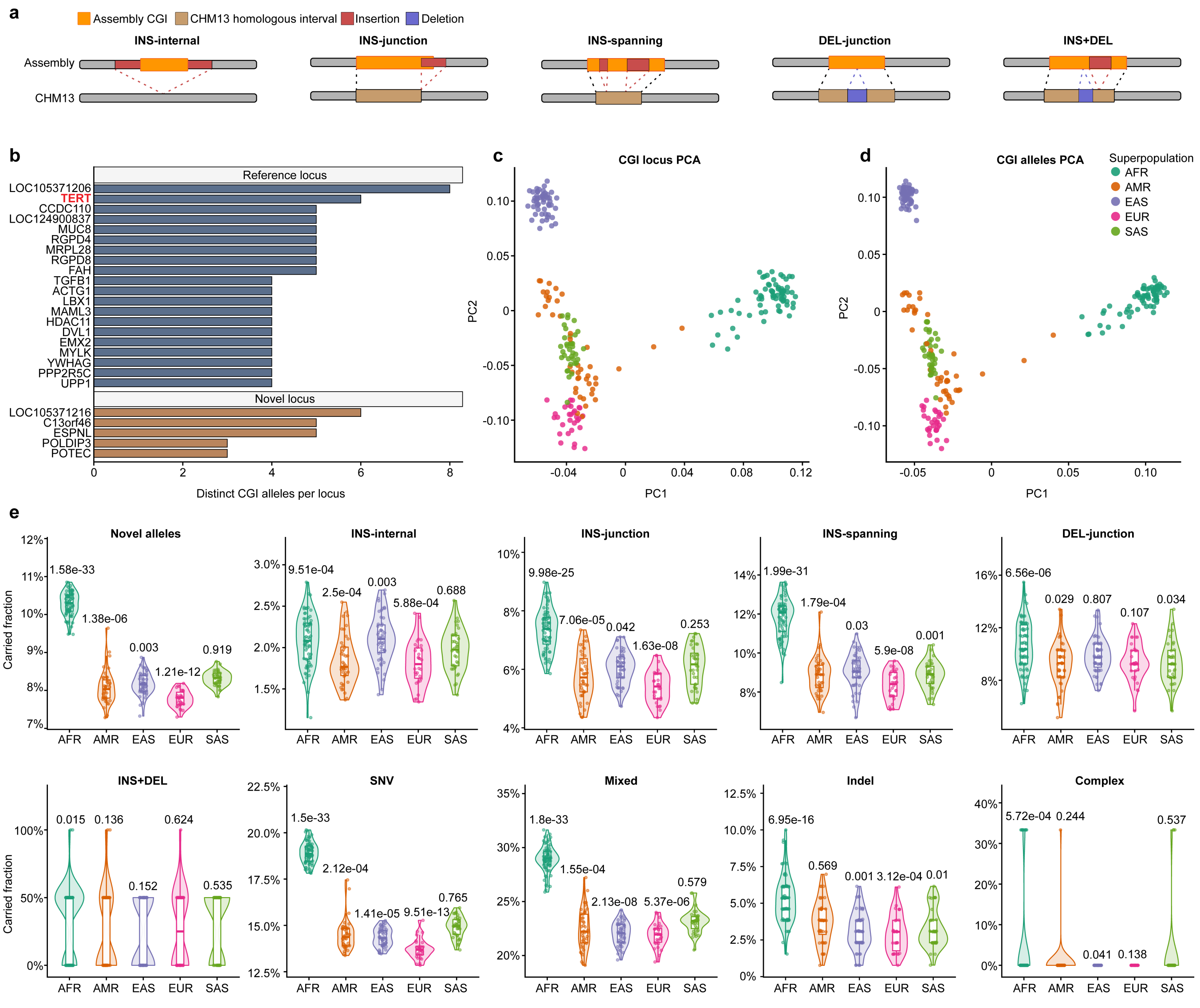


**Extended Data Fig. 7. Sequence origins, population structure and individual carriage of CpG-island alleles.**

a, Representative sequence configurations for novel CpG-island (CGI) alleles associated with insertions (INS) and deletions (DEL): CGIs within inserted sequence (INS-internal), crossing insertion junctions (INS-junction), spanning inserted sequence (INS-spanning), spanning deletion junctions (DEL-junction), or associated with both an insertion and a deletion (INS+DEL). Schematics locate each CGI relative to the structural variant and flanking reference-homologous sequence and define the mutually exclusive sequence-origin classes used in **Fig.**

**3e**. b, Number of distinct CGI alleles at promoter-associated reference and novel loci. *TERT* is highlighted as a reference locus with extensive allelic diversity. c,d, Principal-component analysis (PCA) of CGI presence dosage for 231 individuals, summarized at locus (c) and allele (d) resolution. Points denote individuals and colours denote African (AFR), Admixed American (AMR), East Asian (EAS), European (EUR) and South Asian (SAS) superpopulations. The axes show the first two principal components. e, Fraction of callable CGI alleles in each displayed category carried by an individual. Each point denotes one individual. Labels give raw two-sided Wilcoxon rank-sum *P* values comparing each superpopulation with all remaining individuals within the same panel. Violins show the distributions; boxes show medians and interquartile ranges, and whiskers extend to 1.5 times the interquartile range. Haplotype-level carried fractions are shown in **Supplementary Fig. 7**.

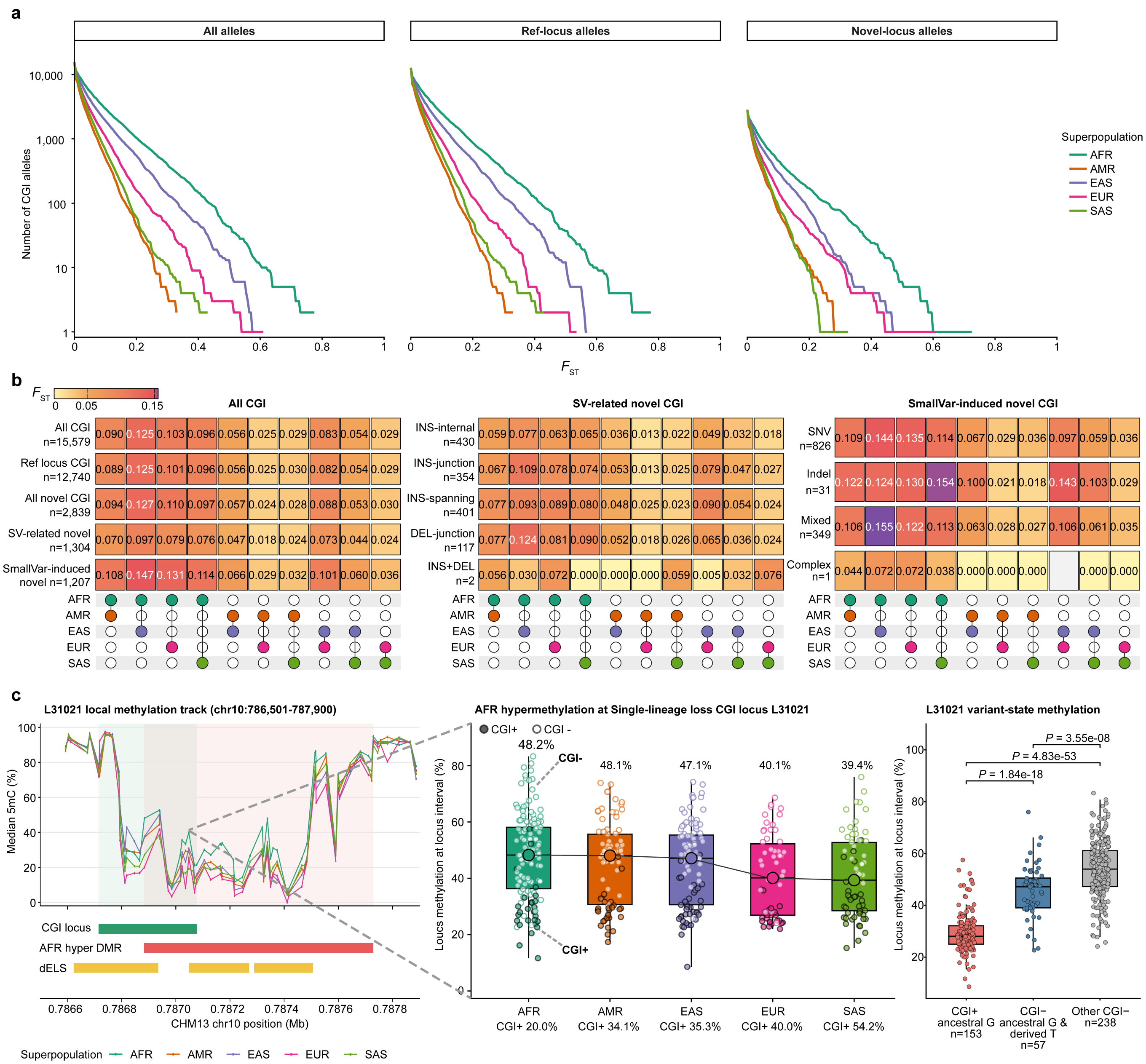


**Extended Data Fig. 8. Population differentiation and methylation of CpG-island states.**

a, Reverse cumulative distributions of Hudson's fixation index ($F_{ST}$) from one-superpopulation-versus-remainder comparisons for 15,579 autosomal polymorphic CpG-island (CGI) alleles after filtering for a global call rate of at least 0.90 and a minor-allele frequency of at least 0.01. These comprised 12,740 reference-locus and 2,839 novel-locus alleles. The *y* axis, shown on a logarithmic scale, gives the number of alleles with $F_{ST}$ at least as large as the value on the *x* axis. $F_{ST}$ was calculated from binary CGI-presence allele counts. b, Pairwise ratio-of-sums Hudson $F_{ST}$ by sequence-origin class and finer mechanism. Each heatmap value is the ratio of Hudson numerator and denominator terms summed across the alleles in that category; the dot matrix identifies the superpopulation pair represented by each column, and labels give the number of alleles. Superpopulations are African (AFR), Admixed American (AMR), East Asian (EAS), European (EUR) and South Asian (SAS). Structural variant (SV), insertion (INS), deletion (DEL), single-nucleotide variant (SNV) and short insertion/deletion (indel) denote the corresponding sequence-origin categories; SmallVar-induced denotes small-variant-induced alleles. c, L31021, a genealogy-supported single-lineage-loss locus overlapping an AFR-hypermethylated differentially methylated region (DMR). Left, superpopulation methylation profiles together with the CGI locus, AFR-hypermethylated DMR and distal enhancer-like signature (dELS) annotations. Middle, haplotype methylation by superpopulation; filled points denote CGI-positive haplotypes and open points denote CGI-negative haplotypes. Labels above the distributions give median locus methylation. Percentages in the *x*-axis labels give the CGI-positive fraction among all 462 haplotypes with a called CGI state (AFR, 20.0%; AMR, 34.1%; EAS, 35.3%; EUR, 40.0%; SAS, 54.2%); methylation distributions comprise the 448 haplotypes with available methylation measurements. Right, methylation across three mutually exclusive sequence states: CGI-positive haplotypes carrying the ancestral G allele ($n$ = 153), CGI-negative haplotypes retaining the ancestral G allele but carrying one or both of two additional C>T substitutions ($n$ = 57), and other CGI-negative haplotypes ($n$ = 238). Median methylation was 28.05%, 47.12% and 53.97%, respectively. Brackets show raw two-sided Wilcoxon rank-sum $P = 1.84 \times 10^{-18}$, $4.83 \times 10^{-53}$ and $3.55 \times 10^{-8}$. Boxes show medians and interquartile ranges; whiskers extend to 1.5 times the interquartile range, and points denote haplotypes with methylation measurements.

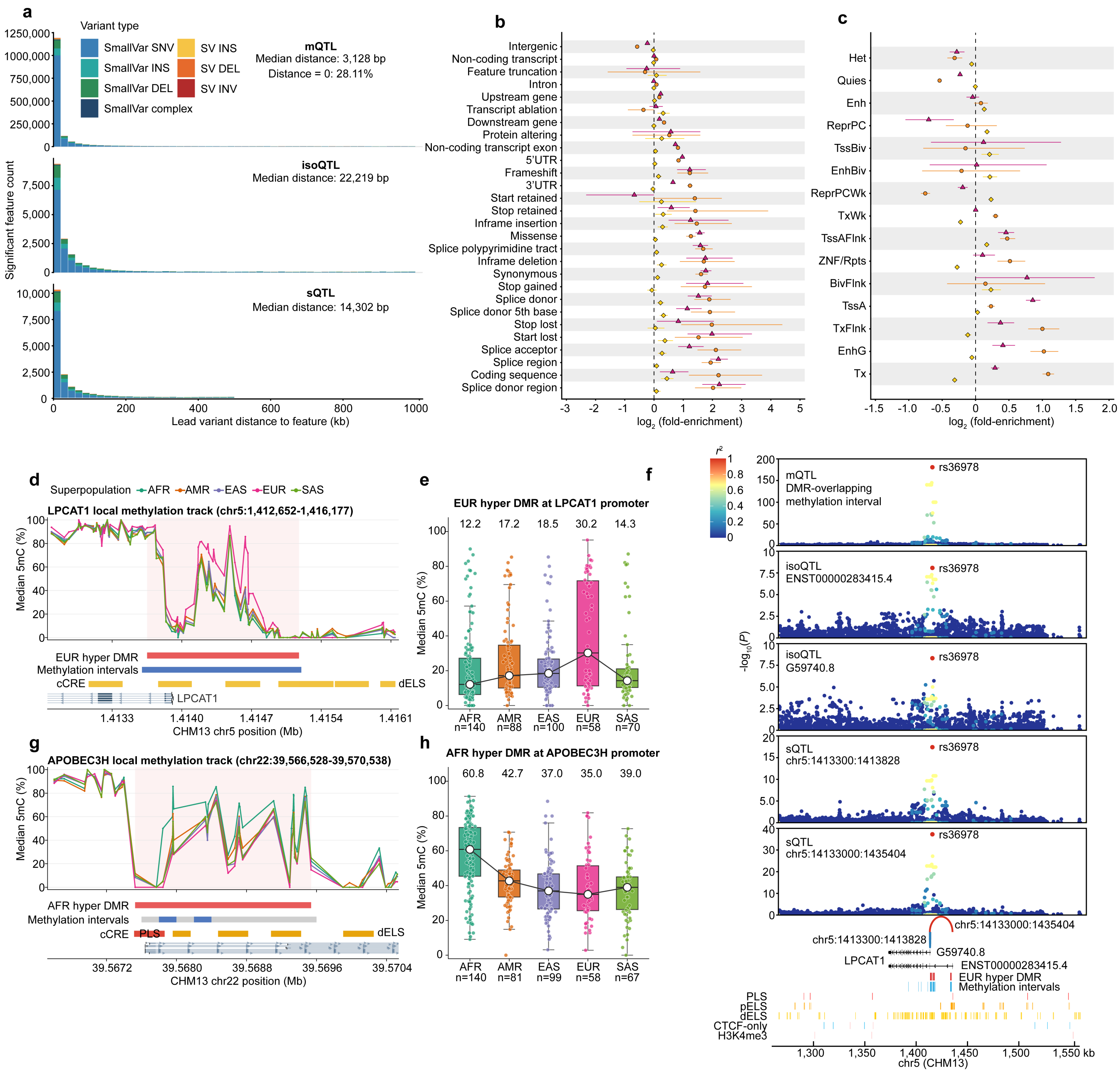


**Extended Data Fig. 9. Genomic and functional context of molecular quantitative trait loci (QTLs).**

a, Absolute distance from each association lead to its corresponding molecular feature, stratified by small-variant and structural-variant leads. For methylation quantitative trait loci (mQTLs), distance was zero when the lead fell within the methylation interval and otherwise was measured to the nearest interval boundary; for isoform quantitative trait loci (isoQTLs), distance was measured to the isoform transcription start site; and for splicing quantitative trait loci (sQTLs), distance was measured to the midpoint of the corresponding splice-junction interval. Across all leads, median distances were 3,128 bp for mQTLs, 22,219 bp for isoQTLs and 14,302 bp for sQTLs. Colours distinguish small-variant leads, comprising single-nucleotide variants (SNVs), small insertions, small deletions and complex variants, from structural-variant (SV) leads, comprising insertions (INS), deletions (DEL) and inversions (INV). b,c, Empirical enrichment of the variant with the highest posterior inclusion probability (PIP) from each independent credible set across Variant Effect Predictor consequences (b) and the GM12878 Roadmap E116 15-state ChromHMM model (c). Yellow diamonds, orange circles and magenta triangles denote mQTLs, isoQTLs and sQTLs, respectively; points show $\log_2$ observed-to-expected overlap and lines show empirical 95% intervals from 1,000 control sets matched for allele frequency, distance to the molecular feature, variant class and, for SVs, length. Coding variants were enriched 1.36-fold for mQTLs and 4.62-fold for isoQTLs; splice-donor-region variants were enriched 4.71-fold for sQTLs. d, Superpopulation median methylation and genomic features across *LPCAT1* on CHM13. Lines show African (AFR), Admixed American (AMR), East Asian (EAS), European (EUR) and South Asian (SAS) superpopulation medians. e, Methylation in the EUR-hypermethylated differentially methylated region (DMR) at the *LPCAT1* promoter across 456 haplotypes. Median methylation was 30.2% in EUR and 12.2–18.5% in the other superpopulations. Each coloured point denotes the median methylation of one haplotype across coverage-filtered CpGs within the DMR. White points show the superpopulation medians and the black line connects these medians in the displayed superpopulation order. Boxes show medians and interquartile ranges; whiskers extend to 1.5 times the interquartile range. f, Association panels at *LPCAT1* for mQTLs, two isoQTLs and two sQTLs. The mQTL panel shows the minimum *P* value for each variant across eight DMR-overlapping methylation intervals. rs36978 is the leading shared variant, is enriched in EUR (allele frequency, 0.466 versus 0.127 in non-EUR haplotypes), and has a positive mQTL effect concordant with the

EUR-hypermethylated DMR. Point colour gives phased-haplotype $r^2$ to rs36978; the variant is marked in red in each association panel. Lower tracks show transcript models, the two colocalized splice junctions, the DMR, methylation intervals and regulatory features. g, Superpopulation median methylation and genomic features across *APOBEC3H* on CHM13, with colours as in d. h, Methylation in the AFR-hypermethylated *APOBEC3H* promoter DMR across 445 haplotypes. Median methylation was 60.8% in AFR and 35.0–42.7% in the other superpopulations. Plot elements are as in e.

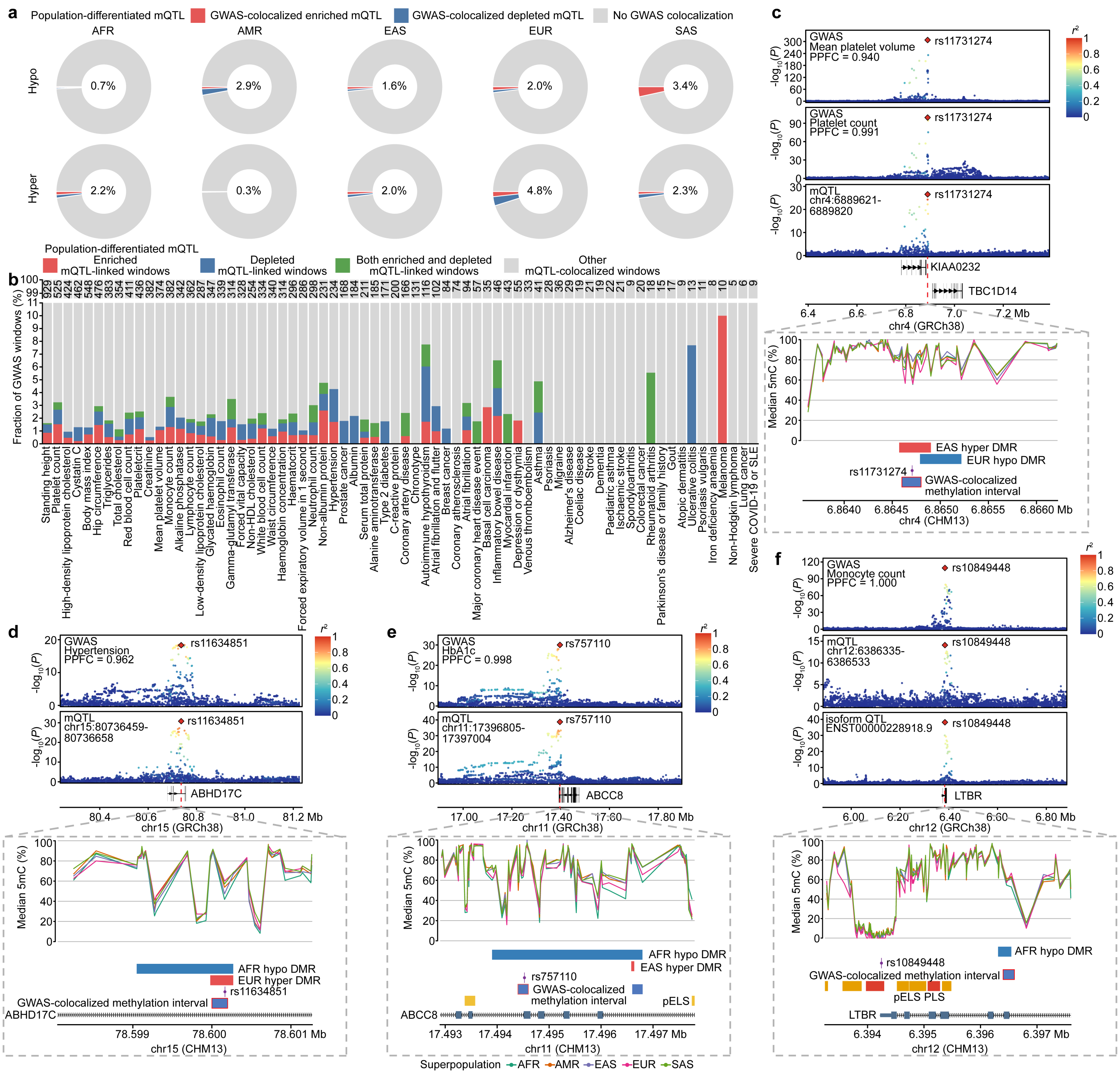


**Extended Data Fig. 10. Population-differentiated methylation quantitative trait loci (mQTLs) at complex-trait loci.**

a, Fraction of population-differentiated mQTL variants that exactly match a HyPrColoc candidate variant, stratified by superpopulation and the direction of the corresponding population differentially methylated region (population DMR). Superpopulations are African (AFR), Admixed American (AMR), East Asian (EAS), European (EUR) and South Asian (SAS). Variants are deduplicated within each panel even when they are associated with multiple methylation intervals or support multiple population DMRs. Among 17,990 variants counted uniquely within each superpopulation, 202 had at least one exact genome-wide association study (GWAS) colocalization (1.12%). Red and blue denote effect alleles enriched in or depleted from the focal superpopulation. b, Fraction of 13,533 independent GWAS windows with an mQTL-containing HyPrColoc result that also contained the same population-differentiated variant-methylation interval pair supporting a population DMR. Red, blue and green denote enriched-only, depleted-only and both enriched-and-depleted population-mQTL support, respectively; grey denotes other mQTL-colocalized windows. In total, 277 windows (2.05%) contained a matching population-differentiated variant-methylation interval pair across 48 of 70 traits: 105 enriched-only, 120 depleted-only and 52 both. Values above bars give eligible windows per trait. c, Mean platelet volume and platelet count at rs11731274 (posterior probability of full colocalization (PPFC) = 0.940 and 0.991, respectively), with an mQTL whose effect allele is enriched in EAS ($\beta = 0.301$, $P = 2.76 \times 10^{-27}$), concordant with an EAS-hypermethylated DMR. d, Hypertension at rs11634851 (PPFC = 0.962), with an mQTL whose effect allele is enriched in EUR ($\beta = 0.378$, $P = 1.64 \times 10^{-31}$). e, Glycated hemoglobin (HbA1c) at rs757110 (PPFC = 0.998), with an mQTL whose effect allele is depleted in AFR ($\beta = 0.153$, $P = 1.86 \times 10^{-39}$). f, Monocyte count at rs10849448 (PPFC = 0.9996), with an mQTL whose effect allele is depleted in AFR ($\beta = 0.278$, $P = 7.85 \times 10^{-15}$) and an isoform quantitative trait locus (isoQTL) for *LTBR* ($\beta = 1.263$, $P = 6.37 \times 10^{-39}$). In c–f, association panels use GRCh38, the common coordinate system used to harmonize GWAS and molecular quantitative trait locus (QTL) variants for HyPrColoc; lower CHM13 panels show superpopulation median methylation together with population DMR, GWAS-colocalized methylation-interval and genomic-feature annotations, where shown. Filled red diamonds mark HyPrColoc candidate variants, point colour gives phased-haplotype $r^2$ to the candidate, blue boxes mark colocalized methylation intervals and red outlines mark the representative interval plotted above. Population allele frequencies and molecular-QTL β values are reported relative to the CHM13 alternate allele. Additional population DMRs and gene labels provide genomic context and do not indicate further population-mQTL matches or target assignments.
