## Supplementary Figures for "The genetic architecture of human DNA methylome variation"

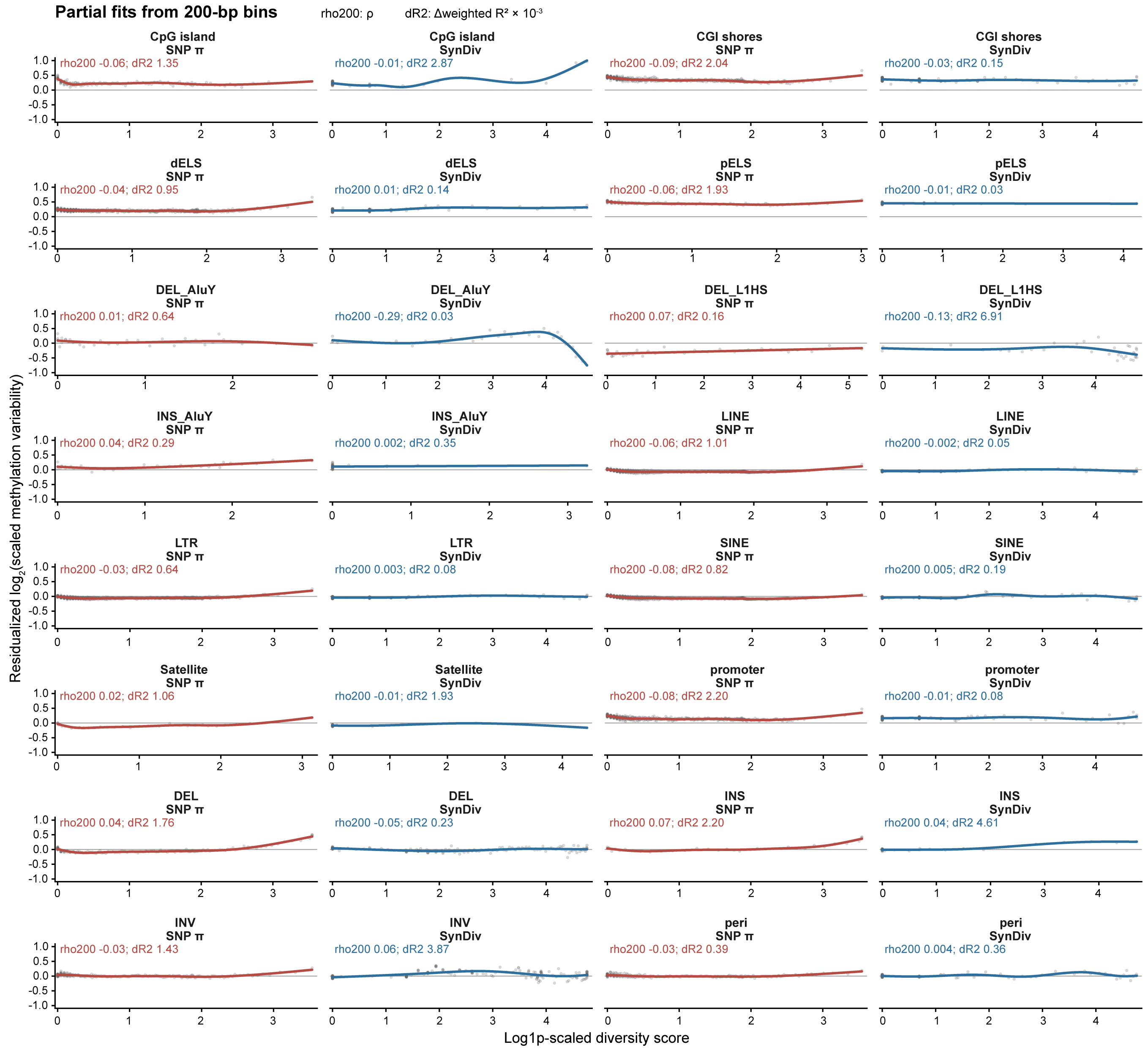


**Supplementary Fig. 1. Methylation variability in relation to nucleotide and synteny diversity across genomic-feature classes.**

Covariate-adjusted relationships between residualized $\log_2$(meVar), where meVar denotes scaled methylation variability, and nucleotide diversity (SNP π) or synteny diversity (SynDiv), which quantifies variation in collinearity among homologous projections, across 16 genomic-feature classes. The analysis contains 9,742,607 200-bp intervals with methylation estimates in at least 20% of haplotypes, of which 6,953,997 overlap at least one displayed class; classes are not mutually exclusive. Grey points are reliability-weighted means in predictor-quantile bins containing at least 50 intervals. Red and blue curves show low-rank cubic regression splines for SNP π and SynDiv, respectively, fitted to the binned summaries. The *x* axis gives ln[1 + diversity/median(non-zero diversity)]. Labels report the 200-bp partial Spearman correlation (rho200) and the incremental weighted $R^2$ attributable to the displayed diversity term (dR2; Δweighted $R^2 \times 10^{-3}$). Five-megabase blocks were used only for the inferential summaries reported in **Fig. 1f**; the displayed curves were fitted to summaries of the 200-bp intervals.

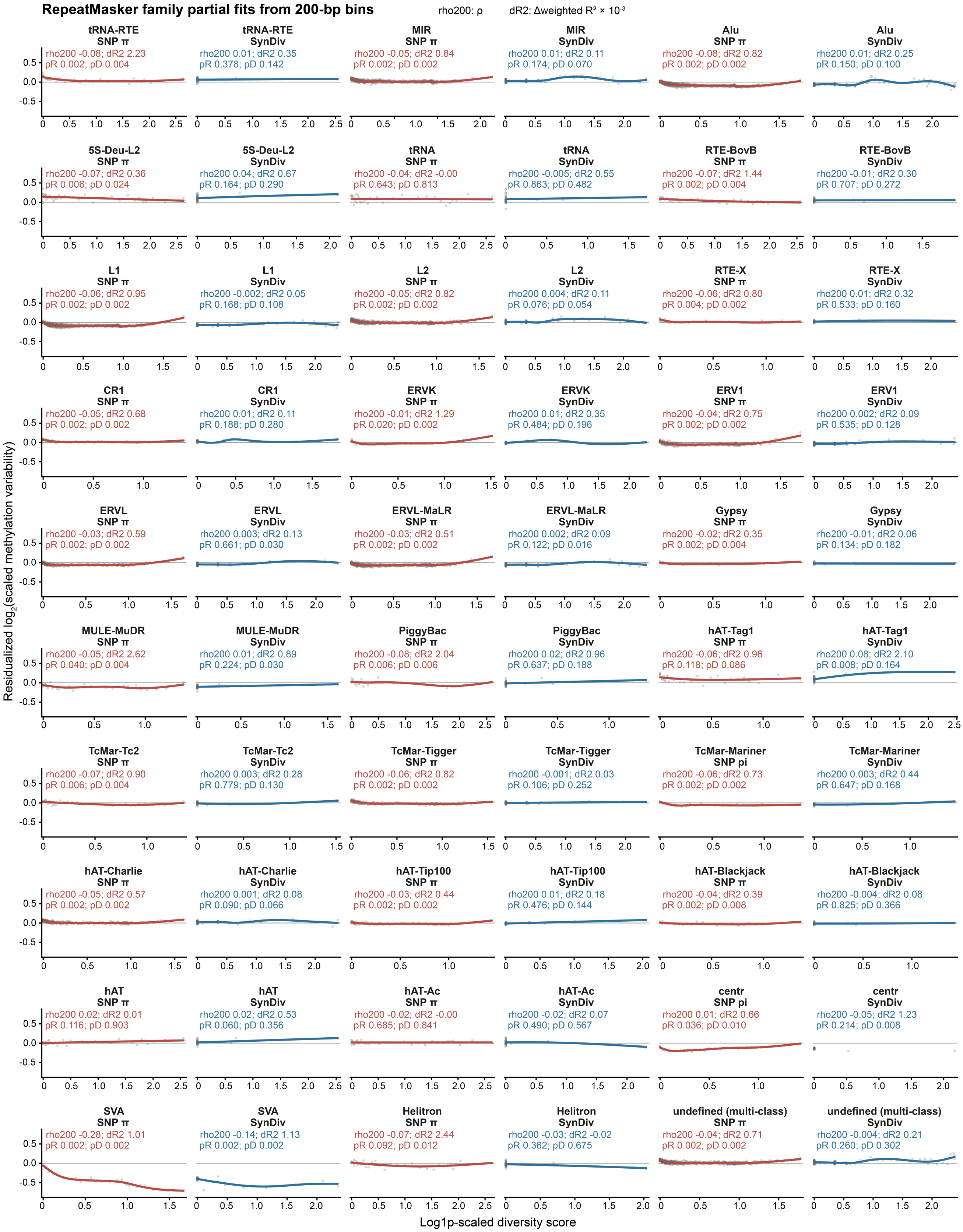
RepeatMasker family partial fits from 200-bp bins
rho200: ρ
dR2: Δweighted R² × 10⁻³
tRNA-RTE SNP π
rho200 -0.08; dR2 2.23
pR 0.002; pD 0.004
tRNA-RTE SynDiv
rho200 0.01; dR2 0.35
pR 0.378; pD 0.142
MIR SNP π
rho200 -0.05; dR2 0.84
pR 0.002; pD 0.002
MIR SynDiv
rho200 0.01; dR2 0.11
pR 0.174; pD 0.070
Alu SNP π
rho200 -0.08; dR2 0.82
pR 0.002; pD 0.002
Alu SynDiv
rho200 0.01; dR2 0.25
pR 0.150; pD 0.100
5S-Deu-L2 SNP π
rho200 -0.07; dR2 0.36
pR 0.006; pD 0.024
5S-Deu-L2 SynDiv
rho200 0.04; dR2 0.67
pR 0.164; pD 0.290
tRNA SNP π
rho200 -0.04; dR2 -0.00
pR 0.643; pD 0.813
tRNA SynDiv
rho200 -0.005; dR2 0.55
pR 0.863; pD 0.482
RTE-BovB SNP π
rho200 -0.07; dR2 1.44
pR 0.002; pD 0.004
RTE-BovB SynDiv
rho200 -0.01; dR2 0.30
pR 0.707; pD 0.272
L1 SNP π
rho200 -0.06; dR2 0.95
pR 0.002; pD 0.002
L1 SynDiv
rho200 -0.002; dR2 0.05
pR 0.168; pD 0.108
L2 SNP π
rho200 -0.05; dR2 0.82
pR 0.002; pD 0.002
L2 SynDiv
rho200 0.004; dR2 0.11
pR 0.076; pD 0.054
RTE-X SNP π
rho200 -0.06; dR2 0.80
pR 0.004; pD 0.002
RTE-X SynDiv
rho200 0.01; dR2 0.32
pR 0.533; pD 0.160
CR1 SNP π
rho200 -0.05; dR2 0.68
pR 0.002; pD 0.002
CR1 SynDiv
rho200 0.01; dR2 0.11
pR 0.188; pD 0.280
ERVK SNP π
rho200 -0.01; dR2 1.29
pR 0.020; pD 0.002
ERVK SynDiv
rho200 0.01; dR2 0.35
pR 0.484; pD 0.196
ERV1 SNP π
rho200 -0.04; dR2 0.75
pR 0.002; pD 0.002
ERV1 SynDiv
rho200 0.002; dR2 0.09
pR 0.535; pD 0.128
ERVL SNP π
rho200 -0.03; dR2 0.59
pR 0.002; pD 0.002
ERVL SynDiv
rho200 0.003; dR2 0.13
pR 0.661; pD 0.030
ERVL-MaLR SNP π
rho200 -0.03; dR2 0.51
pR 0.002; pD 0.002
ERVL-MaLR SynDiv
rho200 0.002; dR2 0.09
pR 0.122; pD 0.016
Gypsy SNP π
rho200 -0.02; dR2 0.35
pR 0.002; pD 0.004
Gypsy SynDiv
rho200 -0.01; dR2 0.06
pR 0.134; pD 0.182
MULE-MuDR SNP π
rho200 -0.05; dR2 2.62
pR 0.040; pD 0.004
MULE-MuDR SynDiv
rho200 0.01; dR2 0.89
pR 0.224; pD 0.030
PiggyBac SNP π
rho200 -0.08; dR2 2.04
pR 0.006; pD 0.006
PiggyBac SynDiv
rho200 0.02; dR2 0.96
pR 0.637; pD 0.188
hAT-Tag1 SNP π
rho200 -0.06; dR2 0.96
pR 0.118; pD 0.086
hAT-Tag1 SynDiv
rho200 0.08; dR2 2.10
pR 0.008; pD 0.164
TcMar-Tc2 SNP π
rho200 -0.07; dR2 0.90
pR 0.006; pD 0.004
TcMar-Tc2 SynDiv
rho200 0.003; dR2 0.28
pR 0.779; pD 0.130
TcMar-Tigger SNP π
rho200 -0.06; dR2 0.82
pR 0.002; pD 0.002
TcMar-Tigger SynDiv
rho200 -0.001; dR2 0.03
pR 0.106; pD 0.252
TcMar-Mariner SNP pi
rho200 -0.06; dR2 0.73
pR 0.002; pD 0.002
TcMar-Mariner SynDiv
rho200 0.003; dR2 0.44
pR 0.647; pD 0.168
hAT-Charlie SNP π
rho200 -0.05; dR2 0.57
pR 0.002; pD 0.002
hAT-Charlie SynDiv
rho200 0.001; dR2 0.08
pR 0.090; pD 0.066
hAT-Tip100 SNP π
rho200 -0.03; dR2 0.44
pR 0.002; pD 0.002
hAT-Tip100 SynDiv
rho200 0.01; dR2 0.18
pR 0.476; pD 0.144
hAT-Blackjack SNP π
rho200 -0.04; dR2 0.39
pR 0.002; pD 0.008
hAT-Blackjack SynDiv
rho200 -0.004; dR2 0.08
pR 0.825; pD 0.366
hAT SNP π
rho200 0.02; dR2 0.01
pR 0.116; pD 0.903
hAT SynDiv
rho200 0.02; dR2 0.53
pR 0.060; pD 0.356
hAT-Ac SNP π
rho200 -0.02; dR2 -0.00
pR 0.685; pD 0.841
hAT-Ac SynDiv
rho200 -0.02; dR2 0.07
pR 0.490; pD 0.567
centr SNP pi
rho200 0.01; dR2 0.66
pR 0.036; pD 0.010
centr SynDiv
rho200 -0.05; dR2 1.23
pR 0.214; pD 0.008
SVA SNP π
rho200 -0.28; dR2 1.01
pR 0.002; pD 0.002
SVA SynDiv
rho200 -0.14; dR2 1.13
pR 0.002; pD 0.002
Helitron SNP π
rho200 -0.07; dR2 2.44
pR 0.092; pD 0.012
Helitron SynDiv
rho200 -0.03; dR2 -0.02
pR 0.362; pD 0.675
undefined (multi-class) SNP π
rho200 -0.04; dR2 0.71
pR 0.002; pD 0.002
undefined (multi-class) SynDiv
rho200 -0.004; dR2 0.21
pR 0.260; pD 0.302
Residualized log₂(scaled methylation variability)
Log1p-scaled diversity score

**Supplementary Fig. 2. Methylation variability in relation to nucleotide and synteny diversity across RepeatMasker families.**

Covariate-adjusted relationships between residualized $\log_2$(meVar), where meVar denotes scaled methylation variability, and nucleotide diversity (SNP π) or synteny diversity (SynDiv), which quantifies variation in collinearity among homologous projections, across 30 RepeatMasker families containing 6,604,530 distinct 200-bp intervals. Grey points are reliability-weighted means in predictor-quantile bins containing at least 50 intervals. Red and blue curves show low-rank cubic regression splines for SNP π and SynDiv, respectively, fitted to the binned summaries. The *x* axis gives ln[1 + diversity/median(non-zero diversity)]. Labels report the 200-bp partial Spearman correlation (rho200), incremental weighted $R^2$ (dR2; Δweighted $R^2 \times 10^{-3}$), raw two-sided 5-Mb block-bootstrap *P* value for the partial correlation (pR) and raw two-sided block-bootstrap *P* value for Δweighted $R^2$ (pD). The RepeatMasker-family panel labelled centr for SynDiv has only three distinct predictor values and is therefore shown as binned points without a fitted curve; absence of the curve does not indicate absence of an association.

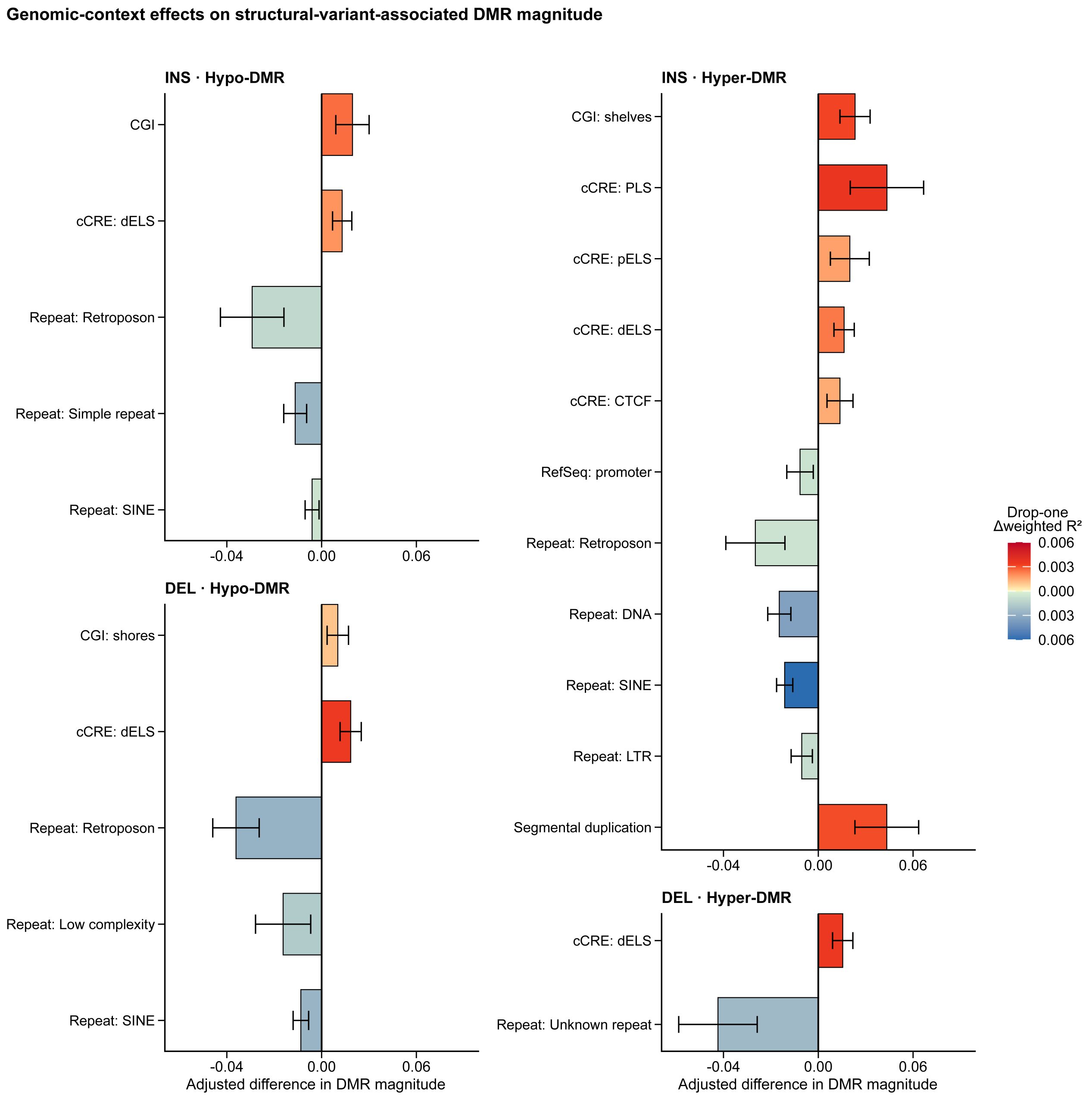


**Supplementary Fig. 3. Genomic-context effects on structural-variant-associated differentially methylated region (DMR) magnitude.**

Directional weighted linear models relating the local genomic context of a DMR to DMR magnitude, defined as the absolute carrier-minus-non-carrier methylation difference for each structural variant (SV)-DMR pair and weighted by the number of CpGs in the DMR. Panels are arranged with insertions (INS) above deletions (DEL), and hypomethylated DMRs to the left of hypermethylated DMRs. Bars give mutually adjusted coefficients for DMR-local genomic features in each SV-class and DMR-direction combination; positive and negative values indicate larger and smaller modelled DMR magnitude, respectively. The displayed terms include CpG-island (CGI), RepeatMasker, RefSeq and ENCODE candidate cis-regulatory element (cCRE) annotations. Horizontal lines are 95% confidence intervals based on SV-clustered heteroskedasticity-consistent type 1 (HC1) standard errors. Bar colour gives the signed drop-one Δweighted $R^2$ of the corresponding term. Terms entered the display when Benjamini–Hochberg $q < 0.05$ within an SV class and DMR direction.

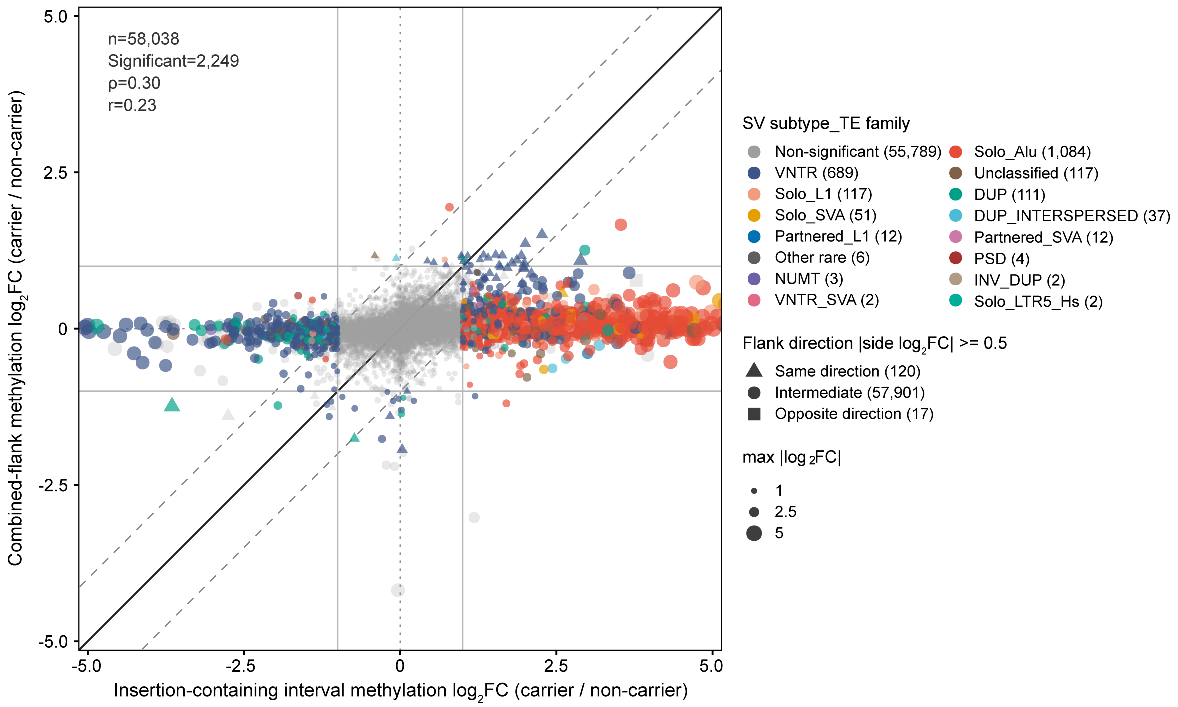


**Supplementary Fig. 4. Methylation changes in insertion-containing and flanking intervals.**

For insertions (INS), methylation $\log_2$ fold changes comparing carrier and non-carrier haplotypes at the projected 200-bp insertion-containing interval and across ten flanking 200-bp intervals for 58,038 insertions. The projected carrier interval includes CpGs within the inserted sequence. Flanks comprise offsets −6 to −2 and +2 to +6; the immediately adjacent intervals are excluded. Left and right flanks were tested separately by two-sided Wilcoxon rank-sum tests and combined by directional Stouffer meta-analysis. The axes show carrier-versus-non-carrier methylation $\log_2$ fold change at the insertion-containing interval and the weighted combined-flank effect. Of the tested insertions, 2,220 met Benjamini–Hochberg false-discovery rate ≤ 0.05 and $|\log_2$ fold change$| \geq 1$ at the insertion-containing interval, 48 met these thresholds in the flank meta-analysis and 19 met them in both, giving 2,249 insertions meeting the thresholds in either compartment. Coloured points denote significant structural-variant (SV) subtype-transposable-element (TE) family combinations; six singleton combinations are grouped as Other rare. Point shape denotes concordant, intermediate or opposing left- and right-flank effects, with concordance or opposition assigned when both side-specific $|\log_2$ fold changes$|$ were at least 0.5. Point size denotes the larger absolute $\log_2$ fold change across the insertion-containing interval and combined flanks. The solid diagonal denotes equal effects, dashed diagonals denote a one-$\log_2$-unit difference and horizontal and vertical reference lines mark $|\log_2$ fold change$| = 1$. Spearman $\rho = 0.297$; Pearson $r = 0.225$.

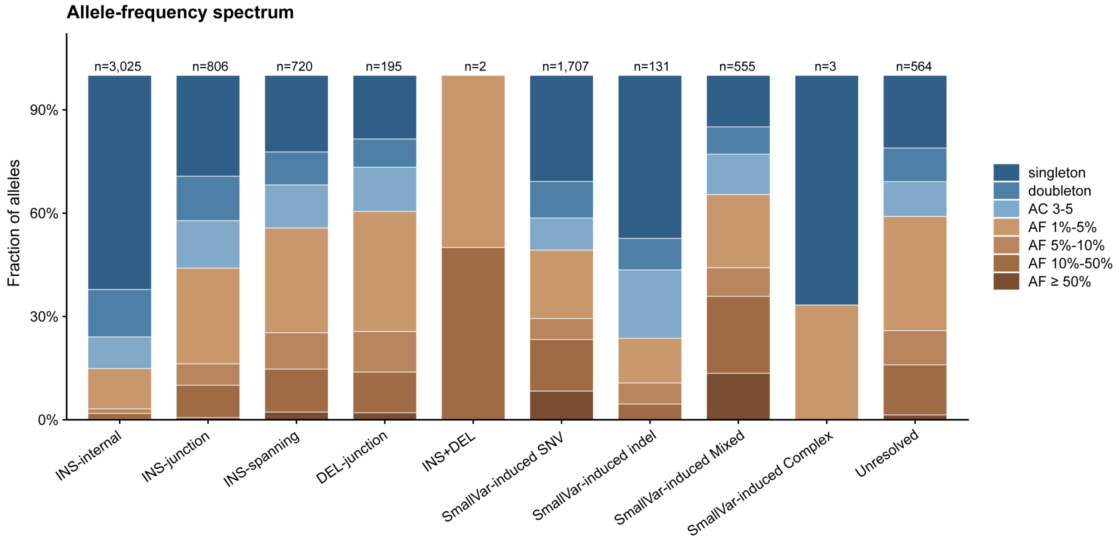


**Supplementary Fig. 5. Allele-frequency spectra of CpG-island polymorphism mechanisms.**

Within-class allele-frequency spectra for autosomal novel-locus CpG-island (CGI) alleles. Sequence-origin classes associated with insertions (INS) and deletions (DEL) are INS-internal, INS-junction, INS-spanning, DEL-junction and INS+DEL; small-variant-induced (SmallVar-induced) classes are single-nucleotide variant (SNV), short insertion/deletion (indel), mixed and complex. Unresolved alleles are shown separately. Each stacked bar shows the fraction of alleles in the indicated allele-count (AC) or allele-frequency (AF) bin. Values above bars give the total number of alleles in each sequence-origin class.

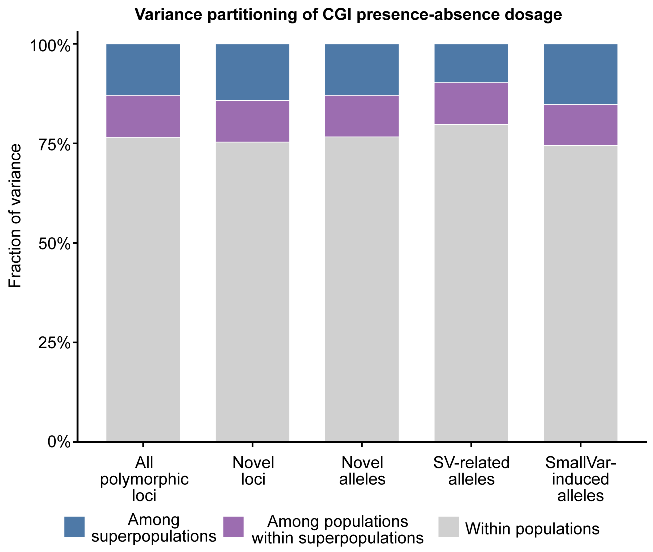


**Supplementary Fig. 6. Population-level variance partitioning of CpG-island presence-absence dosage.**

Variance in CpG-island (CGI) presence-absence dosage partitioned among five superpopulations, among populations within superpopulations, and within populations for 231 individuals. The five bars represent 12,605 polymorphic loci, 5,216 novel loci, 7,695 novel alleles, 4,747 structural-variant (SV)-related alleles and 2,386 small-variant-induced (SmallVar-induced) alleles. The within-population component accounted for 74.4–79.7% of total variance across the five sets; the remaining 20.3–25.6% occurred between population groups.

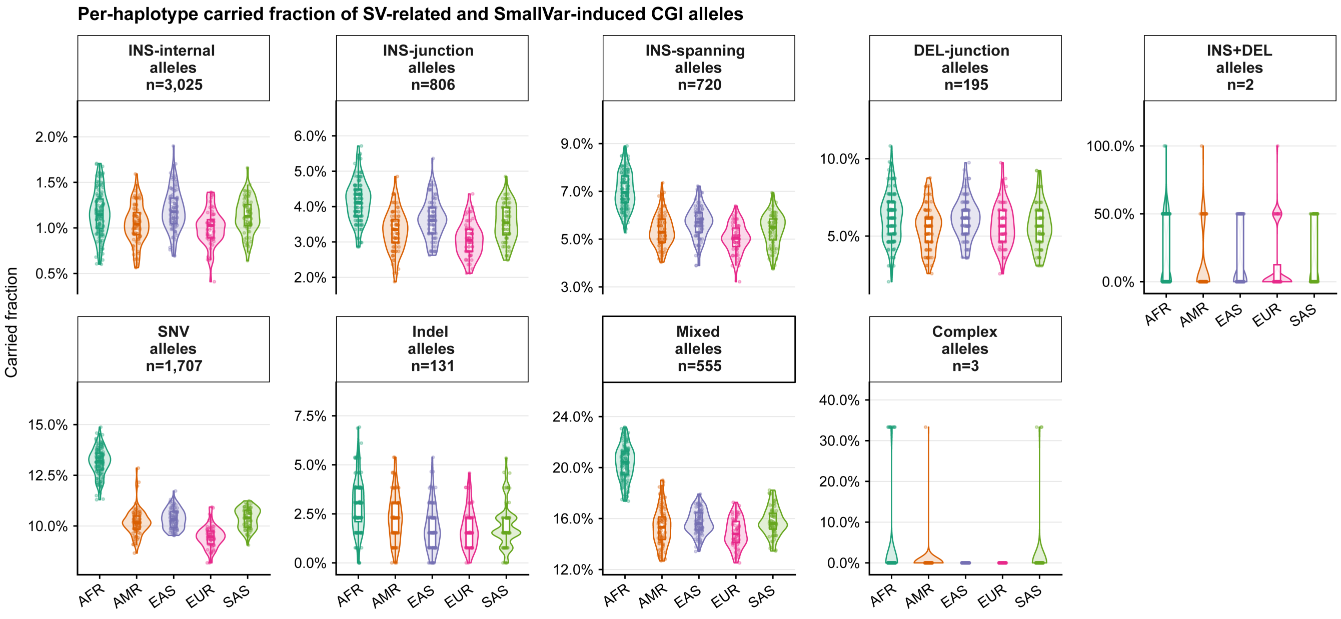


**Supplementary Fig. 7. Per-haplotype carried fraction of structural-variant-related and small-variant-induced CpG-island alleles.**

Per-haplotype carried fractions of autosomal novel CpG-island (CGI) alleles in nine structural-variant (SV)-related and small-variant-induced (SmallVar-induced) sequence-origin classes. Colours denote African (AFR), Admixed American (AMR), East Asian (EAS), European (EUR) and South Asian (SAS) superpopulations. Each point denotes one of 462 phased haplotypes, and the two haplotypes from an individual remain separate observations. The carried fraction is the proportion of callable alleles in the indicated category that are present on that haplotype. Violins show the distributions; boxes show medians and interquartile ranges, and whiskers extend to 1.5 times the interquartile range.

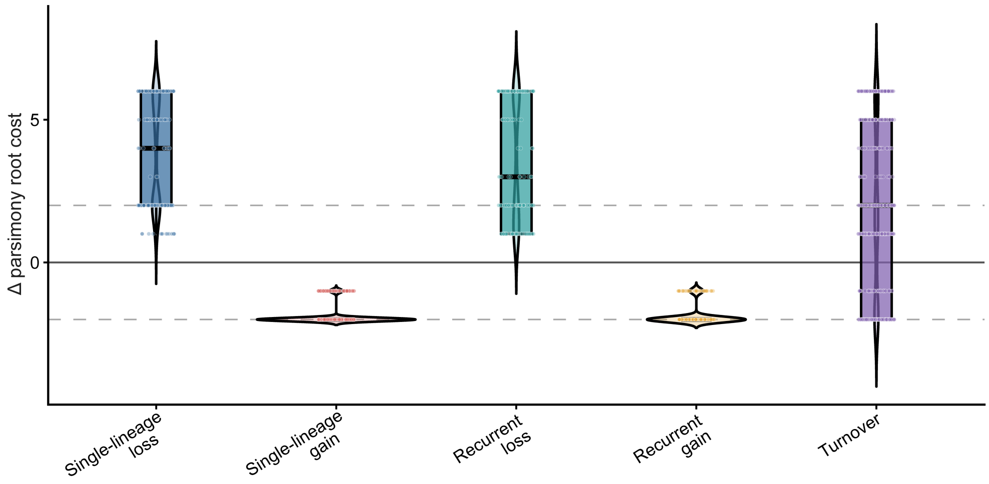


**Supplementary Fig. 8. Parsimony support for genealogy-based CpG-island root states.**

Difference in weighted-parsimony cost between CpG-island (CGI)-negative and CGI-positive root assignments across the five biological trajectory classes. The plotted quantity is the parsimony-cost difference *C*(root negative) − *C*(root positive), where *C* denotes the weighted-parsimony cost; positive values favour a CGI-positive root and negative values favour a CGI-negative root. The solid horizontal line marks equal cost, and dashed lines at +2 and −2 mark the supported-preference threshold. Loci with a cost difference of +1 or −1 retained the lower-cost root assignment but did not meet the supported-preference threshold; exact cost ties were excluded from the five biological classes. Violins show locus distributions; internal boxes show medians and interquartile ranges. Classes are single-lineage loss, single-lineage gain, recurrent loss, recurrent gain and turnover.

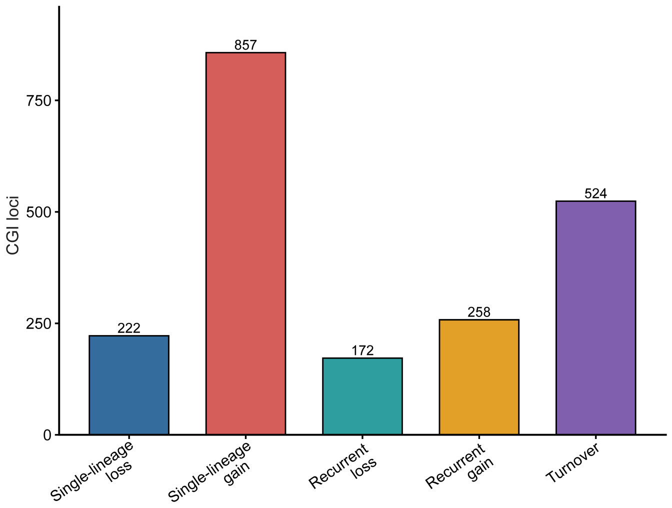


**Supplementary Fig. 9. Counts of CpG-island trajectory classes.**

The five displayed biological trajectory classes comprise 222 single-lineage losses, 857 single-lineage gains, 172 recurrent losses, 258 recurrent gains and 524 turnover loci among 2,279 candidate loci. Among root-resolved loci, single-lineage loss or gain was assigned when haplotypes carrying the changed state were compatible with a single clade under the prespecified clade-support criterion. Otherwise, recurrent loss required at least two losses in every optimal reconstruction and no permitted gain; recurrent gain was defined symmetrically. Remaining direction-resolved loci were classified as turnover. A further 132 loci had an exact root-state cost tie, one required no human-internal transition and 113 lacked a usable local genealogy; these loci were excluded from the five biological trajectory classes.

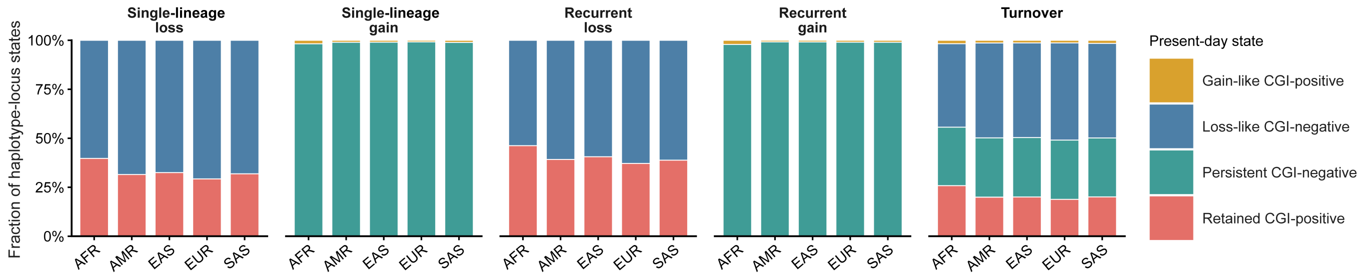


**Supplementary Fig. 10. Root-conditioned present-day CpG-island state composition across trajectories and superpopulations.**

Genealogy-root-conditioned present-day CpG-island (CGI) composition for 2,033 loci assigned to the five biological trajectory classes, stratified by superpopulation. Superpopulations are African (AFR), Admixed American (AMR), East Asian (EAS), European (EUR) and South Asian (SAS). Stacked bars show pooled fractions of retained CGI-positive, loss-like CGI-negative, gain-like CGI-positive and persistent CGI-negative haplotype-locus states. At loci with an inferred CGI-positive ancestor, present-day states were interpreted as retained or loss-like; at loci with an inferred CGI-negative ancestor, they were interpreted as gain-like or persistent. Turnover loci were interpreted according to their locus-specific inferred ancestral state before aggregation. The plot contains 933,151 called haplotype-locus states.

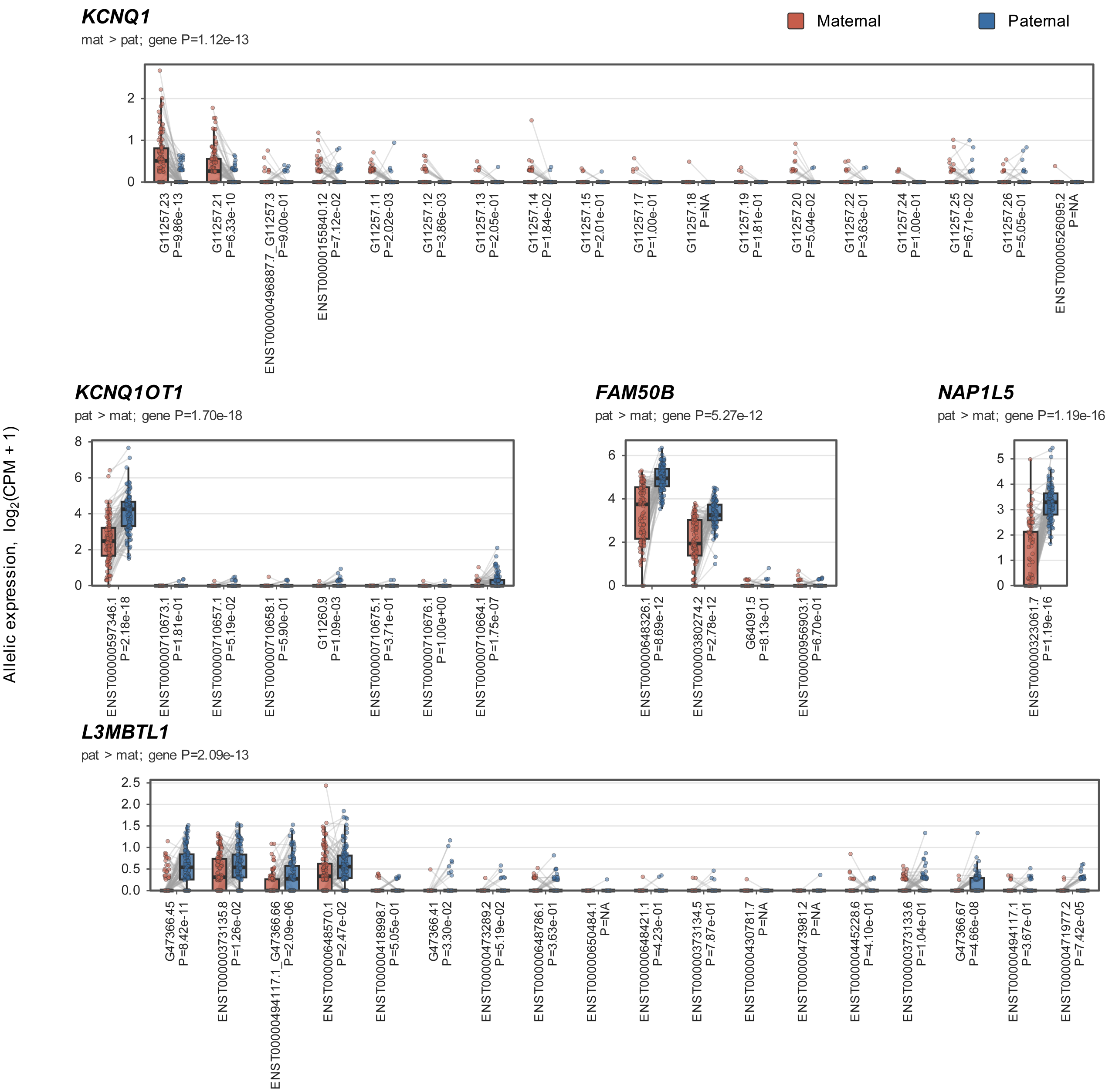


**Supplementary Fig. 11. Imprinted expression supports parental assignment in phased long-read RNA.**

Allele-specific expression at five established imprinted loci in 107 individuals with paired maternal and paternal long-read RNA measurements. Points show $\log_2$(counts per million (CPM) + 1), and grey lines join the maternal and paternal measurements from the same individual. Boxes show medians and interquartile ranges; whiskers extend to 1.5 times the interquartile range. Transcript labels give the complete transcript identifier and two-sided paired Wilcoxon signed-rank *P* value. Gene-level tests showed maternal enrichment for *KCNQ1* and paternal enrichment for *KCNQ1OT1*, *NAP1L5*, *FAM50B* and *L3MBTL1*.

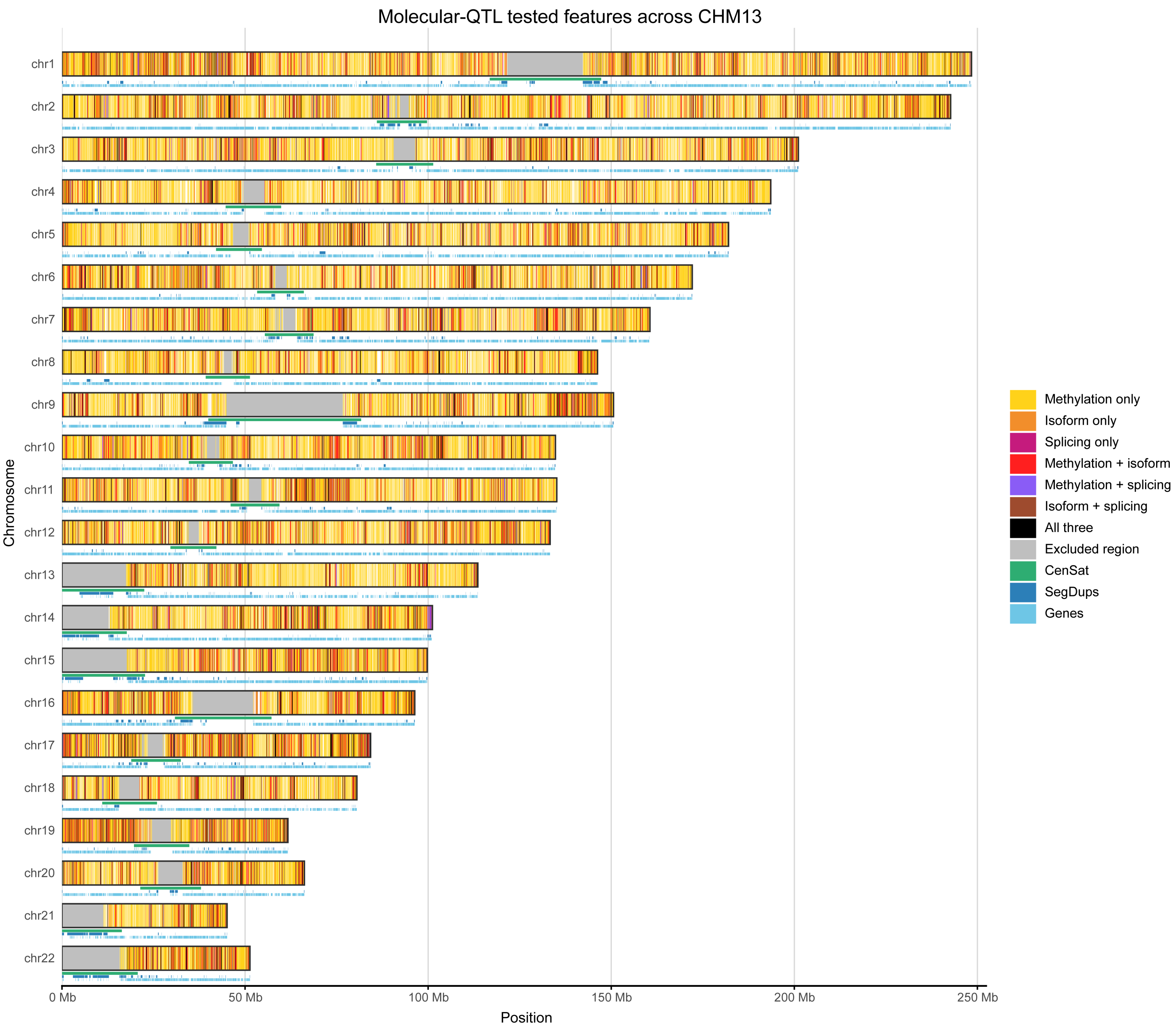


**Supplementary Fig. 12. Genomic distribution of molecular features tested for quantitative trait locus (QTL) mapping.**

Genomic distribution of molecular features tested for methylation, isoform and splicing QTL mapping on the CHM13 autosomes. Colours distinguish intervals covered by tested methylation intervals, isoforms or splice junctions alone, by each pair of feature classes or by all three. Testing included 9,623,107 projected methylation intervals, 66,951 isoforms from 12,726 genes and 49,221 splice junctions from 13,419 genes. Lower tracks show excluded regions, the

CHM13 centromere/satellite repeat annotation (CenSat), segmental duplications (SegDups) and genes.

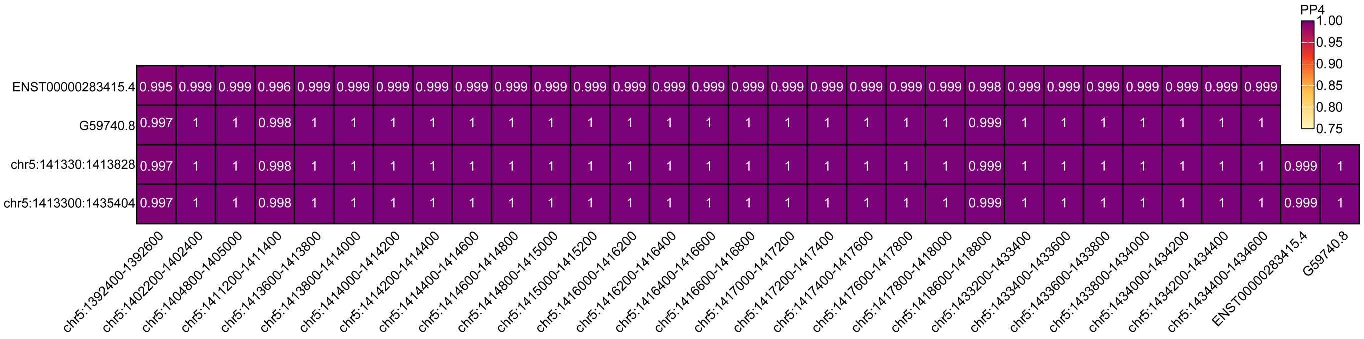


**Supplementary Fig. 13. Colocalization of methylation, isoform and splicing quantitative trait loci at *LPCAT1*.**

Pairwise coloc analysis at *LPCAT1* across 29 methylation intervals, two isoforms and two splice junctions. Heatmap colour and cell labels show the posterior probability that each pair of molecular features shares one causal signal (PP4). Median PP4 values were 0.9988 across 58 isoform-methylation pairs, approximately 1.0 across 58 splicing-methylation pairs and 0.9994 across four isoform-splicing pairs; the corresponding minimum values were 0.9946, 0.9974 and 0.9988. rs36978 was the leading shared variant across the methylation-isoform, methylation-splicing and isoform-splicing comparisons.

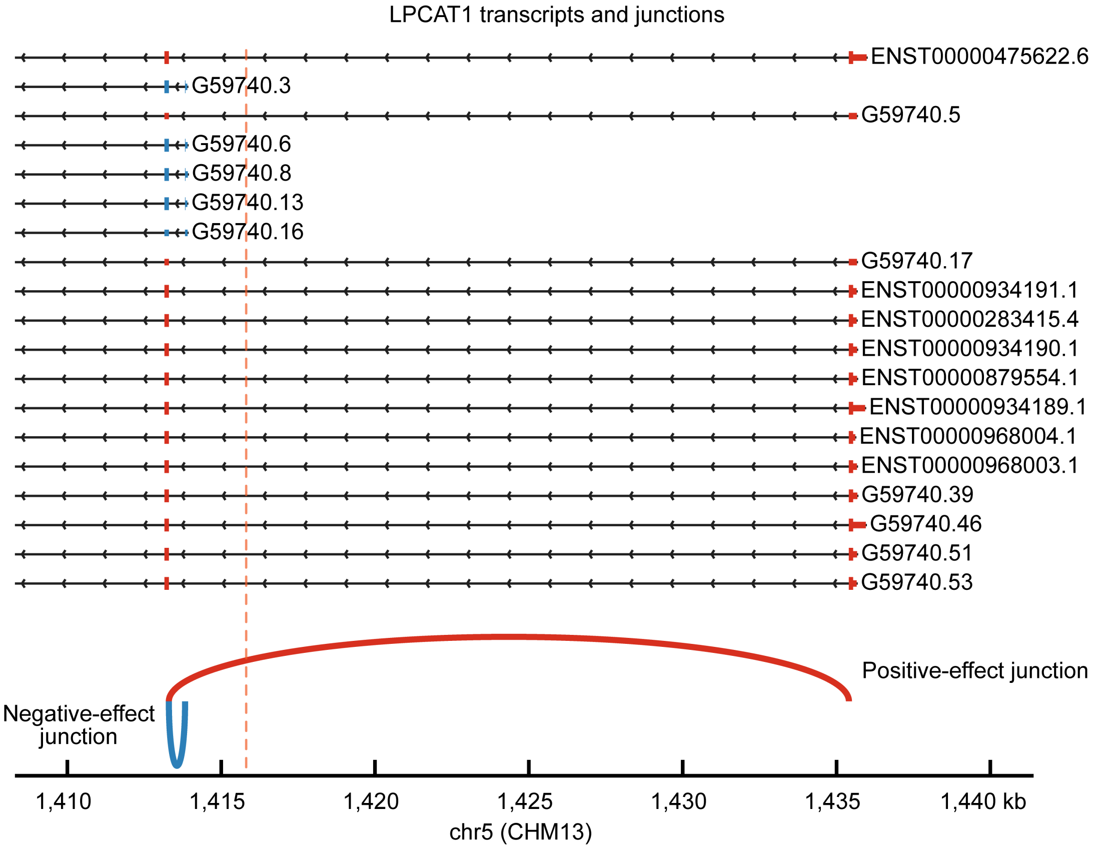


**Supplementary Fig. 14. Transcript structures containing the colocalized *LPCAT1* splice junctions.**

Nineteen *LPCAT1* transcript models containing the two colocalized splice junctions: eight reference-matched and 11 novel models from the HPRC2 transcript catalogue generated in this study, comprising five novel-not-in-catalogue, two novel-in-catalogue and four incomplete-splice-match isoforms. Blue and red arcs mark the junctions with usage effects of −0.387 and 0.333, respectively, relative to the rs36978 T allele, resolving reciprocal splice choices across the displayed transcript structures. The red vertical dashed line marks the position of the shared leading variant, rs36978. Models are from the long-read transcript catalogue and were quantified on CHM13.

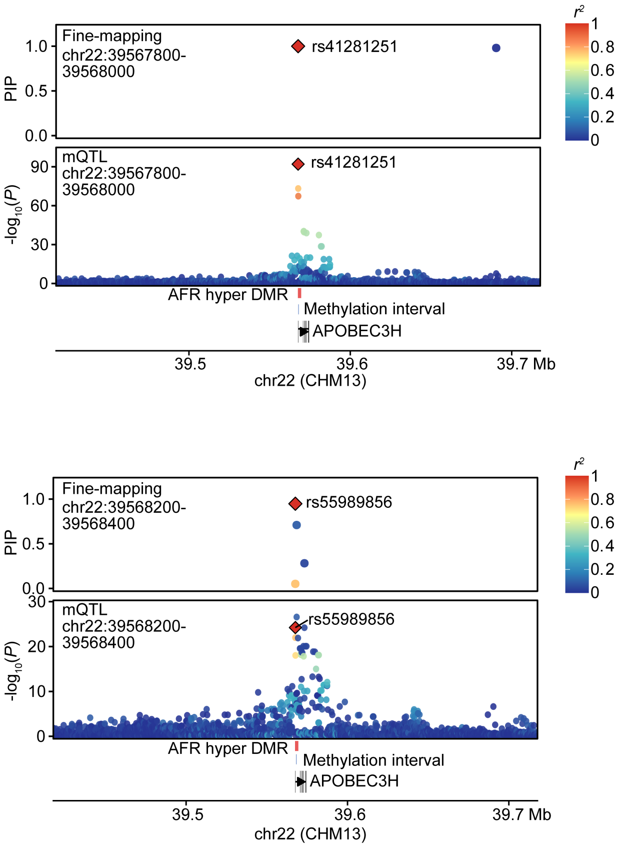


**Supplementary Fig. 15. Fine-mapped methylation quantitative trait loci (mQTLs) at *APOBEC3H*.**

Fine-mapping of two *APOBEC3H* methylation intervals. Red diamonds mark the highlighted fine-mapped variants, and point colour gives phased-haplotype $r^2$ to the highlighted variant in each panel. rs41281251 had a posterior inclusion probability (PIP) of 1.0 for chr22:39,567,800–39,568,000 ($\beta = 0.703$; $P = 9.17 \times 10^{-93}$), was the nominally most significant variant for the interval and was enriched in African (AFR) haplotypes (allele frequency, 0.629 versus 0.094 in non-AFR haplotypes). rs55989856 had PIP = 0.948 for chr22:39,568,200–39,568,400 ($\beta = 0.382$; $P = 5.92 \times 10^{-25}$) and was enriched in AFR (0.700 versus 0.100). It was not the nominally most significant variant for that interval. Both positive mQTL effects were concordant with hypermethylation of the differentially methylated region (DMR) at the *APOBEC3H* promoter in AFR.

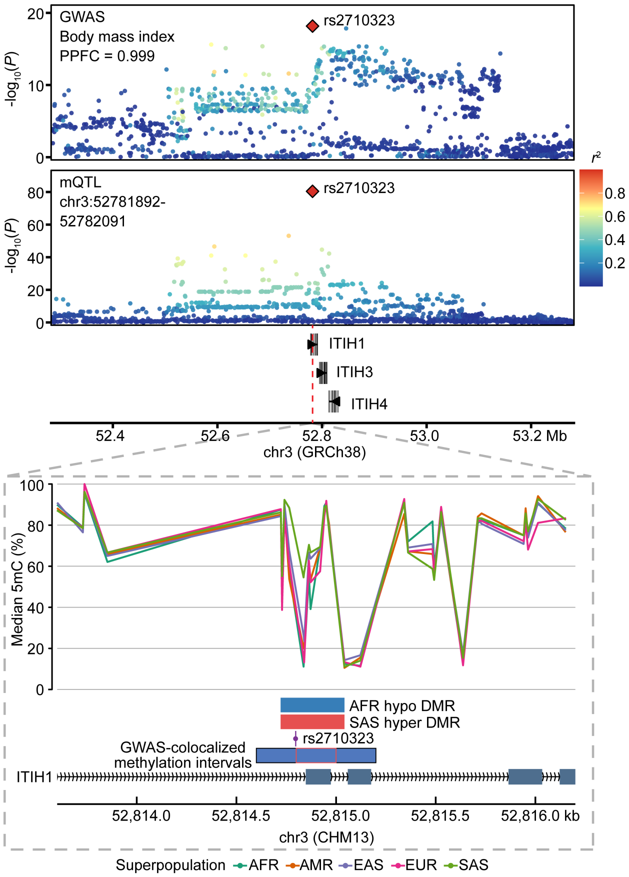


**Supplementary Fig. 16. Body mass index colocalization with a methylation quantitative trait locus (mQTL) at the ITIH locus.**

Body mass index association at rs2710323 colocalizes with three methylation intervals across the *ITIH1-ITIH3-ITIH4* locus (HyPrColoc posterior probability of full colocalization (PPFC) = 0.999). The association panel uses GRCh38, whereas the lower genomic track uses CHM13. Blue boxes in the lower CHM13 track mark the colocalized methylation intervals and the red outline marks the representative interval plotted above. The representative mQTL has $\beta = -0.362$ ($P = 3.97 \times 10^{-81}$) relative to the CHM13 alternate allele. The effect allele is most frequent in African (AFR; 0.743) and least frequent in South Asian (SAS; 0.236) haplotypes. The allele-frequency and methylation directions were concordant with both the AFR-hypomethylated and SAS-hypermethylated differentially methylated regions (DMRs). The effect allele was enriched in AFR (0.743 versus 0.472 in non-AFR haplotypes; 1.574-fold) and depleted in SAS (0.236 versus 0.613 in non-SAS haplotypes; 2.598-fold). A filled red diamond marks the HyPrColoc candidate variant and point colour gives phased-haplotype $r^2$ to it. rs2710323 is the genome-wide association study (GWAS) lead; gene labels provide genomic context and do not assign targets.
